# Nucleotide and structural polymorphisms of the eastern oyster genome paint a mosaic of divergence, selection, and human impacts

**DOI:** 10.1101/2022.08.29.505629

**Authors:** Jonathan B. Puritz, Honggang Zhao, Ximing Guo, Matthew P. Hare, Yan He, Jerome LaPeyre, Katie E. Lotterhos, Kathryn Markey Lundgren, Tejashree Modak, Dina Proestou, Paul Rawson, Jose Antonio Fernandez Robledo, K. Bodie Weedop, Erin Witkop, Marta Gomez-Chiarri

## Abstract

The eastern oyster, *Crassostrea virginica*, is a valuable fishery and aquaculture species that provides critical services as an ecosystem engineer. Oysters have a life-history that promotes high genetic diversity and gene flow while also occupying a wide range of habitats in variable coastal environments from the southern Gulf of Mexico to the southern waters of Atlantic Canada. To understand the interplay of genetic diversity, gene flow, and intense environmental selection, we used whole genome re-sequencing data from 90 individuals across the eastern United States and Gulf of Mexico, plus 5 selectively bred lines. Our data confirmed a large phylogeographic break between oyster populations in the Gulf of Mexico and the Atlantic coast of the USA. We also demonstrated that domestication has artificially admixed genetic material between the two ocean basins, and selected lines with admixed ancestry continue to maintain heterozygosity at these sites through several generations post admixture, possibly indicating relevance to desirable aquaculture traits. We found that genetic and structural variation are high in both wild and selected populations, but we also demonstrated that, when controlling for domestication admixture across ocean basins, wild populations do have significantly higher levels of nucleotide diversity and copy number variation than selected lines. Within the Atlantic coast, we detected subtle but distinct population structure, introgression of selected lines within wild individuals, an interaction between structural variation and putatively adaptive population structure, and evidence of candidate genes responding to selection from salinity. Our study highlights the potential for applying whole genome sequencing to highly polymorphic species and provides a road map for future work examining the genome variation of eastern oyster populations.

## Introduction

Aquaculture, one of the fastest growing food production sectors with more than 500 diverse species in culture, offers tremendous opportunities to meet increased human nutritional demands (Houston et al. 2020). Unlike traditional terrestrial crop and livestock species whose domestication began over 12,000 years ago and has resulted in significant genetic, genomic, and phenotypic alterations that benefit mankind (Diamond 2002), most aquatic animals reared in captivity for food production are relatively few generations removed from natural populations (Teletchea and Fontaine 2014). They have not been subject to artificially manipulated evolutionary processes (genetic drift, inbreeding, domestication selection, and targeted selective sweeps) and consequential changes in phenotypic and genetic diversity to the same extent as their terrestrial counterparts ((Jensen 2014; Bouwman et al. 2018);(Dutta et al. 2020). Most aquatic species are not considered domesticated or profoundly changed from their wild ancestors (Hedgecock 2012); however, many are undergoing artificial selection for the genetic improvement of commercially important traits.

Many marine fish and shellfish species have external fertilization, very high fecundity and high early life history mortality that tend to be associated with increased genetic diversity and high levels of polymorphism (Williams 1975). Williams argued that very high fecundity - millions of eggs from single females - enhances the capacity for genetically selective deaths each generation and therefore allows for rapid response to temporal or spatial variation in selection pressures. His theoretical conjectures are still relatively unexplored in natural populations (Marshall et al. 2010)(Rey et al. 2020), but are consistent with rapid genetically-based gains of 5-15% per generation realized in multiple fish and shellfish selective breeding programs (Hedgecock 2011)(Olesen et al. 2003). This exceptionally high fecundity and genetic diversity, coupled with emerging genomic resources (Yáñez, Newman, and Houston 2015), enables simultaneous domestication and selection in aquaculture species within decades compared to terrestrial animals and plants (Larson and Fuller 2014).

Eastern oysters, *Crassostrea virginica*, native estuarine cup oysters that support some wild fisheries and a rapidly growing aquaculture sector in the western North Atlantic, are broadly distributed from New Brunswick, Canada, to Yucatan, Mexico. Several genetically divergent regional subpopulations have been described (Buroker 1983; Reeb and Avise 1990; Karl and Avise 1992; Hoover and Gaffney 2005; Varney et al. 2009) and experimental tests have provided support for local adaptation across latitudes (Bushek and Allen 1996; Dittman 1997; Burford et al. 2014). Eastern oysters are highly fecund, protandrous hermaphrodites with external fertilization and a planktonic larval stage (Thompson, Guo, and Harrison 1996). Early life stages experience high mortality (Osman, Whitlatch, and Zajac 1989; Morgan 1995), but those that survive settle on or near reefs composed of conspecifics (Atwood and Grizzle 2020) and reefs with greater genetic diversity (Hanley et al. 2016). These life history characteristics, in a species occupying a spatially and temporally heterogeneous environment, support the potential for strong within-generation selection causing differentiation among wild adults at all spatial scales, but uniquely at distances below average larval dispersal such as within estuaries (Rose, Paynter, and Hare 2006). This same evolutionary capacity may also contribute to the success of selective breeding for commercially valuable traits that increase food production.

Eastern oyster breeding efforts generally targeted increased growth and survival in response to natural disease epizootics (Gjedrem and Baranski 2009; Davis and Barber 1999; Calvo et al. 2003; Yu and Guo 2004; Frank-Lawale, Allen, and Dégremont 2014; Casas et al. 2017; Leonhardt et al. 2017; Guo 2021). The initial eastern oyster selection lines were created after an intense natural selection event in the early 1960’s in Delaware Bay involving infection with the haplosporidian parasite that causes the disease known as MSX (Haskin and Ford 1979). Performance evaluations of MSX survivors and their offspring were the first to demonstrate population-level acquisition of heritable disease resistance traits, and they evolved resistance within a few generations (Ford and Haskin 1987). Since then, a handful of eastern oyster selected lines have been developed at regional scales for research and industry use including UMFS, NEH, DEBY, LOLA, and OBOYS2 (Table 2, see also supplementary data for a more detailed history of selective breeding in this species). The approach to and degree of selection in this species varies by geographic region, but historically has relied on mass selection, whereby individuals with desired trait values are crossed in successive generations (Gjedrem and Baranski 2009; Guo 2021). However, environmental heterogeneity with respect to temperature, salinity, and biological stressors (e.g. naturally occurring pathogens) as well as demonstrated performance advantages at the site of selection, necessitates genetic improvement at or below the regional scale (Rose, Paynter, and Hare 2006; Proestou et al. 2016). These regional selective breeding programs would benefit from improved genomic resources, facilitating knowledge on regional differences in genome architecture and population structure, including assessment of the degree of within-population diversity vs among-population diversity contributing to the realized products of selective breeding.

We resequenced whole genomes from 90 eastern oysters, including both wild populations (naturally recruiting) and artificial selection lines, to investigate population structure attributable to demographic history and selection across multiple spatial scales (species range, the eastern United States, within estuaries). Within-population genetic and structural diversity is assessed relative to population/line history, and we explore the interplay between selection, divergence, and genomic architecture across all lines and populations, and across only wild populations from the Atlantic. Lastly, we perform two pairwise contrasts of wild populations that differ in salinity looking for candidate genes that may be responding to selection.

## Methods

### Sample collection and environmental data

Ninety adult eastern oysters were collected in the fall of 2017 from four water bodies in the United States: one in the Northeast (Damariscotta River, Maine), two in the Mid-Atlantic region (Delaware Bay and Chesapeake Bay), and the fourth in the northern Gulf of Mexico (Louisiana). Within each water body, two wild populations from sites with putatively divergent salinity regimes were sampled. In addition, at least one selected population commercially grown in each estuary was sampled (Figure with map of populations, Table 1). A total of thirteen localities/lines were evaluated, 8 wild and 5 selected. A detailed description of the history of the selected lines is provided in the Supplementary Text.

**Table 1.** Wild individual collection localities and environment

| Population abbreviation | Location | N | Lat | Long | Salinity | Local Farming history? |
| --- | --- | --- | --- | --- | --- | --- |
| HI | Hog Island, Damariscotta River, ME | 6 | 44.0133 | -69.541689 | High | Yes |
| SM | Sherman Marsh, Sheepscot River, ME | 6 | 44.0141558 | -69.5949957 | No data | No |
| HC | Hope Creek, Delaware Bay, NJ | 6 | 39.447222 | -75.519444 | Low | No |
| CS | Cape Shore, Delaware Bay, NJ | 6 | 39.074218 | -74.913879 | Med | Yes |
| CLP | Chlora's Point, Chesapeake Bay, MD | 6 | 38.6270278 | -76.135671 | Low | No |
| HC_VA | Hummock Cove, Chesapeake Bay, VA | 6 | 37.617443 | -75.663169 | High | No |
| SL | Sister Lake, LA | 6 | 29.249167 | -90.92111 | Low | Unknown |
| CL | Calcasieu Lake, LA | 6 | 29.815556 | -93.348889 | Med | Unknown |

Mean annual salinity "High" for high salinity (>24ppt), "Med" = med salinity(15-24ppt), "Low" = low salinity(<15ppt)

Temperature and salinity data for each site where oysters were collected and/or selected were obtained through various data sources including on-site data loggers from the oyster collector and nearby NERR, NOAA, or USGS environmental data buoys. For most sites, data were collected at 15-30 minute intervals year round. Mean annual temperature (degrees Celsius) and salinity (parts per thousand) were calculated using data collected for three or more years leading up to this study.

### Whole Genome Sequencing

#### Sequencing

Oyster genomic DNA was extracted from mantle tissue of six individuals per population using a modified CTAB protocol (Winnepenninckx, Backeljau, and De Wachter 1993). Purified genomic DNA from each individual was used to prepare paired-end libraries (Lucigen Shotgun library preparation, PCR free) and sequenced on an Illumina HiSeq X PE 150 bp platform to 10-20X coverage at the Centre d’ Expertise et de Services, Genome Quebec. Twelve samples were included in sequencing and variant calling from known inbred experimental lines and populations as part of a different research project. They were removed after variant calling and not used for any population level analyses.

#### Nucleotide Variant Calling

All bioinformatic code, scripts, and supplemental files can be found in the GitHub repository (https://github.com/The-Eastern-Oyster-Genome-Project/2022_Eastern_Oyster_Population_Genomics) An RMarkdown document is provided to fully reproduce all bioinformatic and population genomic analysis. Raw sequencing reads were processed with a modified version of the dDocent pipeline (Puritz, Hollenbeck, and Gold 2014). Briefly, reads were trimmed for low quality bases and adapter sequences using the program fastp (Chen et al. 2018). Trimmed reads were then mapped to the haplotig-masked version of the eastern oyster genome (Puritz et al. 2022) using bwa (Li and Durbin 2010) with modified mismatch and gap-opening parameters (-B 3 -O 5) to help compensate for the high levels of polymorphism in the oyster sequences. Duplicates were marked using Picard (Institue 2019), and subsequent BAM files were filtered with samtools (Li et al. 2009) to remove low quality mappings, secondary alignments, and pcr duplicates. Samples were then genotyped for small nucleotide variants (SNPs, InDels, small complex events) using freebayes (Erik Garrison and Marth 2012).

Raw variants were filtered in parallel using a combination of bcftools (Danecek et al. 2021) and vcftools (Danecek et al. 2011). In short, variants were first filtered based on allelic balance at heterozygous loci (between 0.1 and 0.9) and quality to depth ratio of greater than 0.1. Variants were further filtered based on mean-depth, excluding all loci above the 95th percentile. Variants were then decomposed into SNPs, and InDels using vcflib (E. Garrison 2016). SNPs were further filtered to allow for no missing data and only biallelic SNPS, and then variants separated into two sets of variants, one with a minor allele frequency (MAF) of 1% and the other with a MAF of 5%.

SNPs were also subset to create a dataset exclusively for wild Atlantic populations (SM, HC, CS, CLP, HC_VA) referred to as the WILDAE subset hereafter. MAF was recalculated for these datasets and filtering reapplied. For analysis sensitive to linkage between loci, we used the “snp_autoSVD’’ function of bigsnpr (Privé et al. 2018) with the settings (min.mac =4,size=10) to clump SNPs based on physical distance, minor allele frequency, and linkage and to remove outliers, hereafter referred to as “LD-pruned SNPs”.

#### Structural variant calling

Structural variants (SVs), including deletions, insertions, duplications and inversions, account for a significant fraction of many genomes, including humans. To identify candidate SVs, we used the short-read sequence data with the reference genome. Structural variants were called on unfiltered BAM files using the program Delly (Rausch et al. 2012) following the “germline sv calling” (https://github.com/dellytools/delly#germline-sv-calling). SVs were then filtered using Delly with the “germline” filter. BCFtools (Danecek et al. 2021) was used to convert inversion SVs to a bed file and then switch “LowQual” genotypes to missing and SVs were filtered to a subset with no missing data. Using this filtered SV subset, read based copy number (RDCN) for insertions, deletions, and duplications were extracted to a tab delimited list.

### Population Structure at the Nucleotide Level

#### Linkage decay and relatedness

We assessed the decay of linkage disequilibrium on each chromosome using vcftools for the LD calculations. Across all LD decay analyses, we used eight different base pair window sizes for pairwise calculation of the squared correlation coefficient between genotypes (*r*^*2*^) (50, 200-250, 450-500, 4500-5000, 9500-10000, 49500-50000, 99500-100000, 499500-500000). The mean *r*^*2*^ for each window size was summarized for each chromosome. We analyzed two specific subsets, the first being (1) Atlantic wild populations (WILDAE) and individuals from (2) Atlantic selection lines (UMFS, NEH, DEB, LOLA), with one of a pair of highly related individuals excluded. We used vcftools (Danecek et al. 2011) --relatedness option to calculate the Ajk statistic using the method of Yang et al, Nature Genetics 2010 (doi:10.1038/ng.608), and excluded one of a pair of individuals with Ajk > 0.5 for subsequent analyses.

#### PCA

To explore broad patterns of population structure in 78 individuals across the 13 groups, we converted a compressed VCF file to Plink format using Plink2 (Shaun Purcell et al. 2007; S. Purcell and Chang, n.d.) and imported into the bigsnpr (Privé et al. 2018) package within the R computing environment (Team and Others 2013). PCA analysis was using the “LD-pruned SNPs. PCA analysis was also performed on subsets of individuals constituting all wild collection localities (WILD) and all wild Atlantic coast (WILDAE).

#### Pairwise population differentiation

To investigate the genomic differentiation among oyster populations, global mean *F*_ST_ for each population contrast was estimated by averaging across non-overlapping windows using popgenWindows.py (https://github.com/simonhmartin/genomics_general), with 150 SNPs per window. Pairwise global *F*_ST_ contrasts were visualized using ComplexHeatmap (Gu, Eils, and Schlesner 2016).

#### Global *F*_*ST*_ and outlier detection

Global measures of population structure were calculated from biallelic SNPs with no missing data allowed across all individuals. The program OUTFlank (Whitlock and Lotterhos 2015) was used to calculate *F*_*ST*_ using all SNPs. Outliers were inferred relative to a null *F*_*ST*_ distribution based on trimmed SNP datasets with heterozygosity greater than 0.1 and using a random set of 50,000 SNPs sampled from the LD-pruned set of SNPs (Privé et al. 2018). Outlier detection was repeated with the WILD and WILDAE locality subsets. Within locality sample size was only six for this study, so a conservative FDR of 5 × 10^−4^ was used for selecting outlier loci.

#### Admixture analysis

Putative neutral SNP sets were identified after removing candidate adaptive loci (outliers) and accounting for linkage disequilibrium (10Kb clumping interval, across all samples). The Bayesian clustering program STRUCTURE v2.3.4 (Pritchard, Stephens, and Donnelly 2000) was then used to characterize population structure and admixture using three neutral datasets: all populations, Atlantic populations, and populations from the Gulf of Mexico. We randomly selected 10K SNPs from each dataset for STRUCTURE analysis. The admixture model with correlated allele frequencies was applied with a burn-in of 10,000 iterations followed by 100,000 Markov Chain Monte Carlo (MCMC) repetitions. We used different numbers of assumed population genetic clusters (K=1-10) to determine the best-supported K values using the program KFinder (Wang, 2019), repeated 10 times for each K. Three different criteria in KFinder: Pr[X|K] (Pritchard et al., 2000), ΔK (Evanno, Regnaut, and Goudet 2005), and Parsimony Index (Wang 2019) were estimated for each dataset. Among them, the Parsimony Index selects the K that repetitively returns the minimal mean admixture of individuals and Wang (2019) shows that this approach has improved performance over ther two former estimations under a variety of population structure (e.g. low population differentiation, hierarchical structure) and sampling scenarios.

### Comparison of genomic diversity among wild and selected individuals

#### Nucleotide diversity

Nucleotide diversity (*π*) was calculated within each line and locality, calculated per site and across 10 kilobase windows of the genome using VCFtools (Danecek et al. 2011). The *π* for selected lines was calculated as an average of values from the DEBY, LOLA, NEH, OBOYS, and UMFS samples and the *π* for wild localities was calculated as an average across CL, CLP, CS, HC, HC_VA, HI, SL, and SM. Differences between wild and selected nucleotide diversity were tested globally using a t-test pooling all windows from wild and all windows from selected groups. Additionally, *π* was recalculated for wild localities from the Atlantic coast (CLP, CS, HC, HC_VA, HI, and SM) and for selected lines originating from only Atlantic wild individuals (NEH, UMFS) to help control for the selected lines DEBY and LOLA which have a known history of admixture between both ocean basins.

#### Heterozygosity

Per-site values of observed heterozygosity were calculated with vcflib (E. Garrison 2016) within each line and locality. Per-site values were averaged across 10 kilobase windows using the program BEDtools (Quinlan and Hall 2010), and mean values for wild and selected lines were determined by averaging values in the identical manner as was performed for nucleotide diversity. Differences between wild and selected heterozygosity were tested globally using a t-test across all windows.

#### Tajima’s *D*

Tajima’s *D* was calculated within each line and locality, across 10 kilobase windows of the genome using the software VCF-kit (Cook and Andersen 2017). Values for wild and selected lines were determined by averaging values in the identical manner as was performed for nucleotide diversity. Differences between wild and selected Tajima’s *D* were tested globally using a t-test across all windows.

#### Copy number variation

Read based copy number (RDCN) for CNV loci were read into the adegenet R package (Jombart et al. 2008) as a haploid marker. Loci were filtered for a minor allele frequency greater than 5%. Allelic richness (AR) and gene diversity (the probability of any two individuals having different CN; GD) were calculated for each group using the heirfstat R package (Goudet 2005). Both AR and GD values were then converted to bed files and then averaged across 10 kbp windows using bedtools (Quinlan and Hall 2010), and values for wild and selected lines were determined by averaging values in the identical manner as was performed for nucleotide diversity. Differences between wild and selected AR and GD were tested globally using a t-test pooling all windows from wild and all windows from selected groups. Additionally, both statistics were recalculated for wild localities from the Atlantic coast (CLP, CS, HC, HC_VA, HI, and SM) and for selected lines originating from only Atlantic wild individuals (NEH, UMFS) to help control for the selected lines DEBY and LOLA which have a known history of admixture between both ocean basins.

#### Assessing genomic architecture Large putative inversions

Preliminary results indicated the presence of at least two large putative inversions in the genome. To infer boundaries of these regions, we combined information from multiple sources. First, we used PCAdapt (Luu, Bazin, and Blum 2017) on the full SNP data set (without thinning for LD) to calculate a PCA and associated p-values for each individual SNP as an outlier. Without thinning for LD, PCAdapt is known to be sensitive to areas of low-recombination and will group individuals into homozygous and heterozygous inversion arrangement clusters (Lotterhos 2019). We also compared differences in nucleotide divergence and heterozygosity between wild and selected lines, looking for continuous areas of pronounced differences. For all three data sets rolling means were calculated based on(1) PCAdapt p-values, (2) the wild and selected difference in heterozygosity and (3) the wild and selected difference in nucleotide diversity, across 1 megabase and 5 megabase windows. Large putative inversions were areas of the genome that were above the 90th percentile in at least four of the six different rolling means (three statistics X two different window sizes), indicating a strong signal across multiple measure and genomic intervals. After this first pass, three candidate intervals were expanded by relaxing the criteria to being above the 90th percentile in at least three of the six different rolling means. One detected inversion of chromosome 5 (61,560,000 to 80,150,000) intersects with a known misassembly located from approximately from 61,754,114 to 97,911,248 bp (Puritz et al. 2022), so for analysis, this inversion was split into two regions, one from 61,560,000 to 64,020,000 and a second from 64,030,000 to 80,150,000 which corresponds to a linkage group that is predominantly on chromosome 6.

#### Smaller putative inversions

Inversions were also called during structural variant genotyping with the software package DELLY ((Rausch et al. 2012); See Structural Variant Calling Section). Using this filtered SV subset, inversions were selected and converted to BED format using BEDtools (Quinlan and Hall 2010). Smaller inversions that intersected with the large putative inversion regions above were removed, leaving a curated set of small inversions.

### Examining differentiation across environments, domestication, and genomic architecture

#### Signals of differentiation across structural variation

To examine the role of structural variation in potentially shaping adaptive loci, we separately examined *F*_*ST*_ outliers within large putative inversion regions, small inverted regions detected by Delly, and those outliers that were outside of all known structuration variation. For the large and small inversion sets, we used BEDtools (Quinlan and Hall 2010) to intersect detected *F*_*ST*_ outliers with inverted regions. We visualized genotypes across samples at inversion-bound outlier loci using the GenotypePlot package (modified version 0.9; (Whiting et al. 2021). We also visualized variation at these loci with a PCA plot across the first two axes using the Adegenet R package (Jombart et al. 2008).

For outliers outside of structural variants, we investigated arge windows of elevated *F*_*ST*_ were identified by calculating rolling means of *F*_*ST*_ values calculated by OUTFlank (Whitlock and Lotterhos 2015) across 10, 50, and 100kb sliding by 10kb across the genome, and then selecting 10kb windows that were the top 95th percentile of all three categories. Elevated *F*_*ST*_ windows we again used BEDtools to remove any windows inside of any known structural variants. Genotypes of individual outlier SNPs (identified as described above) within these outlier regions were visualized with genotype plots in a similar fashion to outlier SNPs within large inversions. We also visualized variation at these loci with a PCA plot across the first two axes using the Adegenet R package (Jombart et al. 2008).

#### Identifying divergent copy number variants between populations and lines

Copy number variants (CNVs) were filtered for CNVs with a minor allele frequency greater than 5%. Differentiation at CNVs was calculated using the *V*_*ST*_ statistic (Redon et al. 2006) as implemented in (Steenwyk et al. 2016) and plotted across the genome. Divergent CNVs were identified by selecting CNVs within the 99.9 percentile *V*_*ST*_. CN variation was examined with a PCA plot across the first two axes using the Adegenet R package (Jombart et al. 2008) using all filtered CNVs and a subset of only the most divergent CNVs. Lastly, the copy number for the CNVs in the 99.9 percentile of *V*_*ST*_ for each individual was plotted in a heatmap.

### Signals of selection and differentiation across wild Atlantic oysters

Preliminary analysis showed that significant genomic structure, at the nucleotide and architectural level, was partitioned across samples originating from the two different coastlines (Gulf of Mexico and the Atlantic Coast of the USA). With only two wild localities sampled in the Gulf of Mexico, we focused subsequent analysis on samples across the six wild Atlantic localities.

#### Signals of differentiation across structural variation

Large windows of elevated *F*_*ST*_ were identified by calculating mean *F*_*ST*_ values calculated by OUTFlank (Whitlock and Lotterhos 2015) across 10 kilobase windows, and then selecting windows that were in the top 95th percentile. Outlier loci within these elevated windows of mean *F*_*ST*_ values were then partitioned into outliers within large inversion regions, within small inversions detected by DELLY (Rausch et al. 2012), and outlier SNPs outside of all detected structural variants. Genotypes of individual outlier SNPs (identified as described above) within these outlier regions were visualized with genotype plots in a similar fashion to outlier SNPs within large inversions and analyzed using principal components analysis in adegenet (Jombart et al. 2008).

#### *F*_*ST*_ genome scan using pairwise contrasts across salinity gradient contrasts in two Atlantic estuaries

To identify genomic signatures under selection, we conducted genome scans for specific population contrasts and examined the common shared outliers. We used VCFtools to exclude identified inversions and filter out loci with a minor allele frequency less than 0.05 (--maf 0.05) for each contrast VCF input (Danecek et al. 2011). A python script popgenWindows.py (https://github.com/simonhmartin/genomics_general) was used to compute Hudson *F*_*ST*_ in 1 Kb sliding windows with a 200 bp step size. Windows with fewer than 5 genotyped sites (-m 5) were ignored. The windowed *F*_*ST*_ was then Z-transferred (Z*F*_*ST*_) to normalize the resulting values (Pendleton et al. 2018). We identified the outlier windows in the upper 99.9th percentile of the windowed Z*F*_*ST*_ distribution in each contrast and merged the overlapping outliers with a customized R script. For those outliers that appeared to have functional relevance or shared overlapping coordinates by multiple contrast groups, we examined the heterozygosity and single SNP *F*_*ST*_ patterns inside and surrounding the outlier windows. For genomic ranges and individuals of interest we visualized genotypes with a clustered genotype plot (Whiting et al. 2020).

We extracted the gene annotation and gene ontology (GO) terms for outliers from an enhanced annotation of the genome (Proestou and Sullivan 2020; Sullivan and Proestou 2021). We then performed GO_MWU enrichment analyses based on Fisher’s exact test (Wright et al., 2015, https://github.com/z0on/GO_MWU), using binary measure (1 or 0, i.e., either identified as outlier windows or not) as the input. GO categories with a significant non-random distribution within the binary input, relative to genomic composition of annotations, were considered enriched and displayed in a hierarchical clustering tree with as.phylo function implemented in Ape (Paradis, Claude, and Strimmer 2004).

## RESULTS

### Whole Genome Sequencing

#### Sequencing

We generated over 3,558,207,970 read pairs for the entire project with an average of 39,535,644 +/-1,018,131 read pairs per sample. On average, 97.14% +/-2.49% of reads were retained after quality trimming and adapter removal, and 96.75% +/-2.48% of trimmed reads mapped to the genome with an average duplication level of 6.60% +/-0.17%. Total mapped reads covered 87.41% +/-0.23% of the reference genome with 1X coverage on average per sample. Post-filtration for duplicates, low-quality mapping, multi-mapping reads, and secondary alignments, left 72.90% +/-1.84% of total trimmed reads with high quality mapping on average per sample, and post-filtration mappings covered on average 80.45% +/-0.32% of the reference genome per sample. Per sample statistics for sequencing, read mapping, and percent of the genome covered can be found in Supplemental Table 2.

#### Nucleotide Variant Calling

After initial bioinformatic processing, 82,390,891 variants were called. After switching genotypes with less than five reads to missing, removing mitochondrial SNPs, and filtering by a call rate of 75%, 40,205,129 variants remained. Further filtering by allelic balance, mean depth, quality to depth ratio, redacted the dataset down to 29,917,303 variants. Freebayes (Erik Garrison and Marth 2012) is a haplotype-based genotyping software, so variant calls encompass SNPs, multiple nucleotide polymorphisms (MNPs), insertion/deletion polymorphisms (InDels), and small complex events (a combination of SNP, MNP, and InDel). The variant calls were decomposed into 13,323,451 short InDel variants and 52,971,541 SNPs. SNPs were subsequently filtered by call rate and minor allele frequency cutoff, and results for multiple combinations can be found in <u>Supplemental Table 3</u>. Population genomic analyses were performed using two primary data sets: both filtered to 100% call rate and biallelic SNPs only, one with a minor allele frequency of 1% (12,149,052 SNPs) and one with a minor allele frequency of 5% (5,574,080 SNPs). SNP datasets for the wild individuals (WILD) and wild individuals from localities along the Atlantic coast of the USA (WILDAE) subsets were also generated with a 5% minor allele frequency, with 5,092,795 and 4,482,328 SNPs respectively.

### Structural Variant Calling

After DELLY germline filtering, 279,389 total structural variants were called (216,912 deletions, 33,012 translocations, 15,347 duplications, 7,310 insertions, and 6,808 inversions). Deletions had a mean length of 823 +/-22 bp, duplications 14,784 +/-464 bp, insertions 48 +/-0 bp, and inversions had a mean length of 118,478 +/-2,857 bp. For copy number variant analysis (CNV), variants were further filtered to only deletion and duplication loci with no missing data, leaving 21,158 deletion variants (mean length of 354 +/-38 bp) and 1,611 duplication variants (mean length of 5996 +/-997 bp). See Modak et al. (2021) for a more thorough examination of structural variation in an earlier version of the eastern oyster genome.

### Population Structure

#### Linkage decay

In both the wild Atlantic samples and the selected lines, linkage disequilibrium decayed rapidly, reaching average levels below an *R*^*2*^ of 0.2 within 300 bp and below 0.1 by at least 5,000 bp (<u>Figure S3</u>). Atlantic wild samples showed a lower background level of genome-wide linkage disequilibrium, reaching an asymptotic *R*^*2*^ < 0.05 by 50,000 bp with selected samples reaching a near asymptotic *R*^*2*^ < 0.075 by 50,000. Wild samples also showed a higher variance of linkage disequilibrium among chromosomes than selected samples.

#### PCA

After pruning across 10kb windows and the removal of long-range outliers of linkage disequilibrium, 1,049,202 SNPs were used to evaluate population structure across all 13 resequenced eastern oyster populations; eight wild localities and five selected lines. The first PC axis (explaining 7.2% of the total variance) clearly differentiated Atlantic individuals from the Gulf of Mexico (GoM; = wild localities SL, CL and selected line OBOYS) (Fig. 1a). Two selected lines with known mixed ancestry between the Atlantic and GoM (DEBY and LOLA) were intermediate on PC1 with DEBY closer to the Atlantic localities and LOLA closer to the GoM populations. PC2 (3% variance explained) clearly distinguished the selected line NEH from all other localities and lines. PC3 and PC4 similarly distinguished the selected lines OBOYS, UMFS, and LOLA from other individuals (Supplemental Fig. 2).

**Figure 1.**
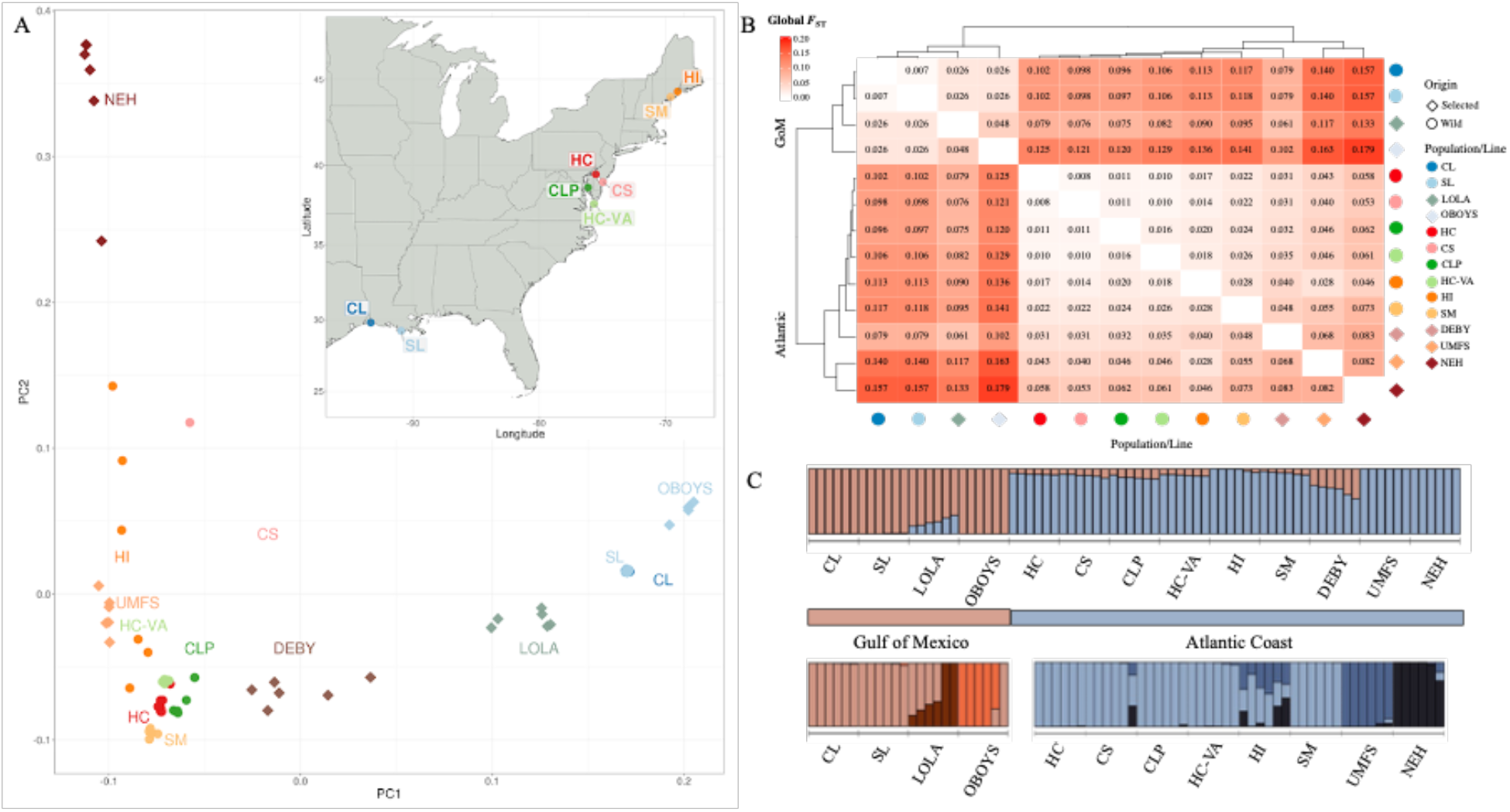
PCA of wild/Selected populations along with map of sampling locations. **Overall population structure of the eastern oyster in the USA**. Panel A is a PCA of wild populations and selected lines along with a map of wild oyster sampling locations. Individual points are color coded by sampling location or selected line and point shape indicates either a wild or selected origin as shown in legend of panel b. The top right panel B is pairwise global mean *F*_*ST*_ estimates for oyster populations and selected lines using all SNPs. Ordering of populations was based on clustering of the mean *F*_*ST*_ values as shown in marginal dendrograms. The bottom right panel C is STRUCTURE results across all populations (best K = 2) above separate analysis subsets within the Gulf of Mexico (K = 3) and the Atlantic Coast (K = 3) populations using 10K random neutral SNPs. The best K were identified based on most consistent K values across Pr[X|K], ΔK, and Parsimony Index methods. Source population colors do not correspond between the 3 analyses.

#### Pairwise population differentiation

Global mean *F*_*ST*_ in pairwise contrasts ranged from 0.0081 to 0.1798. For both wild populations and selected lines, the largest genomic *F*_*ST*_ pairwise contrasts were between Atlantic andGoM regions (mean among region contrasts 0.105 +/-0.008; Figure 1B). Within the Atlantic region, the *F*_*ST*_ among selected line contrasts were generally higher (0.068 - 0.133) than contrasts between wild populations (0.008 - 0.028) and wild populations versus selected lines (0.028 - 0.095). Of the two selected lines from the Chesapeake Bay known to be admixed between Atlantic and GoM broodstock (Table of selected lines), DEBY clustered with Atlantic and LOLA clustered with the GoM populations (Figure 1B).

#### Admixture Analysis

We examined the STRUCTURE outputs generated across all populations using Pr[X|K], ΔK, and Parsimony Index to assess number of clusters; all but the Pr[X|K] estimates had the strongest support for K = 2 (Figure 1c). The K=2 result reflects the Atlantic - Gulf divergence but with slight introgression of Gulf variation into some wild Atlantic populations and variable levels of admixture in DEBY and LOLA selected strain individuals. A separate analysis of Atlantic populations had the most support for K=3 with UMFS and NEH selected strains separated from wild populations (Fig. 1c). Selected lines in the GoM also were each distinct from wild populations (K=3; Fig. 1c)). Wild admixture with selected lines was detected at a low level in lower Delaware Bay (CS) and more extensively in one of the Maine samples (HI).

#### *F*_*ST*_ Outliers Identified

Comparing across all lines and localities, 24,269 SNPs (0.43% of the total SNPs) were identified as outliers with OUTFlank using an FDR of 5 × 10^−4^. For the WILD subset, a larger number of SNPs and percentage of SNPs were identified as outliers (34,523, 0.67%). Restricting analysis to the Atlantic WILDAE subset, an order of magnitude smaller raw number and percentage of SNPs were outliers (1,660, 0.03%). Detailed patterns for outlier subsets are presented below, and bed files of outlier SNPs can be found in the <u>github repository</u>.

### Genetic diversity among wild and selected individuals

Genome-wide estimates of nucleotide diversity (*π*) showed high levels of polymorphism across all wild localities and lines with an overall mean of 0.00534 using sliding 10 kb windows with SNPs above a 1% minor allele frequency (Figure 2). Overall, wild localities and selected lines had similar but significantly different levels of nucleotide diversity with wild samples having a mean *π* of 0.0054 +/-7.36e-06 and selected having a mean *π* of 0.0053 +/-9.13e-06 (t-test, wild > selected; t= 8.7939, df=401216.7, p = 7.2477e-19). The similar estimates of *π* from the overall dataset is potentially skewed by the inclusion of two selected lines with mixed ancestry from wild Atlantic and GoM individuals (DEBY and LOLA) which had estimates of 0.00560 +/-0.00002 and 0.00580 +/-0.00002, respectively. Removing them reduced the mean nucleotide diversity for selected lines to 0.0050 +/-1.13e-05. Furthermore, if the analysis is restricted to only wild Atlantic localities (HI, SM, HC, CS, CLP, HC-VA) and selected lines (NEH, UMFS) that originate entirely from Atlantic populations, and the dataset restricted to SNPs with a minor allele frequency of 1% within these individuals, the difference in nucleotide diversity becomes higher and statistically significant with wild samples having a mean *π* of 0.0043 +/ 7.33e-06 and selected having a mean *π* of 0.0038 +/-1.05e-05 (t-test, wild > selected;*t*=36.05, df=148453.5, *p* = 1.0047e-283). Observed heterozygosity, estimated from genome wide 10 kilobase intervals, was, on average, 0.246, across all samples (<u>Figure S4</u>). Among wild localities, the highest levels of heterozygosity were observed in oysters from the mid-Atlantic localities, with means ranging from 0.245-0.247 (see Table S4 for values, standard deviation, and standard error). Oysters from Northern Atlantic localities had slightly lower heterozygosity estimates than the mid-Atlantic (0.236-0.243). In contrast to estimates of nucleotide diversity, oysters from GoM localities had lower levels of heterozygosity than Atlantic. Across all selected lines of either Atlantic-only or GoM-only origins, heterozygosity ranged from 0.238 (NEH) to 0.252 (UMFS), and the two lines with mixed ancestry (DEBY and LOLA) had the highest overall estimates of 0.265 and 0.255 respectively.

**Figure 2.**
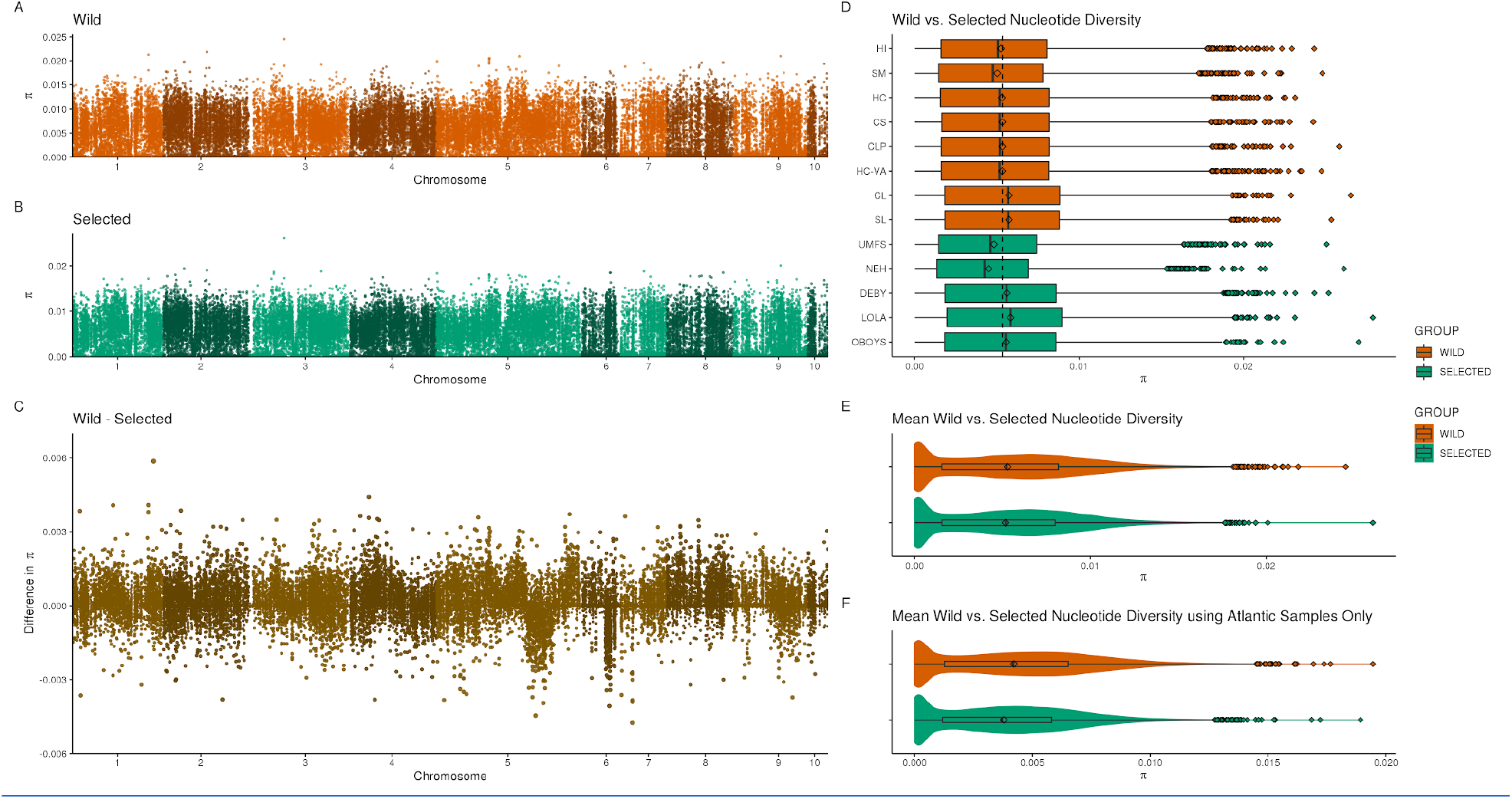
Comparison of nucleotide diversity across wild and selected samples. **Comparison of nucleotide diversity across wild populations and selected lines**. Panel (A) is values of π averaged across all wild populations in 10kb windows across the entire genome. Panel (B) panel is values of π averaged across all selected populations in 10kb windows across the entire genome. Panel (C) is the difference between wild and selected in 10kb windows across the entire genome. Panel (D) is a boxplot of 10kb averaged values broken down by population and line. Panel (E) is a violin and boxplot of 10kb averaged values averaged across groups. Panel (F) is identical to Panel (E), except it uses only loci that are variable within Atlantic selected and wild populations.

In contrast to nucleotide diversity, overall the selected lines had significantly higher levels of observed heterozygosity (t-test, selected > wild; t=-19.465, df= 74319, *p* = 1.78e-84) than wild populations, with selected lines having a mean of 0.251 +/-0.003 and wild localities a mean of 0.242 +/-0.003. The two selected lines with mixed ancestry from wild Atlantic and GoM individuals (DEBY and LOLA) were not primarily driving this pattern, as removing them only lowered the selected line heterozygosity mean to 0.245 +/-0.003. In fact, when restricting this analysis to only wild localities and selected lines that originate entirely from the Atlantic coastline (HI, SM, HC, CS, CLP, HC-VA, NEH, UMFS), and the dataset was restricted to SNPs with a minor allele frequency of 5% within these individuals, selected lines still showed (0.2920 +/-0.0004) significantly higher levels of observed heterozygosity (t-test, selected > wild; t=-9.124, df= 67495.2, *p* = 3.70e-20) than wild localities (0.2870 +/-0.0003).

Genome-wide estimates of Tajima’s *D* were significantly and substantially higher for selected lines than for wild localities (t-test, selected > wild; t=-120.4395, df= 368899.5, *p* = 0). Both wild localities and selected lines showed positive values of Tajima’s *D* with wild localities having a genome wide average of 0.2268 +/-0.0012 and selected lines having an average of 0.4722 +/-0.0016. Per locality and selected line averages ranged between 0.10 for CS and 0.62 for NEH (<u>Figure S5</u>).

Copy number variation allelic richness (number of different CNs present, AR) and diversity (chance of any two individuals having different CNs) were both influenced by the oceanic basin of origin. When looking across the entire dataset, CNV AR was nearly identical between wild (1.8444 +/-0.4288) and selected (1.8467 +/-0.4023) samples (t-test, selected > wild; t=0.3889, df= 19918.51, p = 0.6513) (<u>Figure S6</u>), and CNV diversity was actually significantly higher (t-test, selected < wild; t=-2.44599, df= 19940.54, p = 0.0072) in selected lines (0.3288 +/-0.1566) than wild localities (0.3236 +/-0.1483; <u>Figure S7</u>). A large number of CNVs were only variable within a single ocean basin, driving a lot of line/locality AR estimates closer to 1 and CNV diversity estimates closer to 0, and the selected lines overall mean was skewed by the inclusion of two selected lines with mixed ancestry. Removing the DEBY and LOLA lines lowered the selected line mean AR to 1.8158 +/-0.4688, leading to a significant difference (t-test, selected < wild; t=5.0116, df=19549.09, p = 5.443e-07); removing DEBY and LOLA also reduced selected line CNV diversity to 0.3190 +/-0.1719, making wild diversity now significantly larger (t-test, wild > selected; t=2.0370, df=19579.63, p = 0.0208). Further, restricting this analysis to only wild localities and selected lines that originate entirely from the Atlantic coastline (HI, SM,HC,CS,CLP,HC-VA,NEH,UMFS), and the dataset restricted to CNVs with a minor allele frequency of 5% within this data subset, the difference is confirmed, with wild localities (1.9052 +/-0.4043) showing significantly higher levels of AR (t-test, selected < wild; t=4.074544, df=17413.7, p = 2.315e-05) than selected localities (1.8773 +/-0.5176; <u>Figure S8</u>). However, with this same subset, CNV diversity is statistically indistinguishable (t-test, selected = wild; t=1.3517, df=17396.18, p = 0.1765) between wild localities (0.3469 +/-0.1483) and selected lines (0.3435 +/-0.1902; <u>Figure S9</u>).

### Putative inversions and population differentiation

#### Putative inversions underlie differentiation across wild populations and selected lines

##### Large inversions in the eastern oyster genome

Three large genomic inversions (>5 Mb) were detected across three different chromosomes in the eastern oyster. The first inversion was found on chromosome 1 (NC_035780.1) spanning at least 6.4 Mb. The second and largest inversion spanned =at least 18.59 Mb. Due to a misassembly in the genome this inversion mapped to two different assembled chromosomes (assembled chromosomes 5 and 6). The third inversion was found on chromosome 6 (NC_035785.1; 29,630,000 to 44,000,000) and spanned at least 14.3 Mb. In total, these large inversions represent 39.36 Mb, or 6.8% of the unmasked portion of the genome assembly.

##### Large inversions contained signals of ocean basin and admixture via domestication

Large inverted genomic regions were delineated by a combination of metrics, including PCAdapt p-values (using all loci), heterozygosity, and nucleotide diversity, and 1,514 outlier loci identified by *F*_*ST*_ outlier analysis were within the bound of the four large inversion regions. Visualization of outlier loci genotypes (Figure 3) showed that inversion genotypes were strongly associated with ocean basin. Wild samples from the GoM were largely fixed for one homozygous genotype, and wild samples from the Atlantic were largely fixed for the alternative homozygous genotype with very few detected heterozygous genotypes in wild individuals. However, the two selected lines with mixed basin ancestry, LOLA and DEBY, both showed a large number of heterozygous genotypes, usually with one individual containing nearly all heterozygous genotypes across outlier SNPs within a single chromosomal inversion. The mixed ancestry and largely heterozygous inversion genotypes for domesticated lines DEBY and LOLA are driving the pattern of reduced heterozygosity and nucleotide diversity in wild samples across inverted regions (Figure 3 and pattern can also be seen in Figure 2). Looking at inversion outlier SNPs variation across principle component space, the Atlantic and GoM distinction is evident on PC 1. A second grouping of Eastern Atlantic samples is present in PC 2, separating wild and selected samples north and south of Long Island Sound.

**Figure 3.**
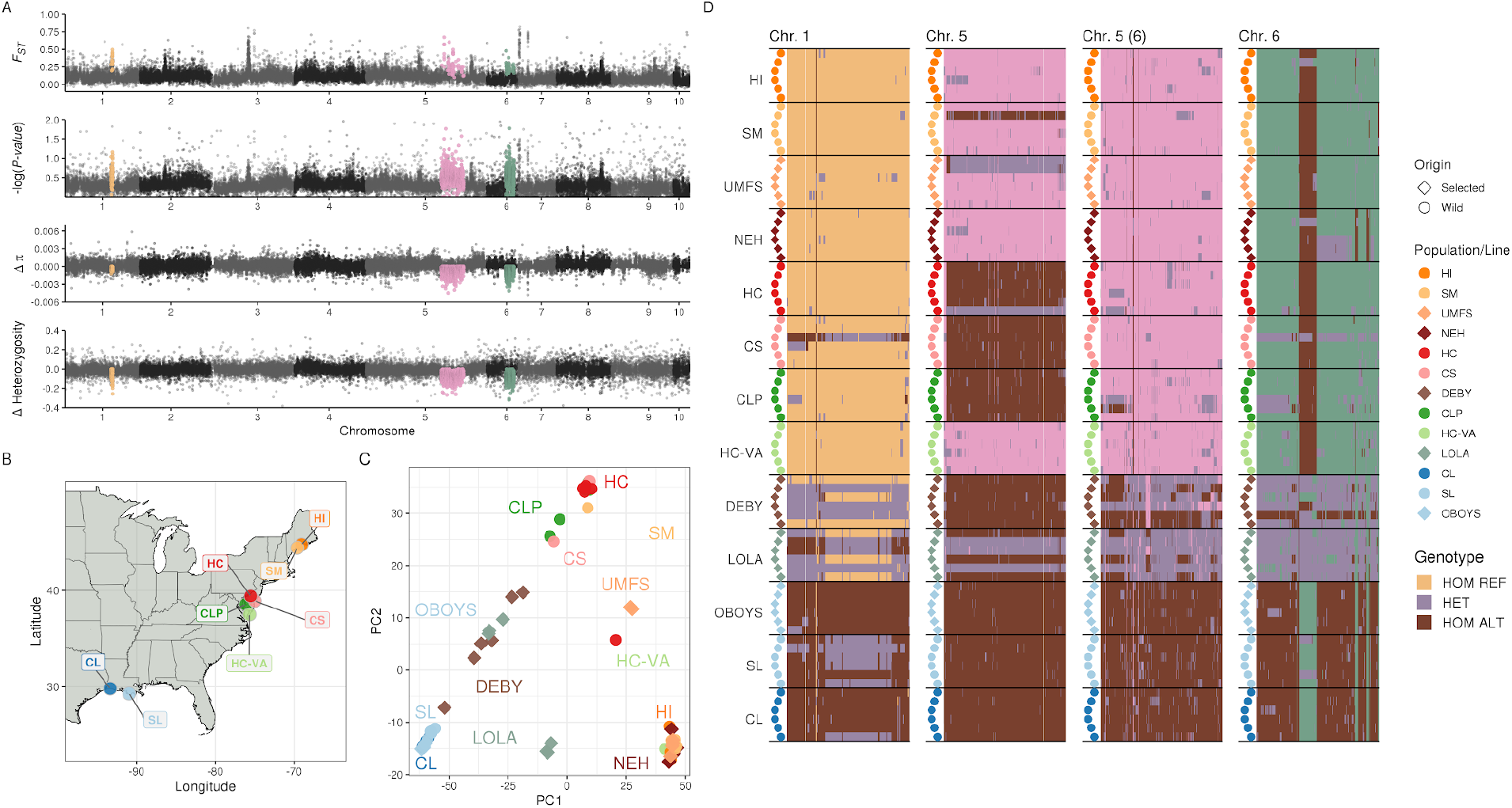
Outlier SNPs found within large, putative inversions. Panel (A)- Manhattan plots of various statistics across the eastern oyster genome. Top Manhattan plot-*F*_*ST*_ values calculated by OUTFLANK and averaged across 10kb windows. Second Manhattan plot - Negative log of P-values from PCAdapt using the full unthinned data set. For unthinned datasets, high levels of LD can lead to blocks of very low p-values which aided in the identification of large inversions. Third Manhattan plot-Mean difference in nucleotide diversity between wild and selected lines, averaged across 10kb windows. Fourth Manhattan plot-Mean difference in heterozygosity between wild and selected lines, averaged across 10 kb windows. For all Manhattan plots, outlier SNPs within putative large inversions are highlighted in color per chromosome (Chr 1-yellow; Chr 5 Pink; Chr 6 Green). Panel (B) is a map of all wild sampling locations. Pangel (C) is a PCA using all large inversion outliers. Panel (D) contains genotype plots for outlier SNPs found in the putative inversion regions. Homozygous reference allele genotypes are colored per chromosome identically to the manhattan plot. Due to a known misassembly, each half of the “pink” inversion maps to different regions of the assembly, even though they are found on the same chromosomal linkage group.

##### Small inversions showed signals of ancestry, admixture, and adaptation

Outlier loci within smaller genomic inversions (2,088) were identified via DELLY (Rausch et al. 2012) across six different chromosomes in the genome (<u>Figure S10</u>). Like for larger inversions, genotypes from loci on chromosome 3 (NC_035782.1) and chromosome 7 (NC_035786.1) were largely homozygous for the reference allele in the populations/lines with Atlantic origin and homozygous for the alternate allele in populations/lines with GoM origin. The exception being individuals from LOLA and DEBY, the two selected lines with mixed basin ancestry showing nearly entirely heterozygous genotypes. One individual from CS showed heterozygous genotypes at the same loci as well. Genotypes from loci on chromosomes 2 (NC_035781.1) and 8 (NC_035787.1) were also largely partitioned by ocean basin, but several individuals from Atlantic wild populations also had heterozygous genotypes, and genotypes from loci on chromosomes 4 (NC_035783.1) and 5 (NC_035784.1) showed a mosaic of the two previously described patterns.

##### Signals of selection outside of inversions

Outside of all detected genomic inversions, 3,194 outlier loci were detected and were largely fixed for the reference allele in localities/lines originating from the Atlantic and fixed for the alternative allele in localities/lines from the GoM (Figure 4). Some individuals from the selected lines DEBY and LOLA (the two lines with mixed basin ancestry), as well as an individual from CS (wild), showed largely heterozygous genotypes at outlier loci. Across chromosome 2 (NC_035781.1), chromosome 5 (NC_035784.1), and chromosome 8 (NC_035787.1) there were a number of Atlantic wild individuals that also had heterozygous genotypes at several loci, and a few homozygous for the alternate allele.

**Figure 4.**
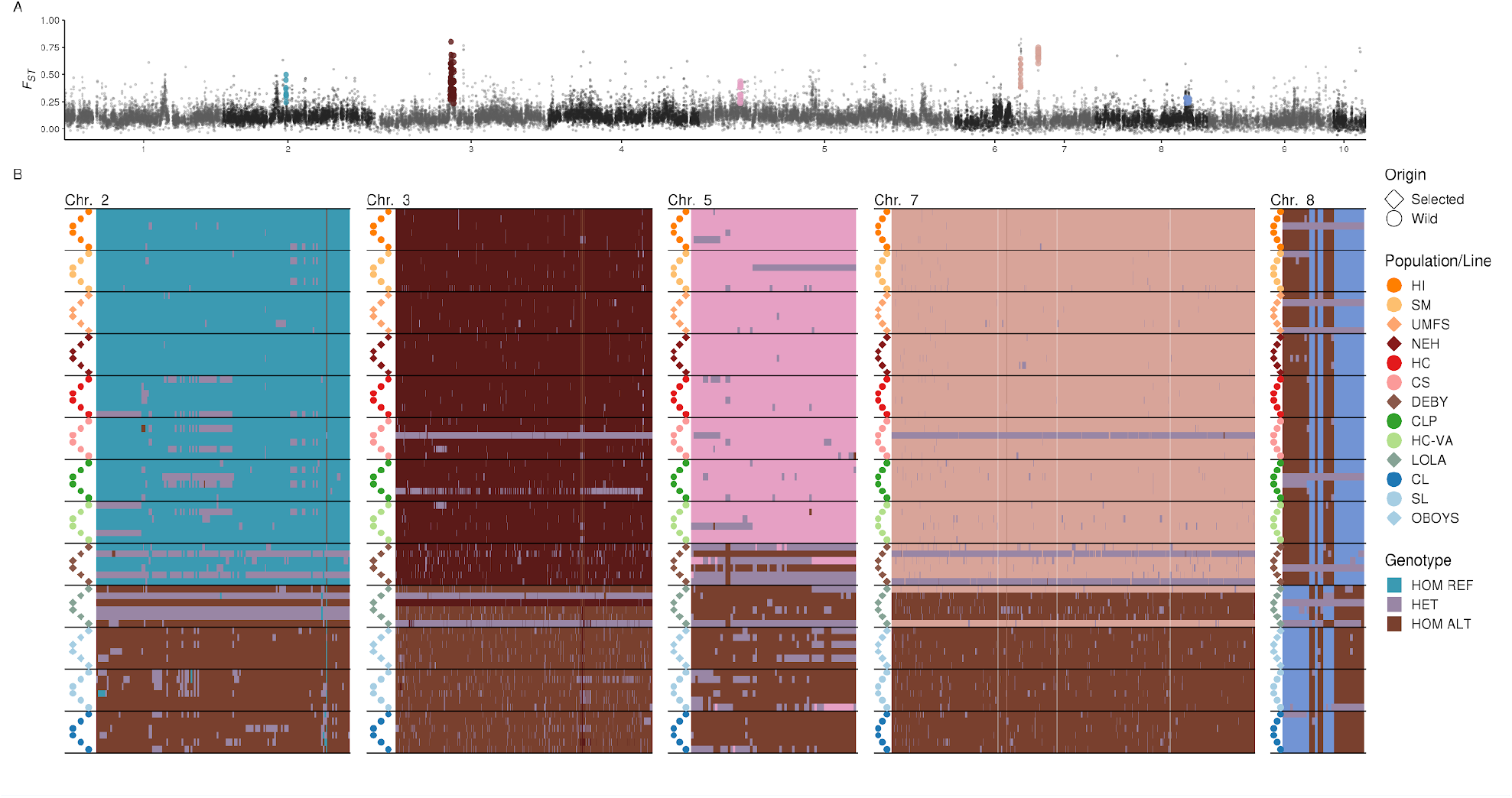
Outlier SNPs from genomic windows with large signals of elevated *F*_*ST*_ outside of identified inversions. **Genotype plots of outlier SNPs (outside of inversions)**. Panel (A)- Manhattan plot of *F*_*ST*_ values calculated by OUTFLANK for SNPs outside of any detected inversions, and averaged across 10kb windows. Examples of clustered-outlier SNPs are highlighted by different colors per chromosome (aqua Chr 2; maroon-Chr 3, orange-Chr 4, pink-chrm 5, peach-chrm 6, blue-chrm 8). Panel (B) contains genotype plots for outlier SNPs separated by chromosome and ordered vertically by sampling location/line. Homozygous reference allele genotypes are colored per chromosome identically to the manhattan plot (aqua Chr 2; maroon-Chr 3, orange-Chr 4, pink-chrm 5, peach-chrm 6, blue-chrm 8), then heterozgotes are uniformly purple and alternate homozygotes are brown.

##### Copy number variants show similar structure to SNPs

After filtering for minor allele frequency, 13,941 CNVs were retained for analysis. CNVs, like SNPs, were not evenly distributed across the genome with the first five chromosomes containing 11,258 (80.75%) of all CNVs. The mean *V*_*ST*_ across all loci was 0.07086 +/-0.00109, ranging from -0.16434 to 0.91913 and the most divergent CNVs in the 99.9th percentile had an average *V*_*ST*_ of 0.7209 +/-0.00005 (<u>Figure S11</u>). Principal Components Analysis (PCA) of all CNV loci showed a clear distinction of GoM and Atlantic samples across PC1 with individuals from the selected lines of LOLA and DEBY appearing between the two main clusters, indicative of their mixed ancestry. On PC2, the majority of wild individuals were clustered along the x-axis and a number of individuals from selected lines had negative PC2 values. PCA of the most divergent CNVs produced a similar pattern to all loci, except that PC2 only separated individuals from the mixed ancestry selected lines (DEBY, LOLA) and the GoM selected line (OBOYS). NEH and UMFS individuals mostly clustered with the wild Atlantic individuals. The most divergent CNVs seemed to largely be driven by loci with close to zero copies in GoM individuals and 2-3 copies for Atlantic individuals. There were three divergent CNVs that showed high copy numbers in GoM individuals. Overall, population structure was less clearly delineated with CNVs that it was with SNP loci.

##### Genomic architecture of outlier SNPs in wild Atlantic populations correlate with patterns of divergence, selection, and admixture

As a large portion of the identified genetic structure was associated with ocean basin, *F*_*ST*_ values were recalculated and outliers were inferred only for the wild Atlantic (WILDAE) individuals. OutFLANK identified 1,660 total outlier SNPs. We restricted this list of outliers further by only retaining SNPs that were located with 10kb windows that were above the 95th percentile of mean *F*_*ST*_. The reduced the outliers to 1,481 SNPs found across eight different chromosomes and across multiple levels of genomic architecture. 897 outlier SNPs were found within the two large inversion regions identified on chromosomes 5 (NC_035784.1) and 6 (NC_035785.1), 202 outlier loci were found within smaller inversions, and 364 outlier loci were found outside of known structural variants.

Principal Components Analysis of the outlier loci revealed differing patterns of population structure across genomic architecture, in contrast to a PCA using the LD-pruned dataset (Figure 5). For the thinned genomic SNPs (Figure 5B), there were three large clusters with a geographic pattern that matched closely to the whole data set PCA (Figure 1). In contrast, outlier SNPs had different patterns across each level of genomic architecture. Outlier SNPs within the two very large inverted regions on chromosomes 5 and 6 divided the wild individuals into two groups along PC1 with SM, HI, and HC-VA in one group and all others in the other (Figure 5D). The one exception was an HC individual that was positioned nearly on the origin between the two groups. Across PC2 SM, HI, and HC-VA were differentiated with the exception of one HI individual clustering with SM. For outliers in smaller inverted regions (Figure 5E), SM was differentiated from all others along PC1 and HC-VA (with the exception of one individual) differentiated from all others across PC2. Outlier SNPs found outside of structural variants (Figure 5F) differentiated SM individuals from all others along PC1 and separated HC-VA individuals along positive values of PC2 and all HI individuals (and one CS) along negative values. HI individuals also tailed towards SM individuals slightly on PC1.

**Figure 5.**
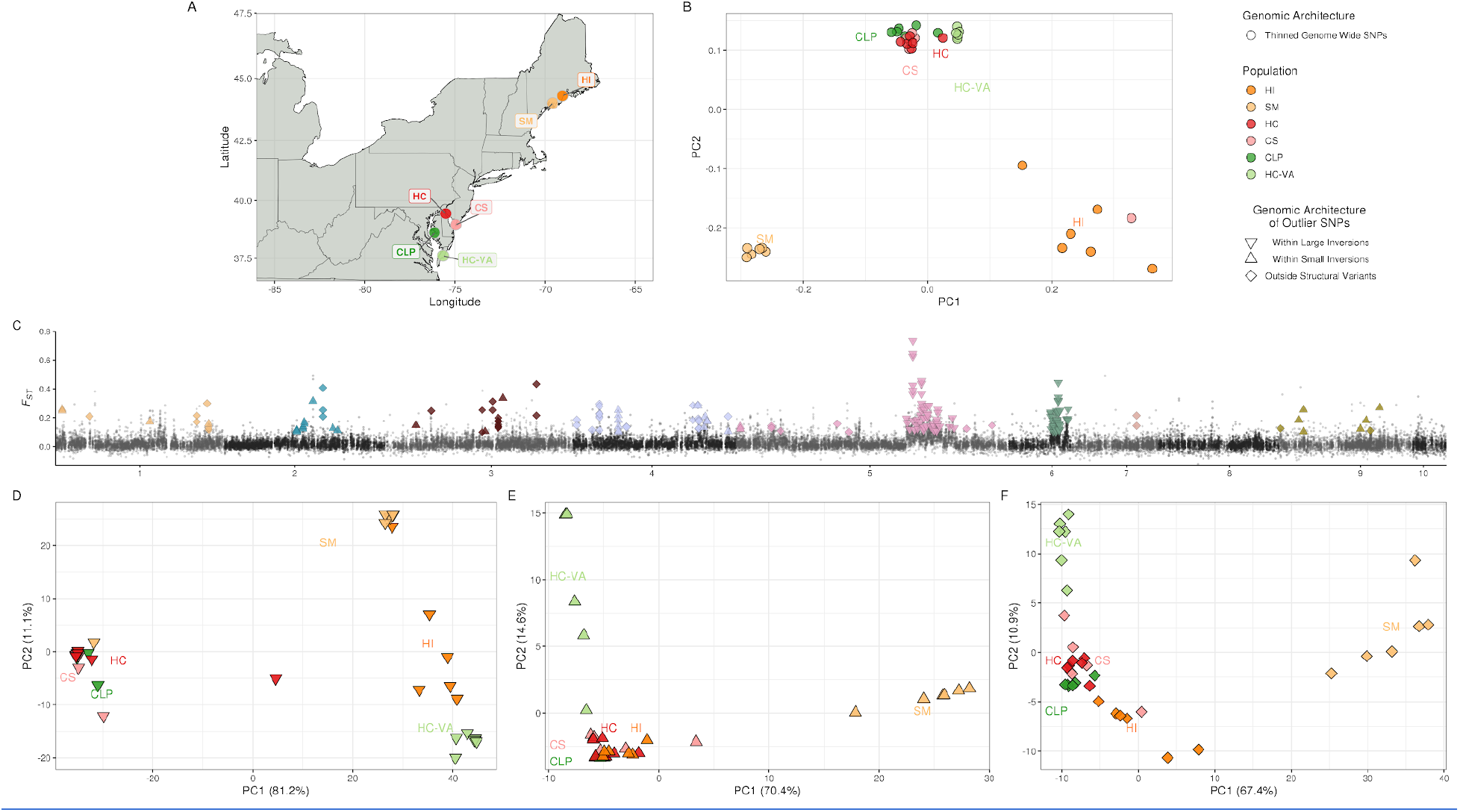
Outlier SNPs from genomic windows with large signals of elevated *F*_*ST*_ in Atlantic wild populations. **PCA analysis of outlier SNPs across Atlantic wild populations**. Outliers genomic windows outside with large signals of elevated *F*_*ST*_ identified across genomic architecture. (A) Map of Atlantic wild sampling locations. (B) PCA analysis of all SNPs after LD clumping and outlier removal using bigsnpr (Privé et al. 2018). (C) Manhattan plot of *F*_*ST*_ values calculated by OUTFLANK and averaged across 10kb windows. Outlier windows are highlighted by different colors per chromosome (yellow-Chr 1; teal-Chr 2; maroon-Chr 3, light blue-Chr 4, pink-Chr 5, green Chr 6, peach-Chr 7, gold-Chr 8). The genomic architecture of the outlier window is signified by the shape of the highlighted points (Upside down triangle-within large inversion regions, triangle-within small inversion regions, diamond-outside of structural variants). (D-F) PCA analysis (without LD clumping or outlier removal) with outlier SNPs from different architectures:(D)-within large inversions, (E)- within small inversions, (F)-outside of structural variants. Colors are representative of sampling location and shape of genomic architecture matching (C).

Genotype plots clustered by similarity provide additional information about alleles and genotypic state separating individuals. For outlier SNPs within the two large inverted regions on chromosomes 5 and 6, genotypes were similar to outliers inferred from the full dataset with the individuals separated by PC1 (from SM, HI, and HC-VA) being largely homozygous for the reference allele on chromosome 5 and all others being predominantly homozygous for the alternative allele (<u>Figure S12</u>). The one exception being HC_6 which was heterozygous for nearly all loci and SM_2 which was homozygous for the alternative allele. The remaining SNPs on the chromosome 5 inversion are from a second region from 64,030,000 to 80,150,000 which corresponds to a linkage group that is likely to actually be on chromosome 6, and those genotypes largely match the ones from the large inversion on chromosome 6. For outlier SNPs within smaller inverted regions, PC1 largely distinguished between SM individuals and all others (<u>Figure S13</u>). For SNPs on chromosomes 1,2,4,5, and 9, SM individuals were homozygous for the alternative allele or heterozygous, while the majority of other individuals were homozygous for the reference allele. SNPs on chromosome 3 showed a slightly different pattern with five of the six HC-VA individuals being distinct from all others with homozygous for the alternative allele or heterozygous genotypes. This partitioning of variation was also captured on PC2. For chromosomes 3 and 4, one CS individual clusters similarly to the SM individuals, and this individual does separate slightly from all other individuals on PC1. For outlier SNPs outside of all known structural variants, the PCA was able to distinguish SM individuals from all others as well as to largely partition out HC-VA and HI individuals from other samples. Unlike the other outlier genotype plots, the genotypes of outlier SNPs outside of structural variants showed a more regional pattern of differentiation (<u>Figure S14</u>). For example, SNPs on chromosome 2 showed both HI and SM individuals with similar genotypes and more heterozygous genotypes than all others. Similarly on chromosome 6, SM individuals were largely homozygous for the alternative allele or heterozygous across all loci, and HI individuals had a few loci with the SM pattern while all other individuals with the exception of a CS individual here homozygous for the reference allele at all chromosome 6 loci. On chromosomes 3,4,5, and 9, SM individuals were distinct from all others with largely homozygous for the alternative allele or heterozygous genotypes across all loci. Outlier SNPs on chromosome 1 were partitioned largely by locality, with individuals from SM, CLP (and one HC individual), and HC_VA clustering into distinct groups from all others. Groups were discernible by shared homozygous for the alternative allele or heterozygous genotypes for a particular subset of loci.

##### Signals of selection by salinity based on pairwise contrasts within estuaries

We investigated pairwise patterns of allele frequency differentiation between high and low salinity samples within estuaries. Using 1 kb windows of Hudson’s *F*_*ST*_, genome scans based on top 99.9% *ZF*_*ST*_ percentile have identified 897, 980, and 888 outlier windows in contrasts CS(mean 20 psu) - HC(11) in Delaware Bay, HC-VA(30) - CLP(12) in Chesapeake Bay, and HI (29.9) - SM (salinity unknown) in adjacent Maine rivers, respectively. After merging overlapping or adjacent outlier windows, a total of 229, 118, and 173 independent outlier segments were found for these three contrasts. To test for any parallel response to salinity adaptation, we checked for shared (overlapping) outlier windows among contrasts. Outlier windows were shared at only one locus on chromosome 2, and only between the Chesapeake Bay and Delaware Bay contrasts. The shared outlier segment included 7 parallel shared outlier windows spanning 3 adjacent genes: protein phosphatase 1 regulatory subunit 7-like (PPP1R7, LOC111121780), CDP-diacylglycerol--glycerol-3-phosphate 3-phosphatidyltransferase (PGS1, LOC111118137), and dynein beta chain, ciliary-like (DYHC, LOC111120585). Close inspection in Manhattan plots comparing differences in heterozygosity as well as *F*_*ST*_ revealed stronger footprints of selection (i.e., high *F*_*ST*_ coupled with high disparity in diversity relative to background) in the Chesapeake contrast where average salinity differences were greater (i.e. HC-VA - CLP, Figure 6b). In this shared outlier segment relatively lower heterozygosity was observed in both low salinity populations where selection was presumably stronger. The opposite contrast between these two estuaries, low-low salinity *F*_*ST*_ compared with high-high *F*_*ST*_, contained no shared outliers. Across the entire 3-gene locus we found homozygosity for the reference allele in most individuals at low salinity, and at the high salinity Chesapeake Bay location (HC-VA), the alternate homozygote was most frequent. The “high” salinity location in Delaware Bay (CS), which experiences an intermediate salinity regime (average of 20 instead of 30 psu), showed mixed genotypes.

**Figure 6.**
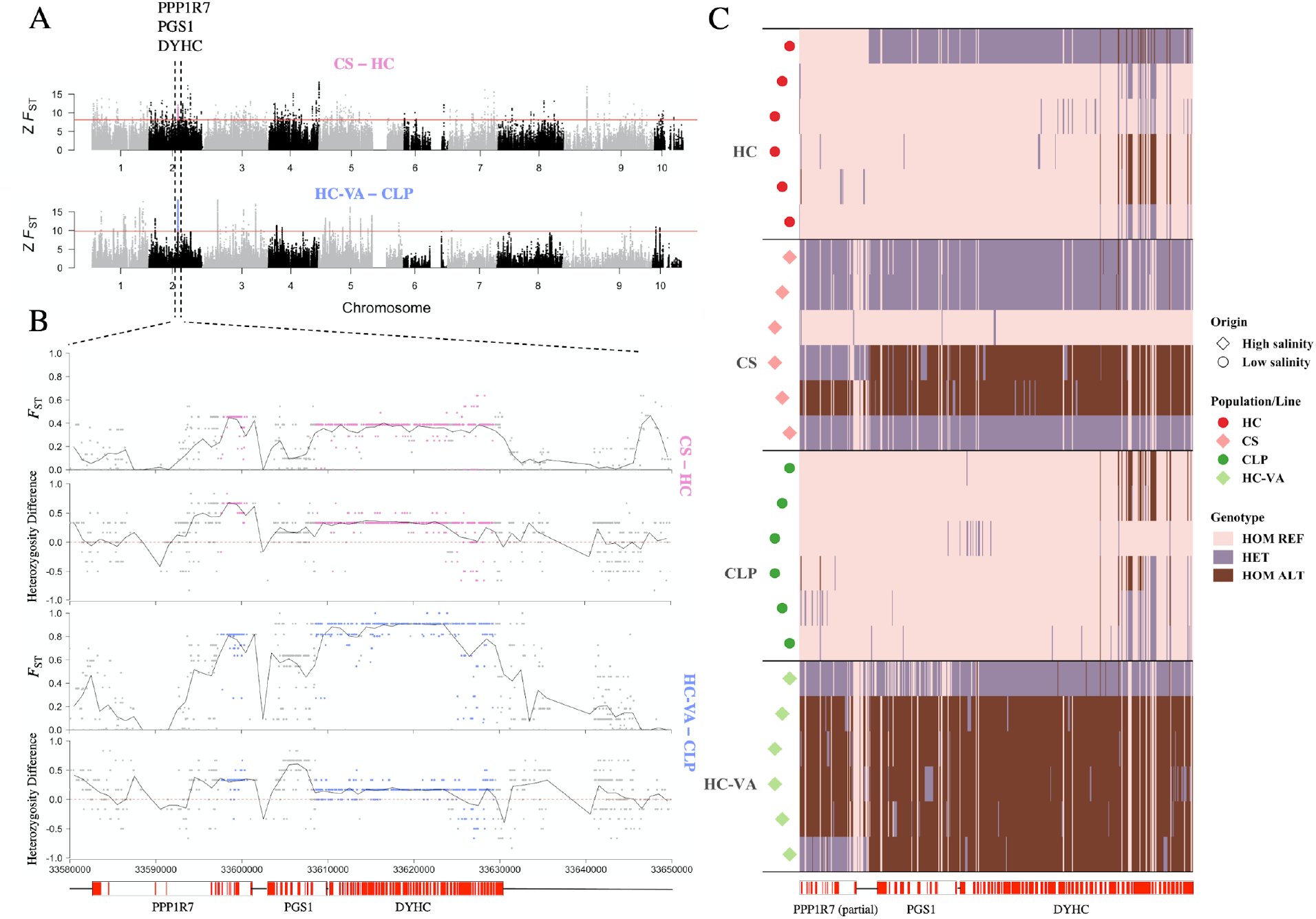
Genomic context and detailed manhattan plots for *F*_*ST*_ and heterozygosity differences, as well as genotype patterns in the shared outlier locus on chromosome 2 between high and low salinity samples. **Candidate non-neutral patterns at the only shared outlier locus between high and low salinity samples**. Outliers from the pairwise within-estuary salinity contrasts between Delaware Bay (pink) and Chesapeake Bay (Blue) are shown in Z-transformed *F*_ST_ (Z*F*_ST_) Manhattan plots at the genome level (A) based on 1Kb sliding window with 200 bp steps, with red lines demarcating the upper 99.9th percentile. Detailed Manhattan plots (B) show SNP-specific *F*_ST_ and heterozygosity difference (high salinity minus low salinity) for the only locus with shared outliers, using colored points within the genomic shared windows to indicate significance. The solid lines depict the average *F*_ST_ and heterozygosity difference based on 1Kb windows. Genes or partial genes coincident with the shared outlier segment are shown in register with coordinates on the X axis, labeled with gene symbols. Genotypes within the shared outlier segment (C) are shown with a heatmap indicating homozygosity for the reference allele at nearly all SNPs in low-salinity samples.

In contrast to these few shared differences, there were a larger number of sites on every chromosome that were outliers for genetic differentiation between higher and lower salinity sites within each estuary that were not shared among estuaries. <I recommend some quantification of this in terms of a Venn diagram or something less formal>.

## DISCUSSION

In species with high genetic diversity and a life history that enhances within-generation responses to selection, comprehensive whole-genome surveys of polymorphism can be especially important for revealing signatures of natural and artificial selection. Here, we used whole genome re-sequencing data from 90 individuals spanning distinct geographic regions, salinity regimes, and exposures to artificial selection to assess broad scale patterns in divergence, selection, and admixture across multiple levels of genomic architecture in a critical ecosystem engineer and aquaculture species. Our data confirmed the phylogeographic break between eastern oyster populations in the Gulf of Mexico (GoM) and the Atlantic coast of the USA and quantified its genomic extent. Populations on either side of this break showed multiple loci with fixed differences inside and out structural variations, including two large chromosomal inversions differentiating Atlantic and GoM populations. We also demonstrated that domestication has artificially admixed genetic material between the two ocean basins, and this admixture has helped preserve genetic diversity in these selected lines. While nucleotide diversity and heterozygosity are high in both wild and selected populations, we demonstrated that, when controlling for domestication admixture across ocean basins, wild populations showed significantly higher levels of nucleotide diversity and copy number variation than selected lines. Within the Atlantic coast, we present evidence of (a) subtle but distinct population structure, (b) introgression of selected lines within wild individuals, (c) an interaction between genomic architecture and putatively adaptive population structure, and (d) potential evidence of candidate genes responding to selection from salinity.

### Broad scale divergence across ocean basins and selected lines

The clear genetic break we detected between eastern oyster populations inhabiting estuaries along the western Atlantic coast and the Gulf of Mexico using SNPs distributed throughout the genome is supported by several previous studies with small (5 – 58 loci) panels of varied marker types including mitochondrial DNA RFLPs (Reeb and Avise 1990), nuclear DNA RFLPs (Karl and Avise 1992; Hoover and Gaffney 2005) and SNPs (Thongda et al. 2018). Observed genetic differences between Atlantic and Gulf of Mexico (GoM) populations likely are a consequence of complex historical and contemporary hydrogeographic features that regulate gene flow potential, especially across Cape Canaveral, FL ((Hare and Avise 1996; Murray and Hare 2006)). The broad scale pattern of differentiation is consistent for both the native wild populations and selected lines examined in our study (Figure 1), further underscoring a prominent role for neutral vicariant processes in driving this specific example of genetic differentiation (Murray and Hare 2006).

We also detected a number of outlier loci that showed patterns of near fixation across ocean basins. Some of these outlier loci were within four large inverted regions of the genome along chromosomes 1, 5, and 6. Three of these inverted regions showed a strong pattern of fixation of the alternative allele in GoM samples, and one inverted region (first region on chromosome 5) showed fixation within the GoM and differentiation within the Atlantic wild samples as well. Inverted genomic regions may play an important role in local adaptation, especially in high-gene flow species (Tigano and Friesen 2016; Wellenreuther and Bernatchez 2018; Faria et al. 2019; Wellenreuther et al. 2019), by effectively limiting the homogenizing effects of gene flow (Tigano and Friesen 2016; Kirkpatrick 2010; Fuller et al. 2019; Schaal, Haller, and Lotterhos 2022). Given the current data set, it is not possible to infer if these large inversions evolved during periods of high gene flow between GoM and Atlantic samples, but patterns of *F*_*ST*_ match expectations of a highly polygenic architecture, moderate to strong selection, and high levels of gene flow ((Schaal, Haller, and Lotterhos 2022). Outlier SNPs within smaller inversions (Figure S10) and outside of all detected structural variants (Figure 4) also showed large amounts of fixation between ocean basins, but with noticeably more heterozygous genotypes within each ocean basin, highlighting the interplay between divergence, selection, and recombination across the eastern oyster genome.

Across all samples, the second biggest distinction in population structure was directly related to the selected lines. The genetic differentiation patterns we saw in selected lines (based on both neutral and outlier SNPs) can be explained largely by the geographic origin of the founding populations and their selection history (Table 2). NEH and UMFS lines, both of which are derived from Long Island Sound oysters, are positioned similarly along principal component 1 in Fig 2. Separation of the two groups along principal component 2 could reflect differences in the site at which selection occurred and the extent of selection. NEH has been under selection for fast growth and resistance to the protozoan parasites *Haplosporidium nelsoni* (causative agent of MSX disease) at Cape Shore, NJ since 1966 and *Perkinsus marinus* (causative agent of Dermo disease) since 1990 (∼16th generation as of 2016 when oysters were sampled for this study) (Yu and Guo 2004), while UMFS underwent only two generations of intense selection for fast growth in cold water and survival to epizootics of Juvenile Oyster Disease (JOD), and were then maintained in the Damariscotta River, ME, where JOD is now enzootic (Yu and Guo 2004; Davis and Barber 1999). The OBOYS line is the most highly differentiated from NEH and UMFS based on PCA and pairwise *F*_*ST*_ (0.179 and 0.163, respectively). This line derives from wild oysters collected from Oyster Bayou (Cameron Parish), LA that were spawned and selected for four generations at Grand Isle, LA where *P. marinus* (but not MSX) is enzootic (Yu and Guo 2004; Casas et al. 2017; Leonhardt et al. 2017). DEBY and LOLA, the selected lines produced and maintained by the oyster breeding program at the Virginia Institute of Marine Science, exhibit intermediate levels of differentiation between the Maine, New Jersey, and Louisiana lines. DEBY, sourced from several wild Delaware Bay populations that survived an intense selection event caused by an outbreak of MSX disease, has been undergoing selection for 13 generations in the York River, VA where MSX and Dermo disease are enzootic (Frank-Lawale, Allen, and Dégremont 2014). The LOLA line is a mix of LA germplasm that was selected in the Coan River, VA for at least four generations. Wild Chesapeake Bay and LA genetic material has been introgressed with DEBY and some introgression of LOLA and DEBY has also occurred (SK Allen Jr. personal communication).

**Table 2.** Selected line information

| Line | Primary Geographic Origin | Current Genetic Make-up | Site of Selection | F0 | Gen of Select | Selection Conditions | Reference | N |
| --- | --- | --- | --- | --- | --- | --- | --- | --- |
| UMFS | Long Island Sound (Oyster Bay, NY) | 100% LIS | Upper Damariscotta River, ME | 1986 | 2 | ROD endemic; Salinity ~28-31 ppt | Davis and Barber 1999 | 6 |
| NEH | Long Island Sound (Milford, CT and Oyster Bay, NY) | 100% LIS | Cape Shore, Delaware Bay, NJ | 1966 | 16 | MSX and Dermo endemic; Salinity ~18-22 ppt | Ford and Haskin 1987; Yu and Guo 2004 | 6 |
| DEBY | Delaware Bay | 77% Delaware Bay DEBY; 14% Mobjack Bay, VA; 9% Wild Grande Terre, LA | York River, Chesapeake Bay, VA | 1987 | 13 | MSX and Dermo endemic; Salinity ~18-23 ppt | Ragone Calvo et al. 2003; S.K. Allen Jr., pers. comm. | 6 |
| LOLA | Louisiana | 10% Wild Caminada Bay, LA; 40% Wild Grande Terre, LA; 30% OBOY; 20% DEBY | Coan River, Chesapeake Bay, VA | 2008 | 4 | Salinity 8-15 ppt | S. K. Allen Jr., pers. Comm. | 6 |
| OBOY | Louisiana | 100% Oyster Bayou, Cameron Parish, LA | Grand Isle, Gulf of Mexico, LA; | 1999 | 4 | Dermo endemic; Salinity ~20 ppt | Leonhardt et al. 2017; Casas et al. 2017 | 6 |

### Genomic impacts of domestication

Looking across all samples, measures of within population genetic variation (nucleotide diversity, heterozygosity, and Tajima’s D) and copy number variation (diversity and allelic richness) appear fairly uniform across the sampled populations and lines, indicating a minimal effect of selective breeding. However, the mixed ocean basin ancestry of the LOLA and DEBY lines obscure a detectable loss of genetic diversity from domestication. For nucleotide diversity, there is a small but significant difference in mean *π* in wild populations vs selected lines when looking across all individuals (0.0054 vs 0.0053), but this difference increases when the analysis is restricted to only wild and selected localities that originate entirely from the Atlantic coastline and SNPs that are only variable within the Atlantic (0.0043 vs. 0.0038). CNV allelic richness showed a similar pattern, with wild samples having significantly higher allelic richness than selected lines when restricted to Atlantic originating samples and CNVs that were variable only within the Atlantic (Figure S6, S8). CNV diversity showed a similar pattern, but differences between wild and selected groups in the controlled analysis were not significantly different (Figure S7,S8). Only observed heterozygosity was higher in selected lines than wild populations in all comparisons.

Other aquaculture species have shown similar patterns in comparing wild vs selected (domesticated) lines. Similar levels of heterozygosity between wild populations and selected lines are common in salmonid species that also are in the early stages of domestication. Hatchery-based propagation and genetic improvement of Atlantic salmon was initiated in Norway in the 1970’s (Gjedrem and Rye 2018). Additional breeding programs were subsequently established in Finland, Scotland, Ireland, Chile, the United States, and Australia (López, Neira, and Yáñez 2014). In the Baltic Sea, nearly identical genetic diversity in hatchery stocks and their wild progenitor populations was quantified (based on microsatellite markers) with an average loss of heterozygosity in the hatchery fish of only 1.4% (Koljonen et al. 2002). A comparison of four wild Norwegian populations and Cermaq broodstock farmed in British Columbia (but derived from the original Norwegian Mowi founder strain) genotyped at 6.5K SNP loci detected no significant difference in heterozygosity or inbreeding between the wild and farmed fish after 12 generations of selection (Gutierrez, Yáñez, and Davidson 2016).

Furthermore, the positive Tajima’s D values detected in both wild and selected eastern oyster populations in our study are consistent with population contraction caused by overfishing, habitat loss, and increased intensity of abiotic and biotic stressors over the last few centuries (Gutierrez, Yáñez, and Davidson 2016; Zu Ermgassen et al. 2012). The slightly, but significantly higher Tajima’s D genomic average in the selected group may reflect the loss of rare alleles and preservation and persistence of heterozygous loci present in the small number of line founders (Ross-Ibarra et al. 2008; Puzey, Willis, and Kelly 2017). This hypothesis is supported by higher levels of linkage in the selected lines and slower decay of linkage across genomic distance in the selected lines relative to the wild populations (Figure S3).

### The intersection of natural selection, artificial selection, and genomic architecture

The selected lines DEBY and LOLA have ancestry from both ocean basins and can provide insights into the interaction of genomic architecture and selection in the eastern oyster genome. The DEBY line, originally sourced from the Delaware Bay, has undergone 13 generations of selection and during that time has been purposefully introgressed with wild oyster populations from Louisiana (estimated at 9%; SK Allen Jr. personal communication). The LOLA line was founded with multiple Louisiana oyster populations, but has also been purposefully introgressed with DEBY. The LOLA line is estimated to be 20% DEBY genetic material from controlled husbandry, so approximately 18.2% Atlantic genetic heritage. Results from PCA and *F*_*ST*_, as discussed above, are generally consistent with these percentages of introgression.

Comparing the mixed basin heritage percentages of DEBY and LOLA to the genotypes at outlier loci, selection may be maintaining higher proportions of admixture at these loci. This pattern is most pronounced in outliers within large inversions (Figure 3). The expectation would be that the large inversions would reduce recombination and effective gene flow in these regions (Tigano and Friesen 2016; Fuller et al. 2019; Kirkpatrick 2010); our results indicate that the majority of DEBY and LOLA individuals are heterozygous for alleles that are nearly fixed in each respective ocean basic (with the exception of the first inverted region on Chromosome 5 which has both alleles in the Atlantic). A plausible explanation is that these inversions played a role originally in local adaptation (Tigano and Friesen 2016; Wellenreuther and Bernatchez 2018; Wellenreuther et al. 2019; Faria et al. 2019) for eastern oysters, and now artificial selection during the creation of selective lines are promoting balanced polymorphisms at these loci. This pattern is present, though less apparent, in the outlier loci within smaller inversions (Figure S10; strikingly LOLA is more heterozygous than DEBY at those loci) and within outlier loci that are outside of detected structural variants (Figure 4). Our sample sizes per line are small, and this pattern could be a sampling artifact, but as the LOLA and DEBY lines undergo further generations of selection, combining phenotypic QTL markers and further genomic assays may provide critical information about both natural selection and the genomic basis of traits in the eastern oyster.

### The population genomic structure of wild Atlantic oysters

The wild Atlantic populations included in this study have subtle but distinct population structure. The limited Atlantic coast population structure evident in both the principal component and pairwise *F*_*ST*_ analyses (Figure 1) may be a consequence of substantial genetic exchange promoted by widespread larval dispersal between oyster populations inhabiting estuaries along the east coast of North America (Hare and Avise 1996)). However, the existing level of homogeneity also could have resulted from large-scale historical dredging and transplantation of wild oysters between Delaware Bay, Chesapeake Bay, and northeast U.S. estuaries during the 19^th^ and early 20^th^ centuries to supplement native, depleted oyster fisheries (MacKenzie and United States. National Marine Fisheries Service. Scientific Publications Office 1996). While it is beyond the scope of this study to determine the mechanisms responsible for the structure in Atlantic oyster populations, our results are consistent with previous reports (Hoover and Gaffney 2005; Thongda et al. 2018; Bernatchez et al. 2019).

Outlier detection methods found an order of magnitude fewer outlier loci within the wild Atlantic populations compared to the full data set (1,660 SNPs vs 24,269 SNPs) and a much smaller percentage of the overall loci (0.03% vs. 0.43%). The difference in absolute and relative number of outlier loci may be due to a lack of power, as the sample size of individuals is more than halved (78 to 36) when limiting analysis to only wild Atlantic populations, but it may also reflect the complex relationship wild oyster populations and human mediated activities such as restoration and aquaculture. Examining outlier loci across different levels of genomic architecture paints a picture of this genomic mosaic. PCA analysis shows distinct patterns for the different groups of loci (Figure 5), and genotype plots help to provide some clarification (Figures S12,S13,S14). Outlier loci within large chromosomal inversions indicate that there has been introgression of selected line genetic material into individuals from SM, HI, and HC-VA. The first large inversion on chromosome 5 distinctly separates these three localities from all others, as they are nearly all homozygous for reference alleles (Figure 5;S12). Looking back at the entire data set, outlier loci on the first large inversion on chromosome 5 do not strictly line up with ocean basin of origin (Figure 3). All GoM samples are homozygous for the alternative allele, but so are HC, CS, and CLP. Heterozygous individuals are found only in wild individuals (HC_6) and selected lines UMFS and LOLA. With the data in hand, it is not possible to discern between selection, introgression, or incomplete lineage sorting driving this pattern, but given the geographic distribution, intense selection or introgression of selected lines seems more likely than incomplete lineage sorting. The other two large chromosomal inversions separate out HI and HC-VA. These two inversions showed a clear pattern fixation by ocean basin of origin in the full dataset, suggesting that the observed pattern may be again introgression from selected lines with mixed basin ancestry. Other analyses, including the overall PCA and Atlantic only structure analysis, already suggested that HI individuals have selected line ancestry. One possibility is that the introgression in HC-VA happened several generations ago, so that the strongest evidence is in recombination limited outliers. Outlier loci within smaller inversions differentiate SM individuals from all others along PC1 (explaining 70.4% of the variance) and HC-VA individuals along PC2 (explaining 14.6% of the variance; Figure S13). Genotype plots show that loci on chromosomes 1,2,4,5,9 most SM individuals are either heterozygous or homozygous for the alternate allele while most others are homozygous for the reference allele, and this pattern is similar for HC-VA individuals on chromosome 3. It seems possible that this could be a pattern of selection,incomplete lineage sorting, or demographic processes. Lastly, outlier loci outside of all structural variants, again strongly differentiate SM from all others (PC1 67.4% variance, but also differentiate HC-VA and HI across different ends of PC2 (10.9%). Of note in these loci, is that on chromosome 2 and chromosome 7, genotypes are similar between HI and SM, the two most northern populations, perhaps indicating some role of environmental selection. Otherwise, most of the loci clearly differentiate SM individuals which are either heterozygous or homozygous for the alternate allele while most others are homozygous for the reference allele. Clearly, the SM population is distinct from the other wild Atlantic populations and it is difficult to simply explain this pattern with selection, introgression, or drift alone with the data in this manuscript. One possible caveat is that this study does not survey any of the northern range of the eastern oyster beyond US waters, and individuals in the SM locality could have had historical or contemporary gene flow (natural or human-mediated) with more northern populations.

### Parallel adaptive divergence within estuaries

It is hypothesized that high gene flow within estuarine gradients generally prevents local adaptation to physically marginal habitats (Sanford and Kelly 2011). For species with high fecundity and early mortality, however, the recurrent viability selection across habitat heterogeneities may contribute to spatial population differences among adults (Pavey et al. 2015; Schmidt and Rand 2001). For *C. virginica*, evidence has shown that adults collected from low and high-salinity reefs differ in their gene expression profiles across several common garden treatments (Eierman and Hare 2016). Similarly, reciprocal transplants and challenge experiments support within-estuary local adaptation of *Ostrea lurida* to hyposalinity stress (Bible and Sanford 2015). Here we observed subtle genome-wide differentiation between low and high-salinity adult populations within several estuaries where genetic homogeneity predominates (Figure 1). A genomic scan using Z*F*_ST_ windows identified hundreds of outlier segments in each independent estuarine contrast, and a single shared segment exhibiting parallel allele frequency differentiation in two estuaries (Figure 6; see outliers and their annotations in <u>Table S5</u>). Hyposalinity selection is suggested by the pattern of almost complete homozygosity for the reference allele in low salinity up-bay populations of both estuaries, whereas genotypic patterns were mixed down-bay. Although other environmental stressors may be involved in this spatially divergent selection (like MSX and Dermo disease), disease is typically minimal in low salinity up-bay regions (Munroe et al. 2013).

Heritability of extreme hyposalinity survival in *C. virginica* has been estimated to be moderate (McCarty et al. 2020). A subsequent study using Chesapeake Bay oyster families found several significant QTL for this trait and days to death. QTL were on chromosomes 1 and 7, but genomic prediction accuracies also indicated a polygenic basis including many genes of small effect (McCarty, Allen, and Plough 2022). Under these conditions (as with QTL studies) there may be little likelihood of finding parallel adaptive architectures, especially if pleiotropy supports many polygenic ‘solutions’ to the adaptive challenge of the moment (Lotterhos 2022). Here, the outlier segment shared between Delaware and Chesapeake Bay populations pairs included three adjacent genes: protein phosphatase 1 regulatory subunit 7-like (PPP1R7), CDP-diacylglycerol--glycerol-3-phosphate 3-phosphatidyltransferase (PGS1), and dynein beta chain, ciliary-like (DYHC). PPP1R7 plays a pivotal role in glucogen formation and metabolism, potentially providing the energy required by osmoregulation and cellular homeostasis (Yan et al. 2017). DYHC is a component of the axonemal dynein complex (GO:0005858), a motor protein that powers cilia/flagellar beating and therefore is likely to have an important impact on both feeding and oxygen supply over bivalve gills (Maynard et al. 2018; Roberts et al. 2013). It is noteworthy that dynein heavy chain 1 axonemal was significantly upregulated in *O. lurida* oysters from a low salinity source population, in response to low salinity treatment, but the same was not true of oysters from moderate and high salinity source populations that had all shared a common environment for two generations (Maynard, et al., 2018).

### Anthropogenic impacts on wild oyster populations

While domestication of the eastern oyster for aquaculture is relatively nascent (Gjedrem and Baranski 2009; Davis and Barber 1999; Calvo et al. 2003; Frank-Lawale, Allen, and Dégremont 2014), human impacts on the eastern oyster are far more extensive, including large-scale transplantation and restoration that dates back to eat least the 19th century (MacKenzie and United States. National Marine Fisheries Service. Scientific Publications Office 1996; Grabowski, Conrad, and James 2012). Our data indicate that there has been introgression of genetic material from non-local oysters to several “wild” oysters. One “wild” individual from CS (CS_3), appears more similar to the DEBY and LOLA selected lines than any other Atlantic wild oyster and likely is not a wild oyster. Structure results also indicated that five of the six HI individuals surveyed have shared ancestry from the UMFS and NEH selected lines. The NEH line has been widely used by Maine hatcheries for commercial seed production for aquaculture (Guo, 2021). Looking at outlier loci within large inversions, it appears likely that at least one individual from wild localities five different wild localites have linked groups of outlier loci with alleles that are more common in the GoM or in selected lines with mixed ocean basin (Figure 3). Similar patterns appear in the outlier loci within smaller inversions (Figure S10), and loci outside of structural variants (Figure 4). Overall, our sample size per locality and line is relatively small, and it is well beyond the scope of this study to investigate human mediated admixture and introgression. With that in mind, some of our results could indicate retention of ancestral variation or potentially similar selective pressures for selected lines and individual wild populations. A recent study looking at natural and restored oyster reefs was able to detect population structure between natural and restored reefs and show that most of the structure was attributable to environmental conditions and not restoration status (Hornick and Plough 2021). Though, given the results from our study, we suggest that the impact of human actions on the eastern oyster genome has been extensive in “wild” populations, and outlier loci within inverted regions may hold clues to different anthropogenic impacts that may otherwise be quickly covered up by recombination.

## Notes

### Competing Interest Statement

The authors have declared no competing interest.

