## Supplemental Information for "Nucleotide and structural polymorphisms of the eastern oyster genome paint a mosaic of divergence, selection, and human impacts"

Supporting Information

### Supplemental text

The extent of Eastern oyster domestication varies by geographic region but is generally based on increased performance (survival) following natural disease epizootics (Table 2). The first documented Eastern oyster breeding program was initiated at Rutgers University in 1960 after an outbreak of a haplosporidian parasite which causes MSX disease killed 90% of the oysters in southern Delaware Bay (DB) planting grounds. Survivor progeny exhibited significantly better performance in response to subsequent MSX exposures, prompting multiple generations of selection across a diverse collection of naïve wild populations with DB and Long Island Sound (LIS) origins at an MSX-endemic site in Cape Shore, NJ (Haskin & Ford, 1979; Ford & Haskin 1987). After nearly 30 years and seven generations of mass selection, extant inbred Rutgers lines originating from the Connecticut coast of LIS were crossed to create the Northeast High Survival line (NEH) in 1992 (Yu & Guo, 2005). The NEH line is represented by several sublines, which are inbred for mass selection and hybridized for production (Guo, 2021). Oysters used in this study were from an inbred subline NEH93, which were spawned in 2016 as the 16th generation since the founding of the NEH lines. Genetic material from Long Island Sound has been incorporated into NEH lines several times during its breeding history. Selection of NEH is ongoing at the Cape Shore site, where a second oyster parasite, *Perkinsus marinus* (causative agent of Dermo disease) is also endemic.

In 1987 the Virginia Institute of Marine Science (VIMS) also began breeding Eastern oysters. The initial goal of the VIMS program was to select oysters for dual resistance to MSX and Dermo by deploying progeny of wild oysters, collected from several MSX-endemic areas throughout southern DB, in the York River, VA where both diseases are enzootic (Ragone Calvo, Calvo, & Burreson, 2003). After four generations of selection and inbreeding, genetic material from a wild Chesapeake Bay (CB) population (Mobjack Bay) and three Gulf of Mexico (Louisiana (LA)) populations was introgressed with the VIMS DEBY line over a ten-year time period (Frank-Lawale, Allen, & Dégremont, 2014). DEBY was actively selected in the York River through 2016 (13 generations) and broodstock continues to be propagated and licensed to commercial aquaculture operations. To accommodate Eastern oyster culture across the wide-ranging CB salinity gradient, VIMS expanded their breeding program to include wild Louisiana germplasm and low-salinity test sites. In 2008, high-performing VIMS lines with LA ancestry were crossed to create the LOLA line. LOLA was actively selected in the Coan River, VA through 2016 for at least four generations. VIMS still maintains the LOLA line and distributes broodstock to industry (SK Allen Jr. personal communication).

A breeding program aimed at developing Dermo-resistant oysters for commercial use in the Gulf of Mexico was initiated in 1999 with wild oysters collected from Oyster Bayou (Cameron Parish), LA. Oyster Bayou oysters were spawned and their progeny selected at the Louisiana Sea Grant Oyster Hatchery (Grand Isle), where Dermo disease is endemic. The resultant OBOY line was subject to four rounds of selection; two (F0 and F3) in the field and two (F1 and F2) in response to controlled disease challenge in the laboratory. Survivors of each selection event were used as broodstock for the next generation (Casas et al., 2017; Leonhardt, Casas, Supan, & La Peyre, 2017).

Unlike the mid-Atlantic and LA breeding programs, fast growth, not disease resistance, was the main selection target at the University of Maine. Most Eastern oyster culture in Maine occurs in the Damariscotta River, where cooler temperatures limit the growing season to just four months of the year (Proestou et al. 2016). In 1986, fast-growing oysters were imported from a commercial oyster aquaculture company, Frank M. Flowers & Sons (FMF) in New York, to found the University of Maine Flowers Select (UMFS) line. FMF oysters deployed in the Damariscotta River were subject to truncation selection, whereby the largest 20% of survivors were used as broodstock for subsequent spawns. After two generations of selection, selected oysters were significantly larger than unselected controls (Davis & Barber, 1999). During UMFS performance trials, UMFS oysters were exposed to an endemic bacterial pathogen, Roseovarius Oyster Disease (ROD), that causes acute mortality in oysters less than 30 mm in size (Maloy, Ford, Karney, & Boettcher,2007). The ROD exposure, coupled with selection of the largest surivors, likely resulted in a faster growing line with some ROD-resistance.

### Supplemental Tables

###### Table S2- Number of reads, total read mapping, number of duplicates identified, and filtered read mapping per sample

| **SAMPLE** | **Raw Reads** | **Trimmed**  **Reads** | **Total Mappings** | **Duplicates Detected** | **Mappings**  **Post Filtering** | **Percent**  **Genome Covered Total Mappings** | **Percent**  **Genome Covered**  **Filtered Mappings** |
| --- | --- | --- | --- | --- | --- | --- | --- |
| CL_1 | 61756328 | 60515962 | 58638967 | 3678309 | 42779510 | 85.3575 | 77.48 |
| CL_2 | 99774458 | 97882558 | 94839920 | 5785436 | 69565701 | 86.9888 | 79.5246 |
| CL_3 | 96409828 | 92943592 | 89955536 | 5728720 | 64336032 | 87.5182 | 79.7683 |
| CL_4 | 57385744 | 56122644 | 54376130 | 3384483 | 39568166 | 84.832 | 76.6349 |
| CL_5 | 51825176 | 49808752 | 48188040 | 2849220 | 34477409 | 84.8765 | 76.5747 |
| CL_6 | 66656598 | 64735606 | 62427500 | 3773383 | 45608980 | 85.751 | 77.9069 |
| CLP_1 | 59304898 | 57214842 | 55227242 | 3427383 | 40531929 | 86.853 | 79.8969 |
| CLP_2 | 70623926 | 68392668 | 65942791 | 4459706 | 48158612 | 87.74 | 80.9544 |
| CLP_3 | 98389272 | 94723928 | 91658037 | 5644819 | 65954366 | 88.6506 | 82.0712 |
| CLP_4 | 69924272 | 67476732 | 65189185 | 4573182 | 46891439 | 87.3222 | 80.3237 |
| CLP_5 | 84034792 | 81291734 | 78410746 | 5500463 | 57109934 | 87.9937 | 81.3275 |
| CLP_6 | 97689796 | 93990226 | 90836203 | 6891883 | 65385413 | 88.7637 | 82.3502 |
| CS_1 | 76082270 | 73729270 | 71169316 | 4784194 | 51922862 | 87.477 | 80.5646 |
| CS_2 | 72128202 | 69666346 | 67261377 | 4260850 | 49558329 | 87.3619 | 80.5359 |
| CS_3 | 68535568 | 66784598 | 64869270 | 4143160 | 48776174 | 87.9236 | 81.4316 |
| CS_5 | 75917232 | 74059434 | 71760922 | 4403643 | 52503142 | 87.2494 | 80.2374 |
| CS_6 | 64004962 | 61921988 | 59831080 | 4224183 | 43734808 | 86.6288 | 79.692 |
| CS_7 | 69777108 | 68030738 | 65877071 | 4122388 | 48391958 | 86.7929 | 79.6759 |
| DEBY_1 | 70340066 | 68129214 | 65984679 | 4204859 | 48013858 | 87.4296 | 80.5127 |
| DEBY_2 | 61820396 | 59393626 | 57501832 | 3569039 | 41033127 | 87.0883 | 79.4019 |
| DEBY_3 | 90050284 | 88202564 | 85646456 | 5163796 | 62737184 | 87.865 | 80.898 |
| DEBY_4 | 54919768 | 52712608 | 51030035 | 3250463 | 37069517 | 86.3471 | 79.0749 |
| DEBY_5 | 66912230 | 65379052 | 63242334 | 4157130 | 46365584 | 86.8178 | 79.5951 |
| DEBY_6 | 69395392 | 67069200 | 64901712 | 4409876 | 46839037 | 87.1723 | 80.1616 |
| HC_1 | 77293636 | 74906388 | 72425035 | 5835437 | 52186665 | 87.2464 | 80.3062 |
| HC_3 | 63112478 | 61557868 | 59524843 | 4346896 | 43322163 | 86.439 | 79.2234 |
| HC_4 | 79754444 | 77660564 | 75074963 | 5501949 | 54611969 | 87.5876 | 80.7381 |
| HC_5 | 90033632 | 87108604 | 84111972 | 5616048 | 61600005 | 87.9425 | 81.3642 |
| HC_6 | 56048260 | 54559620 | 52342661 | 3436210 | 38373096 | 86.5105 | 79.3349 |
| HC_7 | 154599222 | 148626848 | 143848308 | 8804482 | 103414438 | 90.2442 | 84.073 |
| HC_VA_1 | 85940448 | 82636266 | 79780756 | 5891673 | 57681262 | 88.5127 | 81.8589 |
| HC_VA_2 | 82902150 | 79755572 | 77204226 | 5527680 | 55580071 | 88.3048 | 81.6542 |
| HC_VA_3 | 63890928 | 61749506 | 59643898 | 3294694 | 44391895 | 86.7738 | 79.7823 |
| HC_VA_4 | 69606580 | 67810232 | 65615532 | 3866507 | 48549698 | 87.1651 | 80.0706 |
| HC_VA_5 | 79755810 | 77137600 | 74474190 | 5370365 | 54248567 | 87.7156 | 80.9334 |
| HC_VA_6 | 85495276 | 82438178 | 79705400 | 6118503 | 57718329 | 88.1368 | 81.4786 |
| HG_HG0F2 | 81488634 | 79609440 | 78557006 | 4643260 | 65052524 | 99.6608 | 98.1506 |
| HG_HG2F1 | 107490278 | 104562398 | 102891704 | 7466672 | 78838999 | 96.7679 | 94.1229 |
| HG_HG2M5 | 65635204 | 64848234 | 64299271 | 4276150 | 49711444 | 95.4858 | 90.9396 |
| HI_1 | 78587340 | 76634458 | 74051860 | 5107038 | 54685883 | 87.7458 | 81.1001 |
| HI_2 | 118354864 | 114591096 | 110920625 | 8030373 | 81152249 | 89.4068 | 83.4138 |
| HI_3 | 84232622 | 81870440 | 79113148 | 5748565 | 57429647 | 87.8206 | 80.9937 |
| HI_4 | 77417784 | 75521270 | 73141247 | 5281685 | 53400559 | 87.2802 | 80.4514 |
| HI_5 | 104038488 | 101912426 | 98927966 | 6886611 | 73404346 | 88.6163 | 82.3536 |
| HI_6 | 78733988 | 76799260 | 74276699 | 5147995 | 54206331 | 86.7613 | 79.7576 |
| LM_1_pool | 66969314 | 64216034 | 61769102 | 4561219 | 41620194 | 85.8385 | 77.826 |
| LM_3 | 92486240 | 88551324 | 84623488 | 5074628 | 58897350 | 86.1071 | 78.3485 |
| LM_4 | 172376322 | 166674276 | 159956314 | 9420105 | 111641861 | 88.8038 | 81.8859 |
| LM_7 | 75118986 | 72981782 | 70198561 | 4065303 | 50087656 | 85.2545 | 77.3859 |
| LM_8 | 66373270 | 64272486 | 61604541 | 3484785 | 43154467 | 83.8853 | 75.6177 |
| LOLA_1 | 92988582 | 90136906 | 86935582 | 5759948 | 63012286 | 87.7419 | 80.6833 |
| LOLA_2 | 67427816 | 65354868 | 63047433 | 3849653 | 46353891 | 86.1454 | 78.785 |
| LOLA_3 | 70995646 | 68916992 | 66510965 | 4070465 | 48922303 | 86.0747 | 78.6432 |
| LOLA_4 | 81746434 | 78906376 | 76174709 | 5454965 | 54236799 | 87.5018 | 80.3578 |
| LOLA_5 | 68704786 | 66973404 | 64653570 | 4445880 | 46785631 | 85.6134 | 77.9542 |
| LOLA_6 | 77903354 | 75647386 | 73198687 | 4979400 | 52565163 | 86.822 | 79.5417 |
| NEH_1 | 62806886 | 61154804 | 59252416 | 3971608 | 44112015 | 87.5356 | 80.9636 |
| NEH_2 | 73087636 | 71245062 | 69207906 | 4771209 | 51989894 | 87.5727 | 81.0907 |
| NEH_3 | 103930046 | 101703694 | 99125055 | 7295972 | 73002332 | 88.4021 | 81.9887 |
| NEH_4 | 116701620 | 112592462 | 109362296 | 7223266 | 80135399 | 89.2251 | 83.0431 |
| NEH_5 | 64394368 | 62707068 | 61129630 | 4349320 | 45033834 | 87.8081 | 81.1688 |
| NEH_6 | 74547538 | 72648536 | 70714452 | 5006221 | 51309059 | 87.4629 | 80.5809 |
| NG_NH0H4 | 70988650 | 69279808 | 67299078 | 4214975 | 50693549 | 87.8541 | 81.2857 |
| NG_NH2F6 | 64297714 | 62713746 | 61018545 | 4418426 | 45139472 | 85.6114 | 78.6404 |
| NG_NH2F8 | 67198670 | 65281042 | 63378230 | 4452766 | 46583461 | 85.6018 | 78.7202 |
| NG_NH2M1 | 79668514 | 77463988 | 75181995 | 5423725 | 55065837 | 87.5044 | 80.8985 |
| OBOYS2_1 | 65043442 | 63682580 | 61624990 | 3846448 | 45074106 | 85.7239 | 77.9475 |
| OBOYS2_2 | 70055162 | 68520216 | 66194378 | 4446615 | 48035537 | 86.0779 | 78.4112 |
| OBOYS2_3 | 82925952 | 79746152 | 77181900 | 4533366 | 55447440 | 86.9706 | 79.2156 |
| OBOYS2_4 | 56605592 | 54813194 | 52683361 | 3429467 | 37926799 | 85.1607 | 77.2418 |
| OBOYS2_5 | 71635564 | 70124370 | 67818633 | 4475680 | 49208853 | 86.1592 | 78.4691 |
| OBOYS2_6 | 107126892 | 104962500 | 101646882 | 6572431 | 74053208 | 87.7767 | 80.4737 |
| SL_1 | 89147578 | 86978844 | 83523180 | 5065708 | 61078007 | 86.8789 | 79.447 |
| SL_2 | 59238048 | 57637436 | 55584184 | 3426593 | 40568507 | 85.3585 | 77.5398 |
| SL_3 | 93694418 | 91917534 | 88796974 | 5843833 | 64699574 | 87.1454 | 79.744 |
| SL_4 | 61696302 | 60440650 | 58460518 | 3522935 | 42836220 | 85.6596 | 77.8405 |
| SL_5 | 99844072 | 97535802 | 94365829 | 6174554 | 68891226 | 87.5508 | 80.3312 |
| SL_6 | 76386656 | 74247366 | 71756216 | 4338350 | 52284751 | 86.4634 | 78.9062 |
| SM_10 | 83663278 | 81453140 | 78799232 | 5721628 | 57257531 | 87.3661 | 80.4979 |
| SM_11 | 63381898 | 61514580 | 57785915 | 3795790 | 41969957 | 86.6241 | 79.4197 |
| SM_12 | 71593066 | 69750482 | 67385513 | 4770414 | 49074758 | 86.8614 | 79.8611 |
| SM_7 | 95076544 | 93039658 | 90035576 | 7358208 | 64816528 | 87.5971 | 80.8177 |
| SM_8 | 71562510 | 69681052 | 67407309 | 4892705 | 49016663 | 86.9012 | 79.792 |
| SM_9 | 71256028 | 69432742 | 67144886 | 4394338 | 49081725 | 86.9997 | 79.8686 |
| UMFS_1 | 64143576 | 62429272 | 60777720 | 4095873 | 43909923 | 86.1064 | 78.7581 |
| UMFS_2 | 76203502 | 74877078 | 72822380 | 5144437 | 53057441 | 87.1714 | 80.0194 |
| UMFS_3 | 99560248 | 97380516 | 94283095 | 6361244 | 69210032 | 87.6549 | 80.9078 |
| UMFS_4 | 94236090 | 91852914 | 88970604 | 6190590 | 64826142 | 87.9809 | 81.2166 |
| UMFS_5 | 71144408 | 68855926 | 66650390 | 4162606 | 48791995 | 86.5656 | 79.5327 |
| UMFS_6 | 72412090 | 70347494 | 67922828 | 4481380 | 49830314 | 87.1443 | 80.2762 |

#### Supplemental Table S3. Table Containing different number of SNP variants across call rates and minor allele frequencies.

|  | **Call Rate** |  |  |  |
| --- | --- | --- | --- | --- |
| **Minor Allele Frequency** | **90%** | **95%** | **100%** | **100% and Biallelic only** |
| **1%** | 27,926,188 | 23,885,771 | 14,103,332 | 12,149,052 |
| **5%** | 13,748,757 | 11,638,196 | 6,699,719 | 5,574,080 |

Supplemental Table S5. Annotation for outlier windows that identifed using 1 kb sliding window of ZF_ST_ scans. A total of 229, 118, and 173 independent outlier segments (after merging overlapping or adjacent outlier windows) were found in contrasts CS(mean 20 psu) - HC(11) in Delaware Bay, HC-VA(30) - CLP(12) in Chesapeake Bay, and HI (29.9) - SM (salinity unknown) in adjacent Maine rivers, respectively. One shared outlier segment included 7 parallel shared outlier windows were also listed in the table. Chr-chromosome; Start: window start position; End, window end position; Count: number of genes that annotated in the window; Gene description: detailed gene name; Gene ID: gene identity.

| Chr | Start | End | Count | Gene description | Gene ID |
| --- | --- | --- | --- | --- | --- |
| Hope Creek (HC) - Cape Shore (CS) | | | | | |
| NC_035780.1 | 1602600 | 1608600 | 0 |  |  |
| NC_035780.1 | 2383000 | 2384200 | 1 | organic cation transporter protein-like | 111099207 |
| NC_035780.1 | 3592000 | 3595000 | 1 | ATP-dependent DNA helicase Q1-like | 111129935 |
| NC_035780.1 | 3595200 | 3596200 | 1 | ATP-dependent DNA helicase Q1-like | 111129935 |
| NC_035780.1 | 5825600 | 5826600 | 1 | mucin-2-like | 111103323 |
| NC_035780.1 | 5826800 | 5828800 | 1 | mucin-2-like | 111103323 |
| NC_035780.1 | 5829600 | 5832800 | 1 | mucin-2-like | 111103323 |
| NC_035780.1 | 5834200 | 5836400 | 1 | mucin-2-like | 111103323 |
| NC_035780.1 | 7190800 | 7191800 | 1 | lactadherin-like | 111127890 |
| NC_035780.1 | 7751400 | 7752400 | 1 | uncharacterized LOC111120458 | 111120458 |
| NC_035780.1 | 11017400 | 11020400 | 0 |  |  |
| NC_035780.1 | 11220600 | 11224200 | 0 |  |  |
| NC_035780.1 | 14528600 | 14529600 | 1 | serine/threonine-protein kinase greatwall-like | 111117050 |
| NC_035780.1 | 15584200 | 15585200 | 1 | motor neuron and pancreas homeobox protein 1-like | 111125267 |
| NC_035780.1 | 16359600 | 16361400 | 1 | uncharacterized LOC111105980 | 111105980 |
| NC_035780.1 | 16846000 | 16847000 | 1 | uncharacterized LOC111113005 | 111113005 |
| NC_035780.1 | 19760600 | 19761800 | 1 | RNA polymerase II elongation factor ELL-like | 111111579 |
| NC_035780.1 | 28296200 | 28298000 | 1 | transcription factor Sox-5-like | 111110636 |
| NC_035780.1 | 29445200 | 29448400 | 1 | uncharacterized LOC111099732 | 111099732 |
| NC_035780.1 | 46576200 | 46577200 | 0 |  |  |
| NC_035780.1 | 56033600 | 56035200 | 0 |  |  |
| NC_035780.1 | 59128000 | 59129600 | 1 | liprin-beta-1-like | 111111050 |
| NC_035780.1 | 59228200 | 59229200 | 1 | protein lin-9 homolog | 111115224 |
| NC_035780.1 | 59695200 | 59696200 | 1 | hydroxymethylglutaryl-CoA lyase, mitochondrial-like | 111116092 |
| NC_035780.1 | 59745000 | 59746000 | 1 | cryptochrome-1-like | 111103032 |
| NC_035780.1 | 63403200 | 63405200 | 1 | DNA-directed RNA polymerase II subunit RPB4-like | 111132777 |
| NC_035780.1 | 63738200 | 63740600 | 1 | zinc finger and BTB domain-containing protein 49-like | 111128076 |
| NC_035780.1 | 63743800 | 63745600 | 1 | zinc finger and BTB domain-containing protein 49-like | 111128076 |
| NC_035781.1 | 725200 | 727000 | 1 | dystrophin-like | 111117995 |
| NC_035781.1 | 2476200 | 2478400 | 1 | uncharacterized LOC111119145 | 111119145 |
| NC_035781.1 | 2770000 | 2771600 | 0 |  |  |
| NC_035781.1 | 3033000 | 3034200 | 1 | uncharacterized LOC111118480 | 111118480 |
| NC_035781.1 | 7363200 | 7364200 | 1 | GTP-binding protein SAR1-like | 111120699 |
| NC_035781.1 | 7386800 | 7387800 | 1 | WD repeat domain phosphoinositide-interacting protein 3-like | 111120978 |
| NC_035781.1 | 7389400 | 7390400 | 1 | WD repeat domain phosphoinositide-interacting protein 3-like | 111120978 |
| NC_035781.1 | 7494200 | 7496800 | 1 | calcium uptake protein 1, mitochondrial-like | 111119636 |
| NC_035781.1 | 10018600 | 10019600 | 1 | pancreatic triacylglycerol lipase-like | 111122015 |
| NC_035781.1 | 13791400 | 13792600 | 1 | MAP kinase-interacting serine/threonine-protein kinase 1-like | 111118004 |
| NC_035781.1 | 14592600 | 14593600 | 0 |  |  |
| NC_035781.1 | 15356400 | 15360400 | 2 | calreticulin-like;transmembrane protein 59-like | 111119403;111121532 |
| NC_035781.1 | 19414000 | 19415600 | 1 | uncharacterized LOC111119700 | 111119700 |
| NC_035781.1 | 20616000 | 20618200 | 0 |  |  |
| NC_035781.1 | 20620200 | 20621600 | 1 | bifunctional methylenetetrahydrofolate dehydrogenase/cyclohydrolase, mitochondrial-like | 111122078 |
| NC_035781.1 | 26624800 | 26625800 | 1 | uncharacterized LOC111118437 | 111118437 |
| NC_035781.1 | 27562600 | 27564400 | 1 | carbohydrate sulfotransferase 9-like | 111119801 |
| NC_035781.1 | 27653200 | 27654200 | 0 |  |  |
| NC_035781.1 | 27655200 | 27656200 | 1 | cytochrome P450 3A19-like | 111120209 |
| NC_035781.1 | 33483400 | 33484600 | 0 |  |  |
| NC_035781.1 | 33597600 | 33600800 | 1 | protein phosphatase 1 regulatory subunit 7-like | 111121780 |
| NC_035781.1 | 33608600 | 33610600 | 2 | dynein beta chain, ciliary-like;CDP-diacylglycerol--glycerol-3-phosphate 3-phosphatidyltransferase, mitochondrial-like | 111118137;111120585 |
| NC_035781.1 | 33610800 | 33612000 | 1 | dynein beta chain, ciliary-like | 111120585 |
| NC_035781.1 | 33612200 | 33613600 | 1 | dynein beta chain, ciliary-like | 111120585 |
| NC_035781.1 | 33614000 | 33619400 | 1 | dynein beta chain, ciliary-like | 111120585 |
| NC_035781.1 | 33619800 | 33624000 | 1 | dynein beta chain, ciliary-like | 111120585 |
| NC_035781.1 | 33624600 | 33629400 | 1 | dynein beta chain, ciliary-like | 111120585 |
| NC_035781.1 | 33645800 | 33647600 | 1 | PDZ domain-containing protein GIPC1-like | 111120753 |
| NC_035781.1 | 33647800 | 33649000 | 1 | PDZ domain-containing protein GIPC1-like | 111120753 |
| NC_035781.1 | 35299600 | 35300800 | 1 | uncharacterized LOC111120643 | 111120643 |
| NC_035781.1 | 35366400 | 35367400 | 1 | palmitoyltransferase ZDHHC6-like | 111121831 |
| NC_035781.1 | 35523200 | 35524200 | 1 | U6 snRNA-associated Sm-like protein LSm4 | 111119055 |
| NC_035781.1 | 39052200 | 39060400 | 2 | uncharacterized LOC111121479;protein zyg-11 homolog B-like | 111121479;111121423 |
| NC_035781.1 | 41683200 | 41684400 | 0 |  |  |
| NC_035781.1 | 42145800 | 42147200 | 1 | uncharacterized LOC111121431 | 111121431 |
| NC_035781.1 | 44842400 | 44844200 | 1 | transmembrane protein 180-like | 111122326 |
| NC_035781.1 | 46332600 | 46333800 | 1 | ATP-binding cassette sub-family A member 1-like | 111122236 |
| NC_035781.1 | 47950800 | 47951800 | 1 | uncharacterized LOC111120721 | 111120721 |
| NC_035781.1 | 47952000 | 47955400 | 2 | uncharacterized LOC111120722;uncharacterized LOC111120721 | 111120721;111120722 |
| NC_035781.1 | 52536200 | 52537200 | 1 | FMRFamide receptor-like | 111119110 |
| NC_035781.1 | 55301400 | 55303000 | 1 | uncharacterized LOC111120689 | 111120689 |
| NC_035781.1 | 60535000 | 60536400 | 1 | serine protease 44-like | 111121960 |
| NC_035782.1 | 6680200 | 6681200 | 1 | vacuolar protein-sorting-associated protein 25-like | 111123133 |
| NC_035782.1 | 6878000 | 6879000 | 1 | supervillin-like | 111123958 |
| NC_035782.1 | 13862400 | 13863600 | 0 |  |  |
| NC_035782.1 | 15519600 | 15520600 | 1 | uncharacterized LOC111126334 | 111126334 |
| NC_035782.1 | 15521000 | 15522200 | 1 | uncharacterized LOC111126334 | 111126334 |
| NC_035782.1 | 15523200 | 15524400 | 1 | uncharacterized LOC111126334 | 111126334 |
| NC_035782.1 | 15835000 | 15836000 | 0 |  |  |
| NC_035782.1 | 18280400 | 18282000 | 1 | neural cell adhesion molecule 1-like | 111126520 |
| NC_035782.1 | 26062200 | 26063200 | 1 | uncharacterized LOC111126539 | 111126539 |
| NC_035782.1 | 26867800 | 26869000 | 0 |  |  |
| NC_035782.1 | 33604200 | 33605800 | 1 | endoplasmic reticulum mannosyl-oligosaccharide 1,2-alpha-mannosidase-like | 111124585 |
| NC_035782.1 | 33608800 | 33610000 | 1 | endoplasmic reticulum mannosyl-oligosaccharide 1,2-alpha-mannosidase-like | 111124585 |
| NC_035782.1 | 33636600 | 33638000 | 0 |  |  |
| NC_035782.1 | 33640400 | 33641600 | 1 | eukaryotic translation initiation factor 3 subunit G-like | 111124589 |
| NC_035782.1 | 37071000 | 37072400 | 1 | uncharacterized LOC111124719 | 111124719 |
| NC_035782.1 | 37289600 | 37290800 | 1 | coronin-7-like | 111126566 |
| NC_035782.1 | 39433600 | 39435800 | 1 | E3 ubiquitin-protein ligase CBL-like | 111127555 |
| NC_035782.1 | 41377600 | 41379000 | 1 | protocadherin beta-8-like | 111125411 |
| NC_035782.1 | 43173200 | 43174400 | 0 |  |  |
| NC_035782.1 | 45145400 | 45146400 | 1 | sodium bicarbonate cotransporter 3-like | 111126568 |
| NC_035782.1 | 46722600 | 46723800 | 1 | transient receptor potential cation channel subfamily M member 2-like | 111124528 |
| NC_035782.1 | 46756600 | 46757600 | 1 | progesterone-induced-blocking factor 1-like | 111126173 |
| NC_035782.1 | 48020600 | 48022200 | 1 | complement C1q-like protein 4 | 111124282 |
| NC_035782.1 | 48168200 | 48170200 | 1 | cell adhesion molecule-related/down-regulated by oncogenes-like | 111123813 |
| NC_035782.1 | 48358600 | 48359800 | 1 | FMRFamide receptor-like | 111126791 |
| NC_035782.1 | 50111400 | 50114800 | 1 | abnormal spindle-like microcephaly-associated protein homolog | 111125042 |
| NC_035782.1 | 53597600 | 53599200 | 1 | Krueppel-like factor 8 | 111123594 |
| NC_035782.1 | 55410200 | 55411200 | 1 | LIM homeobox transcription factor 1-beta-like | 111126347 |
| NC_035782.1 | 55466600 | 55469600 | 1 | roquin-1-like | 111122645 |
| NC_035782.1 | 55469800 | 55472600 | 1 | roquin-1-like | 111122645 |
| NC_035782.1 | 58413400 | 58414600 | 1 | phospholipid scramblase 1-like | 111127106 |
| NC_035782.1 | 61203200 | 61204400 | 1 | motile sperm domain-containing protein 2-like | 111124916 |
| NC_035782.1 | 62991600 | 62992800 | 1 | cytoplasmic FMR1-interacting protein 1-like | 111125498 |
| NC_035782.1 | 72338200 | 72339400 | 1 | glycine receptor subunit alpha-2-like | 111127344 |
| NC_035782.1 | 74022000 | 74023000 | 1 | histidine protein methyltransferase 1 homolog | 111127352 |
| NC_035783.1 | 6895600 | 6897000 | 0 |  |  |
| NC_035783.1 | 7808800 | 7811200 | 1 | high affinity cAMP-specific and IBMX-insensitive 3',5'-cyclic phosphodiesterase 8B-like | 111128933 |
| NC_035783.1 | 7812200 | 7814800 | 1 | high affinity cAMP-specific and IBMX-insensitive 3',5'-cyclic phosphodiesterase 8B-like | 111128933 |
| NC_035783.1 | 8040200 | 8042000 | 1 | UPF0469 protein KIAA0907 homolog | 111129017 |
| NC_035783.1 | 8174600 | 8175800 | 1 | arrestin domain-containing protein 17-like | 111127994 |
| NC_035783.1 | 8203400 | 8204600 | 2 | uncharacterized LOC111128011;arrestin domain-containing protein 17-like | 111128011;111127994 |
| NC_035783.1 | 10592800 | 10594600 | 0 |  |  |
| NC_035783.1 | 11029400 | 11030400 | 1 | cell growth regulator with RING finger domain protein 1-like | 111130600 |
| NC_035783.1 | 11575600 | 11576600 | 1 | HHIP-like protein 2 | 111130208 |
| NC_035783.1 | 12709000 | 12710200 | 1 | uncharacterized LOC111128677 | 111128677 |
| NC_035783.1 | 13791000 | 13792000 | 1 | translation initiation factor IF-2-like | 111131638 |
| NC_035783.1 | 14341000 | 14342000 | 1 | uncharacterized LOC111130498 | 111130498 |
| NC_035783.1 | 15760400 | 15762800 | 1 | galectin-8-like | 111130470 |
| NC_035783.1 | 15903200 | 15906000 | 0 |  |  |
| NC_035783.1 | 17436400 | 17438000 | 1 | CCA tRNA nucleotidyltransferase 1, mitochondrial-like | 111131868 |
| NC_035783.1 | 18696000 | 18697000 | 0 |  |  |
| NC_035783.1 | 19968800 | 19971400 | 2 | zinc finger protein 318-like;ankyrin repeat domain-containing protein 17-like | 111131753;111131751 |
| NC_035783.1 | 21063600 | 21064800 | 1 | WD repeat-containing protein 81-like | 111128212 |
| NC_035783.1 | 22556200 | 22557200 | 1 | limbic system-associated membrane protein-like | 111131535 |
| NC_035783.1 | 22972200 | 22973400 | 1 | glycoprotein 3-alpha-L-fucosyltransferase A-like | 111128839 |
| NC_035783.1 | 23359000 | 23360400 | 2 | uncharacterized LOC111130136;neuronal acetylcholine receptor subunit alpha-10-like | 111130125;111130136 |
| NC_035783.1 | 24889600 | 24890800 | 1 | glucosylceramidase-like | 111127758 |
| NC_035783.1 | 25838400 | 25839400 | 1 | uncharacterized LOC111130338 | 111130338 |
| NC_035783.1 | 28893800 | 28895600 | 1 | calmodulin | 111129443 |
| NC_035783.1 | 28897400 | 28898400 | 1 | calmodulin | 111129443 |
| NC_035783.1 | 28899200 | 28900800 | 0 |  |  |
| NC_035783.1 | 30127800 | 30128800 | 1 | CWF19-like protein 2 | 111131012 |
| NC_035783.1 | 30129800 | 30131000 | 1 | CWF19-like protein 2 | 111131012 |
| NC_035783.1 | 32258000 | 32259200 | 1 | tetraspanin-3-like | 111128557 |
| NC_035783.1 | 35698800 | 35699800 | 1 | neuron navigator 3-like | 111128034 |
| NC_035783.1 | 35742600 | 35743800 | 1 | uncharacterized LOC111129899 | 111129899 |
| NC_035783.1 | 38221400 | 38222400 | 1 | uncharacterized LOC111131685 | 111131685 |
| NC_035783.1 | 38223200 | 38225200 | 1 | uncharacterized LOC111131685 | 111131685 |
| NC_035783.1 | 50961600 | 50964200 | 1 | sushi, von Willebrand factor type A, EGF and pentraxin domain-containing protein 1-like | 111128942 |
| NC_035783.1 | 55286000 | 55287200 | 0 |  |  |
| NC_035783.1 | 55289400 | 55293800 | 1 | angiopoietin-related protein 7-like | 111128067 |
| NC_035783.1 | 57085200 | 57086400 | 0 |  |  |
| NC_035783.1 | 58563800 | 58565800 | 1 | GRAM domain-containing protein 2B-like | 111130081 |
| NC_035783.1 | 58571000 | 58574600 | 1 | GRAM domain-containing protein 2B-like | 111130081 |
| NC_035784.1 | 935200 | 936400 | 1 | uncharacterized LOC111137177 | 111137177 |
| NC_035784.1 | 7236400 | 7237400 | 0 |  |  |
| NC_035784.1 | 8623400 | 8624400 | 0 |  |  |
| NC_035784.1 | 10219000 | 10221200 | 1 | FACT complex subunit SSRP1-like | 111137659 |
| NC_035784.1 | 13260800 | 13261800 | 1 | D-2-hydroxyglutarate dehydrogenase, mitochondrial-like | 111133078 |
| NC_035784.1 | 15041400 | 15044800 | 0 |  |  |
| NC_035784.1 | 21277200 | 21279000 | 1 | galactose-specific lectin nattectin-like | 111133656 |
| NC_035784.1 | 21285000 | 21287800 | 1 | uncharacterized protein YMR196W-like | 111137731 |
| NC_035784.1 | 24099400 | 24100800 | 1 | multidrug resistance protein 1-like | 111134479 |
| NC_035784.1 | 24189800 | 24191400 | 1 | agrin-like | 111132939 |
| NC_035784.1 | 25031600 | 25033200 | 1 | myosin heavy chain, clone 203-like | 111136891 |
| NC_035784.1 | 25661000 | 25662000 | 0 |  |  |
| NC_035784.1 | 26242600 | 26244200 | 1 | L-fucose kinase-like | 111135841 |
| NC_035784.1 | 26313200 | 26314200 | 1 | uncharacterized LOC111136861 | 111136861 |
| NC_035784.1 | 27589600 | 27591400 | 0 |  |  |
| NC_035784.1 | 28697000 | 28698000 | 1 | E3 ubiquitin-protein ligase Bre1-like | 111134830 |
| NC_035784.1 | 29271400 | 29272400 | 0 |  |  |
| NC_035784.1 | 31045400 | 31046800 | 0 |  |  |
| NC_035784.1 | 33934800 | 33936400 | 0 |  |  |
| NC_035784.1 | 34959800 | 34961600 | 1 | uncharacterized LOC111133256 | 111133256 |
| NC_035784.1 | 35552800 | 35554000 | 1 | integumentary mucin C.1-like | 111132166 |
| NC_035784.1 | 37105800 | 37107000 | 0 |  |  |
| NC_035784.1 | 37256600 | 37258000 | 1 | prostaglandin E2 receptor EP2 subtype-like | 111137929 |
| NC_035784.1 | 38188000 | 38189000 | 1 | spermidine synthase-like | 111137622 |
| NC_035784.1 | 39836000 | 39837600 | 1 | splicing factor, suppressor of white-apricot homolog | 111135697 |
| NC_035784.1 | 46820400 | 46823000 | 1 | uncharacterized LOC111136713 | 111136713 |
| NC_035784.1 | 54562400 | 54563400 | 1 | uncharacterized LOC111135261 | 111135261 |
| NC_035784.1 | 58047600 | 58048600 | 0 |  |  |
| NC_035784.1 | 59586200 | 59587200 | 1 | proton-coupled folate transporter-like | 111137149 |
| NC_035784.1 | 79425400 | 79426400 | 0 |  |  |
| NC_035784.1 | 80477800 | 80478800 | 0 |  |  |
| NC_035785.1 | 535800 | 537200 | 0 |  |  |
| NC_035785.1 | 12696600 | 12697800 | 1 | protein PAT1 homolog 1-like | 111100321 |
| NC_035785.1 | 13935400 | 13936400 | 1 | uncharacterized LOC111101184 | 111101184 |
| NC_035785.1 | 45947600 | 45949400 | 1 | diacylglycerol kinase theta-like | 111100477 |
| NC_035786.1 | 2463800 | 2466200 | 1 | major vault protein-like | 111104657 |
| NC_035786.1 | 8813000 | 8814000 | 0 |  |  |
| NC_035786.1 | 8848000 | 8849600 | 1 | ubiquinone biosynthesis protein COQ9, mitochondrial-like | 111103518 |
| NC_035786.1 | 10066400 | 10067600 | 1 | protein kinase C-binding protein NELL1-like | 111102504 |
| NC_035786.1 | 27538400 | 27540200 | 0 |  |  |
| NC_035786.1 | 29133200 | 29136400 | 2 | alpha-crystallin B chain-like;DNA helicase B-like | 111102823;111105459 |
| NC_035786.1 | 38909800 | 38910800 | 0 |  |  |
| NC_035786.1 | 42418000 | 42419800 | 0 |  |  |
| NC_035786.1 | 45931200 | 45932800 | 1 | uncharacterized LOC111102457 | 111102457 |
| NC_035786.1 | 46244600 | 46245600 | 1 | phospholipase D1-like | 111103356 |
| NC_035786.1 | 49714800 | 49716200 | 1 | 15-hydroxyprostaglandin dehydrogenase [NAD(+)]-like | 111104693 |
| NC_035786.1 | 49735400 | 49736400 | 1 | uncharacterized LOC111102431 | 111102431 |
| NC_035786.1 | 49920200 | 49921200 | 2 | coiled-coil domain-containing protein 191-like;molybdenum cofactor biosynthesis protein 1-like | 111104850;111103061 |
| NC_035786.1 | 52804600 | 52806000 | 0 |  |  |
| NC_035786.1 | 52967200 | 52969400 | 2 | uncharacterized LOC111103624;transmembrane protein 39A-like | 111103624;111103625 |
| NC_035786.1 | 52986400 | 52989000 | 0 |  |  |
| NC_035786.1 | 52990800 | 52992400 | 1 | cerebellin-2-like | 111104570 |
| NC_035786.1 | 56309400 | 56311200 | 0 |  |  |
| NC_035787.1 | 8407600 | 8408600 | 1 | transforming growth factor-beta-induced protein ig-h3-like | 111107561 |
| NC_035787.1 | 16117000 | 16118800 | 0 |  |  |
| NC_035787.1 | 37749000 | 37750200 | 1 | uncharacterized LOC111106352 | 111106352 |
| NC_035787.1 | 37752000 | 37753200 | 0 |  |  |
| NC_035787.1 | 41051200 | 41052200 | 1 | uncharacterized LOC111107659 | 111107659 |
| NC_035787.1 | 41052400 | 41053600 | 1 | uncharacterized LOC111107659 | 111107659 |
| NC_035787.1 | 52795400 | 52797200 | 1 | plasma membrane calcium-transporting ATPase 2-like | 111108086 |
| NC_035787.1 | 57697600 | 57698600 | 1 | synaptotagmin-6-like | 111108510 |
| NC_035787.1 | 64403800 | 64405000 | 1 | laminin subunit beta-1-like | 111109289 |
| NC_035787.1 | 64517800 | 64519400 | 1 | voltage-dependent calcium channel subunit alpha-2/delta-4-like | 111108275 |
| NC_035788.1 | 26614400 | 26618200 | 2 | uncharacterized LOC111113662;uncharacterized transmembrane protein DDB_G0289901-like | 111114791;111113662 |
| NC_035788.1 | 26618800 | 26620600 | 1 | uncharacterized transmembrane protein DDB_G0289901-like | 111114791 |
| NC_035788.1 | 26621400 | 26628000 | 1 | uncharacterized transmembrane protein DDB_G0289901-like | 111114791 |
| NC_035788.1 | 39040000 | 39041200 | 1 | uncharacterized LOC111113818 | 111113818 |
| NC_035788.1 | 46489400 | 46491400 | 1 | uncharacterized LOC111113211 | 111113211 |
| NC_035788.1 | 46491800 | 46492800 | 1 | uncharacterized LOC111113211 | 111113211 |
| NC_035788.1 | 52351200 | 52353400 | 0 |  |  |
| NC_035788.1 | 68422600 | 68423600 | 0 |  |  |
| NC_035788.1 | 71032000 | 71033400 | 0 |  |  |
| NC_035788.1 | 73258400 | 73260200 | 1 | uncharacterized LOC111115190 | 111115190 |
| NC_035788.1 | 78424000 | 78425000 | 0 |  |  |
| NC_035788.1 | 80341000 | 80342000 | 0 |  |  |
| NC_035788.1 | 81386800 | 81388400 | 1 | ras-related protein M-Ras-like | 111112220 |
| NC_035788.1 | 87389600 | 87390600 | 1 | acetylcholinesterase-like | 111115698 |
| NC_035788.1 | 89670200 | 89671400 | 1 | kiSS-1 receptor-like | 111111707 |
| NC_035788.1 | 89799400 | 89801400 | 1 | uncharacterized LOC111111701 | 111111701 |
| NC_035788.1 | 95348000 | 95349000 | 0 |  |  |
| NC_035789.1 | 5065600 | 5067400 | 0 |  |  |
| NC_035789.1 | 7268800 | 7270400 | 1 | uncharacterized LOC111117604 | 111117604 |
| NC_035789.1 | 8842600 | 8844000 | 1 | protogenin B-like | 111117904 |
| NC_035789.1 | 10187200 | 10188800 | 1 | uncharacterized LOC111117060 | 111117060 |
| NC_035789.1 | 22387600 | 22389800 | 0 |  |  |
| Hummock Cove (HC-VA) - Chlora’s Point (CLP) | | | | | |
| NC_035780.1 | 3290200 | 3291200 | 0 |  |  |
| NC_035780.1 | 9485200 | 9486200 | 0 |  |  |
| NC_035780.1 | 10385000 | 10388200 | 2 | D-beta-hydroxybutyrate dehydrogenase, mitochondrial-like | 111105919;111114604 |
| NC_035780.1 | 24490400 | 24492200 | 1 | ATP-dependent 6-phosphofructokinase-like | 111124867 |
| NC_035780.1 | 36228000 | 36229800 | 1 | heat shock protein 27-like | 111137387 |
| NC_035780.1 | 36230000 | 36231200 | 0 |  |  |
| NC_035780.1 | 36573600 | 36581000 | 2 | perlucin-like protein | 111119928;111119911 |
| NC_035780.1 | 36582200 | 36586600 | 2 | perlucin-like protein | 111119895;111119938 |
| NC_035780.1 | 36589200 | 36590200 | 0 |  |  |
| NC_035780.1 | 36591600 | 36595400 | 2 | perlucin-like protein;C-type mannose receptor 2-like | 111119902;111119947 |
| NC_035780.1 | 37620800 | 37621800 | 0 |  |  |
| NC_035780.1 | 38051000 | 38054200 | 1 | protein sym-1-like | 111101232 |
| NC_035780.1 | 38054600 | 38056400 | 1 | protein sym-1-like | 111101232 |
| NC_035780.1 | 38059800 | 38060800 | 1 | protein sym-1-like | 111101232 |
| NC_035780.1 | 38064000 | 38066000 | 1 | protein sym-1-like | 111101232 |
| NC_035780.1 | 38067200 | 38069200 | 1 | protein sym-1-like | 111101232 |
| NC_035780.1 | 38069600 | 38071000 | 1 | protein sym-1-like | 111101232 |
| NC_035780.1 | 38073000 | 38074600 | 1 | 60S acidic ribosomal protein P1-like | 111114718 |
| NC_035780.1 | 38075800 | 38079000 | 2 | thioredoxin domain-containing protein 5-like;60S acidic ribosomal protein P1-like | 111114718;111104099 |
| NC_035780.1 | 49760000 | 49763200 | 1 | clumping factor A-like | 111119246 |
| NC_035780.1 | 49763400 | 49764400 | 0 |  |  |
| NC_035780.1 | 54788000 | 54797000 | 1 | rac guanine nucleotide exchange factor B-like | 111102987 |
| NC_035780.1 | 54806800 | 54808600 | 0 |  |  |
| NC_035780.1 | 54810600 | 54814000 | 1 | xanthine dehydrogenase-like | 111136587 |
| NC_035780.1 | 54819400 | 54824000 | 1 | xanthine dehydrogenase-like | 111136587 |
| NC_035780.1 | 54824200 | 54827800 | 1 | xanthine dehydrogenase-like | 111136587 |
| NC_035780.1 | 54828000 | 54834000 | 2 | xanthine dehydrogenase-like;elongation factor 1-alpha | 111136587;111127225 |
| NC_035780.1 | 54835000 | 54836000 | 0 |  |  |
| NC_035780.1 | 57738600 | 57742400 | 1 | uncharacterized LOC111125477 | 111125477 |
| NC_035780.1 | 58792000 | 58793800 | 1 | UPF0676 protein C1494.01-like | 111099420 |
| NC_035780.1 | 58794400 | 58795600 | 1 | UPF0676 protein C1494.01-like | 111099420 |
| NC_035780.1 | 58807600 | 58809800 | 1 | 60S ribosomal protein L12-like | 111119071 |
| NC_035780.1 | 61125600 | 61126600 | 2 | transcription factor CP2-like;sorting nexin-10-like | 111135003;111100168 |
| NC_035780.1 | 61456600 | 61458200 | 1 | choline/ethanolamine kinase-like | 111099557 |
| NC_035781.1 | 7348600 | 7354400 | 3 | CDK5 and ABL1 enzyme substrate 2-like;Rieske domain-containing protein-like;manganese-dependent ADP-ribose/CDP-alcohol diphosphatase-like | 111117911;111117912;111118272 |
| NC_035781.1 | 7357000 | 7358400 | 1 | agrin-like | 111121060 |
| NC_035781.1 | 33596400 | 33602600 | 1 | protein phosphatase 1 regulatory subunit 7-like | 111121780 |
| NC_035781.1 | 33603200 | 33633800 | 3 | zinc finger protein 511-like;CDP-diacylglycerol--glycerol-3-phosphate 3-phosphatidyltransferase, mitochondrial-like;dynein beta chain, ciliary-like | 111118137;111120585;111121829 |
| NC_035781.1 | 48388400 | 48389400 | 1 | chitobiosyldiphosphodolichol beta-mannosyltransferase-like | 111121065 |
| NC_035781.1 | 48390600 | 48391600 | 1 | chitobiosyldiphosphodolichol beta-mannosyltransferase-like | 111121065 |
| NC_035782.1 | 18097800 | 18112800 | 3 | uncharacterized LOC111126876;uncharacterized LOC111127036;transforming growth factor-beta-induced protein ig-h3-like | 111127036;111124736;111126876 |
| NC_035782.1 | 18114600 | 18117000 | 2 | uncharacterized LOC111124738;uncharacterized LOC111124734 | 111124734;111124738 |
| NC_035782.1 | 18124000 | 18127400 | 1 | uncharacterized LOC111124738 | 111124738 |
| NC_035782.1 | 18132600 | 18134400 | 0 |  |  |
| NC_035782.1 | 18137400 | 18142000 | 1 | transforming growth factor-beta-induced protein ig-h3-like | 111124741 |
| NC_035782.1 | 18145000 | 18146200 | 0 |  |  |
| NC_035782.1 | 18149000 | 18151600 | 1 | transforming growth factor-beta-induced protein ig-h3-like | 111124739 |
| NC_035782.1 | 18182000 | 18183200 | 1 | uncharacterized LOC111124078 | 111124078 |
| NC_035782.1 | 18316800 | 18318600 | 0 |  |  |
| NC_035782.1 | 18344000 | 18345200 | 0 |  |  |
| NC_035782.1 | 21239200 | 21240200 | 0 |  |  |
| NC_035782.1 | 27688400 | 27689600 | 1 | sodium- and chloride-dependent GABA transporter 1-like | 111126767 |
| NC_035782.1 | 35913800 | 35915600 | 1 | carnosine synthase 1-like | 111125295 |
| NC_035782.1 | 38680600 | 38682200 | 1 | sodium/potassium-transporting ATPase subunit alpha-like | 111124927 |
| NC_035782.1 | 48027400 | 48028800 | 1 | myosin heavy chain, striated muscle-like | 111124621 |
| NC_035782.1 | 48032800 | 48036800 | 1 | myosin heavy chain, striated muscle-like | 111124621 |
| NC_035782.1 | 51986000 | 51987000 | 1 | uncharacterized LOC111123328 | 111123328 |
| NC_035782.1 | 53723400 | 53724400 | 1 | alpha-aspartyl dipeptidase-like | 111123988 |
| NC_035782.1 | 53725800 | 53727600 | 1 | alpha-aspartyl dipeptidase-like | 111123988 |
| NC_035782.1 | 61852600 | 61857600 | 1 | uncharacterized LOC111122921 | 111122921 |
| NC_035782.1 | 61857800 | 61863800 | 2 | uncharacterized LOC111122921;low-density lipoprotein receptor-related protein 2-like | 111122921;111122922 |
| NC_035782.1 | 61864000 | 61865400 | 1 | low-density lipoprotein receptor-related protein 2-like | 111122922 |
| NC_035782.1 | 61874800 | 61877000 | 1 | low-density lipoprotein receptor-related protein 2-like | 111122922 |
| NC_035782.1 | 61878000 | 61880600 | 1 | low-density lipoprotein receptor-related protein 2-like | 111122922 |
| NC_035782.1 | 68027800 | 68029000 | 0 |  |  |
| NC_035782.1 | 71505200 | 71506200 | 1 | POU domain, class 4, transcription factor 3-like | 111126745 |
| NC_035783.1 | 10280600 | 10287400 | 2 | ectonucleoside triphosphate diphosphohydrolase 5-like;basal body-orientation factor 1-like | 111129901;111129900 |
| NC_035783.1 | 10291200 | 10292200 | 1 | ectonucleoside triphosphate diphosphohydrolase 5-like | 111129901 |
| NC_035783.1 | 10305000 | 10306600 | 0 |  |  |
| NC_035783.1 | 10308800 | 10309800 | 1 | nuclear receptor ROR-beta-like | 111129823 |
| NC_035783.1 | 10313800 | 10315000 | 1 | nuclear receptor ROR-beta-like | 111129823 |
| NC_035783.1 | 11598400 | 11599400 | 1 | integrator complex subunit 7-like | 111128756 |
| NC_035783.1 | 16644200 | 16647200 | 1 | hornerin-like | 111128845 |
| NC_035783.1 | 16648000 | 16650600 | 1 | hornerin-like | 111128845 |
| NC_035784.1 | 5596800 | 5597800 | 1 | kynurenine formamidase-like | 111133392 |
| NC_035784.1 | 5606400 | 5607400 | 0 |  |  |
| NC_035784.1 | 7643400 | 7644800 | 2 | coiled-coil domain-containing protein 189-like;uncharacterized LOC111136911 | 111136911;111137967 |
| NC_035784.1 | 7648200 | 7649200 | 2 | coiled-coil domain-containing protein 189-like;uncharacterized LOC111136911 | 111136911;111137967 |
| NC_035784.1 | 7651800 | 7653200 | 1 | uncharacterized LOC111136911 | 111136911 |
| NC_035784.1 | 7653600 | 7655200 | 1 | uncharacterized LOC111136911 | 111136911 |
| NC_035784.1 | 8355600 | 8357400 | 1 | uncharacterized LOC111137029 | 111137029 |
| NC_035784.1 | 8360800 | 8362400 | 1 | uncharacterized LOC111137029 | 111137029 |
| NC_035784.1 | 8363000 | 8364000 | 1 | uncharacterized LOC111137029 | 111137029 |
| NC_035784.1 | 8364200 | 8368800 | 1 | uncharacterized LOC111137029 | 111137029 |
| NC_035784.1 | 8370000 | 8371000 | 1 | uncharacterized LOC111137029 | 111137029 |
| NC_035784.1 | 13281400 | 13284000 | 1 | E3 ubiquitin-protein ligase E3D-like | 111137290 |
| NC_035784.1 | 13293000 | 13296600 | 1 | chloride transport protein 6-like | 111099166 |
| NC_035784.1 | 28331400 | 28332400 | 0 |  |  |
| NC_035784.1 | 28332600 | 28336000 | 1 | submaxillary gland androgen-regulated protein 3B-like | 111099173 |
| NC_035784.1 | 36512400 | 36514800 | 1 | CD109 antigen-like | 111133801 |
| NC_035784.1 | 36517000 | 36518800 | 1 | CD109 antigen-like | 111133801 |
| NC_035784.1 | 36828200 | 36831800 | 1 | uncharacterized LOC111135484 | 111135484 |
| NC_035784.1 | 48883600 | 48890200 | 1 | uncharacterized LOC111134463 | 111134463 |
| NC_035784.1 | 48890400 | 48891800 | 0 |  |  |
| NC_035784.1 | 48920400 | 48922600 | 0 |  |  |
| NC_035784.1 | 48922800 | 48925000 | 0 |  |  |
| NC_035784.1 | 48925400 | 48926400 | 0 |  |  |
| NC_035784.1 | 50789000 | 50790000 | 0 |  |  |
| NC_035784.1 | 57455000 | 57456000 | 1 | zinc finger protein 83-like | 111135642 |
| NC_035784.1 | 61768600 | 61770400 | 2 | uncharacterized LOC111136680;S-formylglutathione hydrolase-like | 111136680;111136681 |
| NC_035784.1 | 61997600 | 61999800 | 1 | uncharacterized LOC111132044 | 111132044 |
| NC_035784.1 | 62000400 | 62001400 | 1 | uncharacterized LOC111132044 | 111132044 |
| NC_035784.1 | 62001800 | 62004800 | 1 | uncharacterized LOC111132044 | 111132044 |
| NC_035784.1 | 62009000 | 62011800 | 0 |  |  |
| NC_035784.1 | 62012200 | 62014200 | 1 | cGMP-specific 3',5'-cyclic phosphodiesterase-like | 111134772 |
| NC_035784.1 | 62141600 | 62143400 | 1 | nucleolysin TIAR-like | 111136733 |
| NC_035784.1 | 92997400 | 92999400 | 0 |  |  |
| NC_035785.1 | 43954000 | 43955000 | 0 |  |  |
| NC_035786.1 | 1668000 | 1669200 | 1 | negative elongation factor D-like | 111102475 |
| NC_035787.1 | 6985400 | 6986600 | 0 |  |  |
| NC_035787.1 | 60708400 | 60709800 | 1 | plexin-B-like | 111108439 |
| NC_035787.1 | 62830400 | 62831600 | 1 | dedicator of cytokinesis protein 3-like | 111108426 |
| NC_035788.1 | 21691400 | 21694600 | 1 | ceramide synthase 5-like | 111111442 |
| NC_035788.1 | 46477400 | 46478400 | 0 |  |  |
| NC_035788.1 | 78436800 | 78438200 | 0 |  |  |
| NC_035789.1 | 3726600 | 3727600 | 0 |  |  |
| NC_035789.1 | 3744400 | 3747400 | 1 | spidroin-1-like | 111117433 |
| NC_035789.1 | 8163800 | 8165600 | 1 | uncharacterized LOC111117490 | 111117490 |
| Hog Island (HI) - Sherman Marsh (SM) | | | | | |
| NC_035780.1 | 13372200 | 13373200 | 0 |  |  |
| NC_035780.1 | 13425000 | 13426000 | 1 | amyloid beta A4 protein-like | 111136389 |
| NC_035780.1 | 15391800 | 15393400 | 1 | MAM and LDL-receptor class A domain-containing protein 1-like | 111099722 |
| NC_035780.1 | 15395800 | 15403600 | 1 | MAM and LDL-receptor class A domain-containing protein 1-like | 111099722 |
| NC_035780.1 | 15407200 | 15416400 | 1 | MAM and LDL-receptor class A domain-containing protein 1-like | 111099722 |
| NC_035780.1 | 15420400 | 15426800 | 1 | MAM and LDL-receptor class A domain-containing protein 1-like | 111099722 |
| NC_035780.1 | 15478000 | 15479200 | 0 |  |  |
| NC_035780.1 | 27097200 | 27099000 | 1 | temptin-like | 111099049 |
| NC_035780.1 | 27099600 | 27100800 | 1 | temptin-like | 111102570 |
| NC_035780.1 | 36630000 | 36631000 | 0 |  |  |
| NC_035780.1 | 36644600 | 36650600 | 0 |  |  |
| NC_035780.1 | 36651200 | 36653800 | 0 |  |  |
| NC_035780.1 | 59451200 | 59452200 | 1 | glutaredoxin-2, mitochondrial-like | 111128742 |
| NC_035781.1 | 26084600 | 26085800 | 1 | high-affinity choline transporter 1-like | 111121993 |
| NC_035781.1 | 26963600 | 26964600 | 1 | cell migration-inducing and hyaluronan-binding protein-like | 111122089 |
| NC_035781.1 | 27461400 | 27463000 | 1 | A disintegrin and metalloproteinase with thrombospondin motifs 2-like | 111122374 |
| NC_035781.1 | 27464000 | 27465800 | 1 | A disintegrin and metalloproteinase with thrombospondin motifs 2-like | 111122374 |
| NC_035781.1 | 29909000 | 29912800 | 1 | putative uncharacterized protein DDB_G0271606 | 111121853 |
| NC_035781.1 | 29915200 | 29918000 | 1 | putative uncharacterized protein DDB_G0271606 | 111121853 |
| NC_035781.1 | 43309200 | 43310400 | 0 |  |  |
| NC_035781.1 | 48035400 | 48036600 | 0 |  |  |
| NC_035781.1 | 48037400 | 48040600 | 0 |  |  |
| NC_035781.1 | 50139400 | 50140600 | 0 |  |  |
| NC_035781.1 | 56967200 | 56969200 | 1 | uncharacterized LOC111119968 | 111119968 |
| NC_035782.1 | 18388600 | 18390400 | 1 | uncharacterized LOC111123479 | 111123479 |
| NC_035782.1 | 32374000 | 32375800 | 1 | innexin unc-9-like | 111124985 |
| NC_035782.1 | 38304000 | 38305200 | 1 | probable phosphorylase b kinase regulatory subunit alpha | 111123775 |
| NC_035782.1 | 39438000 | 39440000 | 1 | E3 ubiquitin-protein ligase CBL-like | 111127555 |
| NC_035782.1 | 43700800 | 43702600 | 1 | uncharacterized LOC111123502 | 111123502 |
| NC_035782.1 | 43705800 | 43706800 | 1 | uncharacterized LOC111123502 | 111123502 |
| NC_035782.1 | 43710400 | 43712200 | 0 |  |  |
| NC_035782.1 | 43715800 | 43716800 | 1 | uncharacterized LOC111123503 | 111123503 |
| NC_035782.1 | 43721000 | 43727600 | 1 | uncharacterized LOC111123503 | 111123503 |
| NC_035782.1 | 46064400 | 46066200 | 1 | barH-like 1 homeobox protein | 111126306 |
| NC_035782.1 | 46067800 | 46068800 | 0 |  |  |
| NC_035782.1 | 46077400 | 46078800 | 0 |  |  |
| NC_035782.1 | 46087400 | 46088600 | 1 | barH-like 1 homeobox protein | 111123774 |
| NC_035782.1 | 46092600 | 46093600 | 1 | barH-like 1 homeobox protein | 111123774 |
| NC_035782.1 | 46099600 | 46101800 | 1 | sulfiredoxin-1-like | 111126190 |
| NC_035782.1 | 49576600 | 49577600 | 1 | CAP-Gly domain-containing linker protein 1-like | 111123248 |
| NC_035782.1 | 54968200 | 54973000 | 1 | bestrophin homolog 15-like | 111127339 |
| NC_035782.1 | 55043400 | 55044800 | 1 | succinate dehydrogenase cytochrome b560 subunit, mitochondrial-like | 111127095 |
| NC_035782.1 | 55052000 | 55055600 | 1 | cyclin-dependent kinases regulatory subunit 1-like | 111127096 |
| NC_035782.1 | 55097200 | 55098200 | 0 |  |  |
| NC_035782.1 | 61040400 | 61041400 | 1 | protein inturned-like | 111126123 |
| NC_035783.1 | 2413400 | 2416800 | 1 | neuronal acetylcholine receptor subunit alpha-10-like | 111128920 |
| NC_035783.1 | 2983400 | 2985000 | 0 |  |  |
| NC_035783.1 | 7460000 | 7461200 | 1 | ADP-ribosylation factor-like protein 15 | 111131450 |
| NC_035783.1 | 7465200 | 7466200 | 1 | phosphorylated adapter RNA export protein-like | 111131449 |
| NC_035783.1 | 8006000 | 8007600 | 1 | sodium channel protein 1 brain-like | 111128509 |
| NC_035783.1 | 8009800 | 8010800 | 1 | sodium channel protein 1 brain-like | 111128509 |
| NC_035783.1 | 9366400 | 9368600 | 1 | forkhead box protein N3-like | 111129212 |
| NC_035783.1 | 9633200 | 9634800 | 1 | uncharacterized LOC111128129 | 111128129 |
| NC_035783.1 | 10366600 | 10368000 | 1 | patatin-like phospholipase domain-containing protein 7 | 111130102 |
| NC_035783.1 | 10388200 | 10389200 | 1 | maleylacetoacetate isomerase-like | 111129960 |
| NC_035783.1 | 10389600 | 10391000 | 0 |  |  |
| NC_035783.1 | 10400800 | 10404600 | 1 | protein O-mannosyl-transferase 2-like | 111129961 |
| NC_035783.1 | 10417000 | 10418800 | 0 |  |  |
| NC_035783.1 | 10465000 | 10466000 | 1 | beta-1,4-galactosyltransferase 2-like | 111131415 |
| NC_035783.1 | 10519400 | 10521000 | 0 |  |  |
| NC_035783.1 | 14436800 | 14438200 | 1 | TELO2-interacting protein 2-like | 111128215 |
| NC_035783.1 | 14440000 | 14441600 | 1 | E3 ubiquitin-protein ligase RNF115-like | 111130274 |
| NC_035783.1 | 14445600 | 14446800 | 1 | E3 ubiquitin-protein ligase RNF115-like | 111130274 |
| NC_035783.1 | 14448400 | 14449600 | 1 | AP-5 complex subunit beta-1-like | 111131470 |
| NC_035783.1 | 14458400 | 14459400 | 1 | AP-5 complex subunit beta-1-like | 111131470 |
| NC_035783.1 | 14459600 | 14460600 | 1 | AP-5 complex subunit beta-1-like | 111131470 |
| NC_035783.1 | 14477800 | 14482200 | 1 | AP-5 complex subunit beta-1-like | 111131470 |
| NC_035783.1 | 16652800 | 16654400 | 1 | uncharacterized LOC111129113 | 111129113 |
| NC_035783.1 | 16999000 | 17000000 | 1 | uncharacterized LOC111131263 | 111131263 |
| NC_035783.1 | 17082200 | 17095000 | 2 | thyrostimulin beta-5 subunit-like;thyrostimulin alpha-2 subunit-like | 111128799;111130827 |
| NC_035783.1 | 17095400 | 17097200 | 1 | thyrostimulin alpha-2 subunit-like | 111130827 |
| NC_035783.1 | 17101800 | 17102800 | 0 |  |  |
| NC_035783.1 | 17134000 | 17136600 | 0 |  |  |
| NC_035783.1 | 17148400 | 17149400 | 0 |  |  |
| NC_035783.1 | 17160800 | 17162000 | 1 | homeobox protein otx5-B-like | 111128600 |
| NC_035783.1 | 17162200 | 17165000 | 1 | homeobox protein otx5-B-like | 111128600 |
| NC_035783.1 | 17165800 | 17168400 | 0 |  |  |
| NC_035783.1 | 19833600 | 19834600 | 1 | complex I intermediate-associated protein 30, mitochondrial-like | 111130179 |
| NC_035783.1 | 19834800 | 19839800 | 2 | complex I intermediate-associated protein 30, mitochondrial-like;calcineurin B homologous protein 1-like | 111130180;111130179 |
| NC_035783.1 | 19918200 | 19919400 | 1 | E3 ubiquitin-protein ligase HECTD1-like | 111127862 |
| NC_035783.1 | 28303800 | 28307000 | 1 | sodium/calcium exchanger 2-like | 111129088 |
| NC_035783.1 | 31664000 | 31665000 | 0 |  |  |
| NC_035783.1 | 31671400 | 31672400 | 1 | integrator complex subunit 14-like | 111131786 |
| NC_035783.1 | 31673000 | 31675600 | 1 | integrator complex subunit 14-like | 111131786 |
| NC_035783.1 | 31678400 | 31681000 | 1 | integrator complex subunit 14-like | 111131786 |
| NC_035783.1 | 31681200 | 31683000 | 1 | integrator complex subunit 14-like | 111131786 |
| NC_035783.1 | 33386800 | 33387800 | 1 | uncharacterized LOC111128416 | 111128416 |
| NC_035783.1 | 36406000 | 36407000 | 1 | aprataxin and PNK-like factor | 111129239 |
| NC_035783.1 | 37538200 | 37541000 | 1 | mirror-image polydactyly gene 1 protein-like | 111129210 |
| NC_035783.1 | 37630000 | 37631800 | 1 | tubulin polyglutamylase TTLL5-like | 111129255 |
| NC_035783.1 | 39621800 | 39622800 | 1 | uncharacterized LOC111128771 | 111128771 |
| NC_035783.1 | 43251200 | 43252400 | 1 | laccase-25-like | 111127771 |
| NC_035783.1 | 43276600 | 43279800 | 1 | allene oxide synthase-lipoxygenase protein-like | 111128395 |
| NC_035783.1 | 43281400 | 43283400 | 1 | allene oxide synthase-lipoxygenase protein-like | 111128395 |
| NC_035783.1 | 43290200 | 43293600 | 1 | allene oxide synthase-lipoxygenase protein-like | 111128395 |
| NC_035783.1 | 43423400 | 43424400 | 1 | sodium channel protein type 4 subunit alpha B-like | 111131178 |
| NC_035783.1 | 43427600 | 43428600 | 1 | sodium channel protein type 4 subunit alpha B-like | 111131178 |
| NC_035783.1 | 43429000 | 43430000 | 1 | sodium channel protein type 4 subunit alpha B-like | 111131178 |
| NC_035783.1 | 43446600 | 43447800 | 1 | sodium channel protein type 4 subunit alpha B-like | 111131178 |
| NC_035783.1 | 43448200 | 43450000 | 1 | sodium channel protein type 4 subunit alpha B-like | 111131178 |
| NC_035783.1 | 43478200 | 43479400 | 1 | homeobox protein goosecoid-like | 111127876 |
| NC_035783.1 | 45315600 | 45317200 | 1 | usherin-like | 111129694 |
| NC_035783.1 | 45355400 | 45358600 | 1 | usherin-like | 111129694 |
| NC_035783.1 | 45369800 | 45371000 | 1 | usherin-like | 111129694 |
| NC_035783.1 | 45371600 | 45372600 | 1 | usherin-like | 111129694 |
| NC_035783.1 | 45380400 | 45381400 | 1 | usherin-like | 111129694 |
| NC_035783.1 | 45381800 | 45382800 | 1 | usherin-like | 111129694 |
| NC_035783.1 | 45386800 | 45393600 | 1 | usherin-like | 111129694 |
| NC_035783.1 | 45423200 | 45424400 | 1 | suppressor of cytokine signaling 5-like | 111129019 |
| NC_035783.1 | 45432200 | 45434000 | 2 | suppressor of cytokine signaling 5-like;WD repeat and HMG-box DNA-binding protein 1-like | 111129021;111129019 |
| NC_035783.1 | 45436600 | 45438400 | 1 | WD repeat and HMG-box DNA-binding protein 1-like | 111129021 |
| NC_035783.1 | 45440600 | 45442000 | 1 | WD repeat and HMG-box DNA-binding protein 1-like | 111129021 |
| NC_035783.1 | 45442200 | 45445200 | 1 | WD repeat and HMG-box DNA-binding protein 1-like | 111129021 |
| NC_035783.1 | 45548400 | 45549400 | 1 | DNA polymerase delta subunit 2-like | 111130346 |
| NC_035783.1 | 45576600 | 45577600 | 1 | uncharacterized LOC111131636 | 111131636 |
| NC_035783.1 | 51616000 | 51617000 | 0 |  |  |
| NC_035783.1 | 51617200 | 51618800 | 0 |  |  |
| NC_035783.1 | 51707400 | 51708600 | 1 | uncharacterized LOC111128730 | 111128730 |
| NC_035783.1 | 57995400 | 57996600 | 1 | insulin gene enhancer protein isl-1-like | 111130009 |
| NC_035784.1 | 1519800 | 1521600 | 1 | rap1 GTPase-activating protein 1-like | 111133403 |
| NC_035784.1 | 2169600 | 2171200 | 0 |  |  |
| NC_035784.1 | 6878800 | 6880800 | 1 | CCR4-NOT transcription complex subunit 1-like | 111134304 |
| NC_035784.1 | 8296400 | 8297400 | 1 | uncharacterized LOC111134622 | 111134622 |
| NC_035784.1 | 8297800 | 8300600 | 1 | uncharacterized LOC111134622 | 111134622 |
| NC_035784.1 | 8325800 | 8329800 | 1 | aspartate--tRNA ligase, mitochondrial-like | 111135543 |
| NC_035784.1 | 8334400 | 8336200 | 1 | proline-serine-threonine phosphatase-interacting protein 2-like | 111135544 |
| NC_035784.1 | 8349600 | 8352000 | 2 | uncharacterized LOC111137029;uncharacterized LOC111133767 | 111133767;111137029 |
| NC_035784.1 | 11910800 | 11912600 | 2 | dCTP pyrophosphatase 1-like;uncharacterized LOC111135486 | 111135490;111135486 |
| NC_035784.1 | 11918200 | 11925600 | 3 | dCTP pyrophosphatase 1-like;uncharacterized LOC111135488;ileal sodium/bile acid cotransporter-like | 111135490;111135488;111135487 |
| NC_035784.1 | 11927400 | 11929000 | 0 |  |  |
| NC_035784.1 | 11955200 | 11956600 | 1 | uncharacterized LOC111133506 | 111133506 |
| NC_035784.1 | 13223200 | 13224200 | 2 | eukaryotic translation initiation factor 4 gamma 1-like;lysosomal amino acid transporter 1 homolog | 111133022;111133023 |
| NC_035784.1 | 13338600 | 13339600 | 1 | chordin-like | 111132884 |
| NC_035784.1 | 13342200 | 13344000 | 0 |  |  |
| NC_035784.1 | 13346800 | 13348600 | 0 |  |  |
| NC_035784.1 | 13991800 | 13993000 | 0 |  |  |
| NC_035784.1 | 14060800 | 14061800 | 1 | uncharacterized LOC111133910 | 111133910 |
| NC_035784.1 | 14068200 | 14070800 | 1 | bactericidal permeability-increasing protein-like | 111133911 |
| NC_035784.1 | 14787600 | 14788600 | 1 | uncharacterized LOC111136155 | 111136155 |
| NC_035784.1 | 16572800 | 16574000 | 1 | aspartate beta-hydroxylase domain-containing protein 2-like | 111134889 |
| NC_035784.1 | 16574400 | 16575400 | 1 | aspartate beta-hydroxylase domain-containing protein 2-like | 111134889 |
| NC_035784.1 | 16594600 | 16595800 | 1 | dynein heavy chain 10, axonemal-like | 111134888 |
| NC_035784.1 | 16597000 | 16598400 | 1 | dynein heavy chain 10, axonemal-like | 111134888 |
| NC_035784.1 | 16599200 | 16600600 | 1 | dynein heavy chain 10, axonemal-like | 111134888 |
| NC_035784.1 | 16673800 | 16674800 | 1 | uncharacterized LOC111133063 | 111133063 |
| NC_035784.1 | 27533200 | 27534200 | 0 |  |  |
| NC_035784.1 | 28395600 | 28396800 | 0 |  |  |
| NC_035784.1 | 39654600 | 39655800 | 1 | WD repeat and SOCS box-containing protein 1-like | 111137710 |
| NC_035784.1 | 44693400 | 44695000 | 1 | inactive pancreatic lipase-related protein 1-like | 111134941 |
| NC_035784.1 | 44700200 | 44701800 | 0 |  |  |
| NC_035784.1 | 44702000 | 44705000 | 1 | tyrosine-protein phosphatase non-receptor type 2-like | 111134940 |
| NC_035784.1 | 44706400 | 44707800 | 1 | tyrosine-protein phosphatase non-receptor type 2-like | 111134940 |
| NC_035784.1 | 44716600 | 44717600 | 1 | tyrosine-protein phosphatase non-receptor type 2-like | 111134940 |
| NC_035784.1 | 61681400 | 61683000 | 1 | cytochrome P450 2C14-like | 111137423 |
| NC_035784.1 | 61693200 | 61694200 | 1 | eukaryotic translation initiation factor 2A-like | 111133033 |
| NC_035784.1 | 61694600 | 61695800 | 1 | eukaryotic translation initiation factor 2A-like | 111133033 |
| NC_035784.1 | 83240400 | 83241400 | 1 | muscle M-line assembly protein unc-89-like | 111137832 |
| NC_035784.1 | 83242000 | 83243000 | 1 | muscle M-line assembly protein unc-89-like | 111137832 |
| NC_035784.1 | 92295800 | 92296800 | 1 | uncharacterized LOC111133007 | 111133007 |
| NC_035785.1 | 43980800 | 43981800 | 1 | neuropeptide Y receptor type 2-like | 111101387 |
| NC_035785.1 | 43983600 | 43984800 | 1 | neuropeptide Y receptor type 2-like | 111101387 |
| NC_035786.1 | 48974400 | 48976800 | 0 |  |  |
| NC_035787.1 | 6741800 | 6743200 | 0 |  |  |
| NC_035787.1 | 55068000 | 55069400 | 1 | uncharacterized LOC111110399 | 111110399 |
| NC_035788.1 | 689000 | 690000 | 1 | potassium voltage-gated channel subfamily H member 6-like | 111114644 |
| NC_035788.1 | 690400 | 691800 | 1 | potassium voltage-gated channel subfamily H member 6-like | 111114644 |
| NC_035788.1 | 15202200 | 15204000 | 0 |  |  |
| NC_035788.1 | 52285400 | 52286400 | 0 |  |  |
| NC_035788.1 | 81959400 | 81960600 | 1 | uncharacterized LOC111112049 | 111112049 |
| NC_035788.1 | 90980000 | 90981800 | 0 |  |  |
| NC_035789.1 | 5051200 | 5053000 | 0 |  |  |
| NC_035789.1 | 5116400 | 5118200 | 1 | uncharacterized LOC111116216 | 111116216 |
| NC_035789.1 | 22661400 | 22662400 | 1 | dihydrofolate reductase-like | 111117281 |
| Sharing between HC-CS and HC-CA - CLP | | | | | |
| NC_035781.1 | 33597600 | 33600800 | 1 | protein phosphatase 1 regulatory subunit 7-like | 111121780 |
| NC_035781.1 | 33608600 | 33610600 | 2 | dynein beta chain, ciliary-like;CDP-diacylglycerol--glycerol-3-phosphate 3-phosphatidyltransferase, mitochondrial-like | 111120585;111118137 |
| NC_035781.1 | 33610800 | 33612000 | 1 | dynein beta chain, ciliary-like | 111120585 |
| NC_035781.1 | 33612200 | 33613600 | 1 | dynein beta chain, ciliary-like | 111120585 |
| NC_035781.1 | 33614000 | 33619400 | 1 | dynein beta chain, ciliary-like | 111120585 |
| NC_035781.1 | 33619800 | 33624000 | 1 | dynein beta chain, ciliary-like | 111120585 |
| NC_035781.1 | 33624600 | 33629400 | 1 | dynein beta chain, ciliary-like | 111120585 |

### Supplemental Figures

#### Supplemental Figure 1. Graphical history of selection lines.

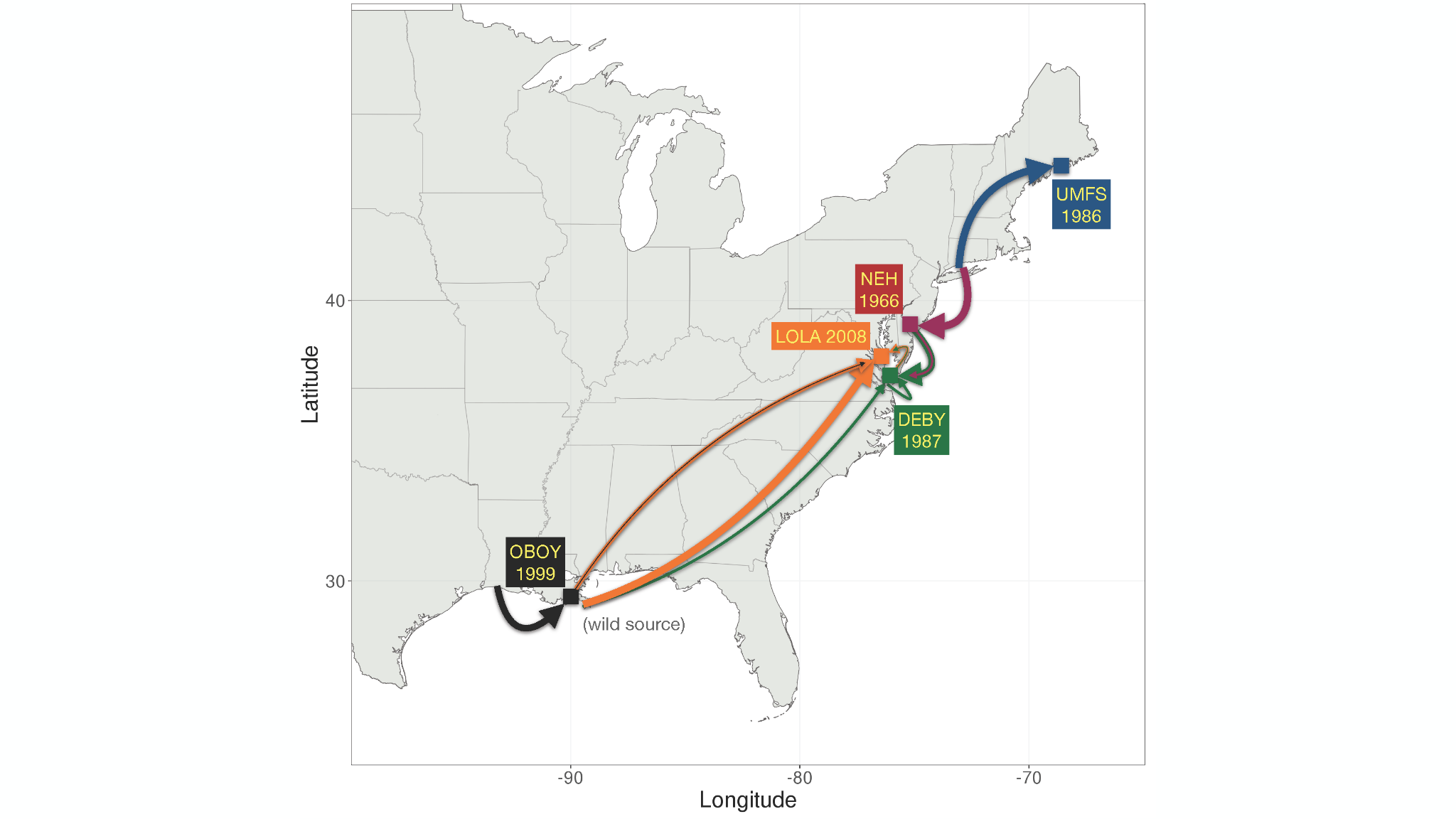

#### Supplemental Figure S2. PCA

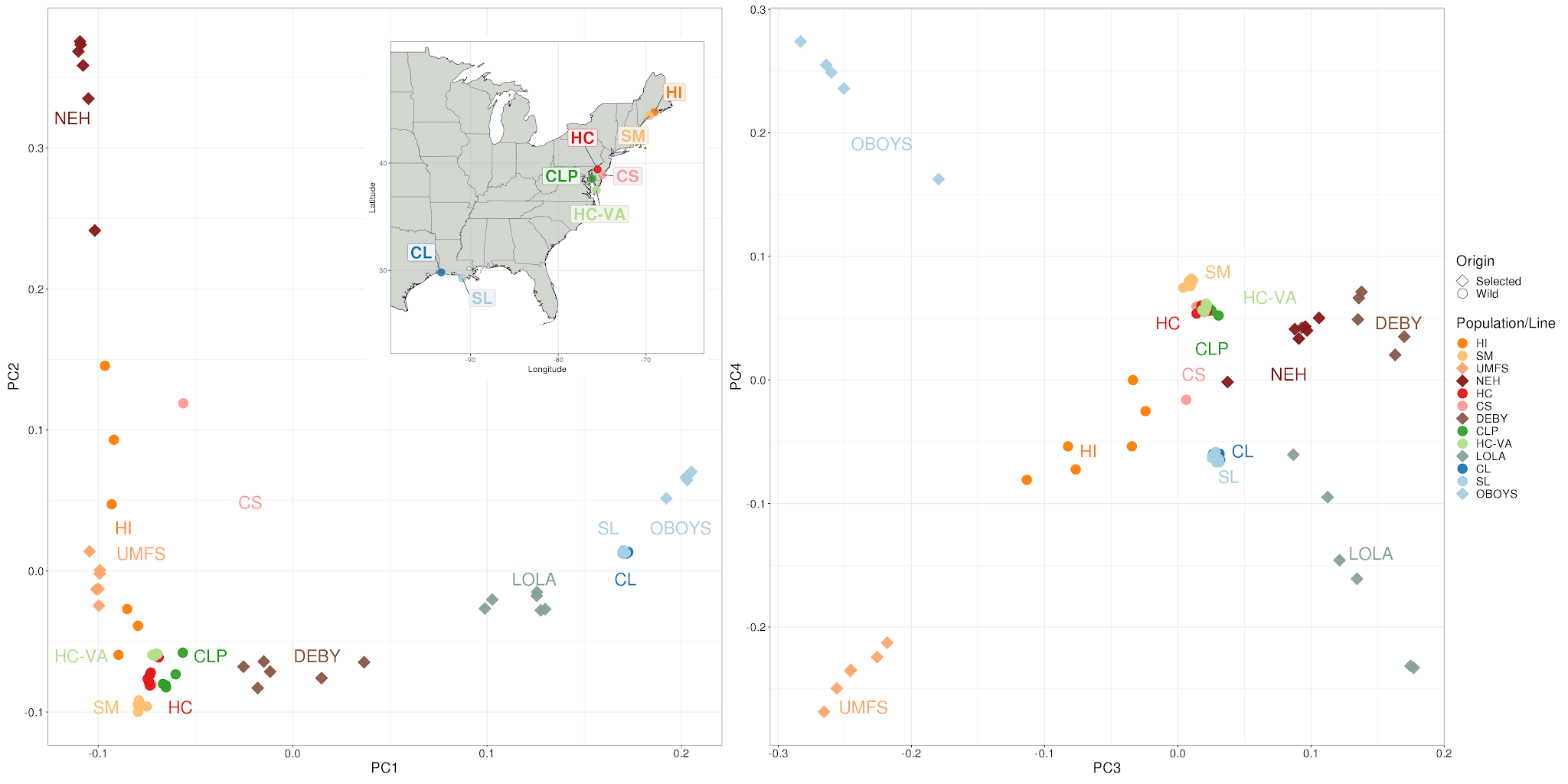

**Graphical representation of overall population structure of the eastern oyster.** Panel A is a PCA of wild populations and selected lines along with a map of wild oyster sampling locations (PC 1 and PC2). Panel B is PC 3 and PC 4. Both PCAs used all SNPs after LD clumping and outlier removal using bigsnpr (Privé et al. 2018). [Full resolution figure.](https://github.com/The-Eastern-Oyster-Genome-Project/2021_Genome_Submissions/raw/main/Population_Genomics/Output/Figures/Supplemental/Figure.S2.PCAs.png)

#### Supplemental Figure S3. Linkage Disequilibrium across chromosomes and data subsets. Wild populations on left, selected strains on right. [Full Resolution Panel 1](https://github.com/The-Eastern-Oyster-Genome-Project/2021_Genome_Submissions/blob/main/Population_Genomics/Output/Figure.S3A.AtlanticEastern.png) and [Full Resolution Panel 2](https://github.com/The-Eastern-Oyster-Genome-Project/2021_Genome_Submissions/blob/main/Population_Genomics/Output/Figure.S3B.selected.png)

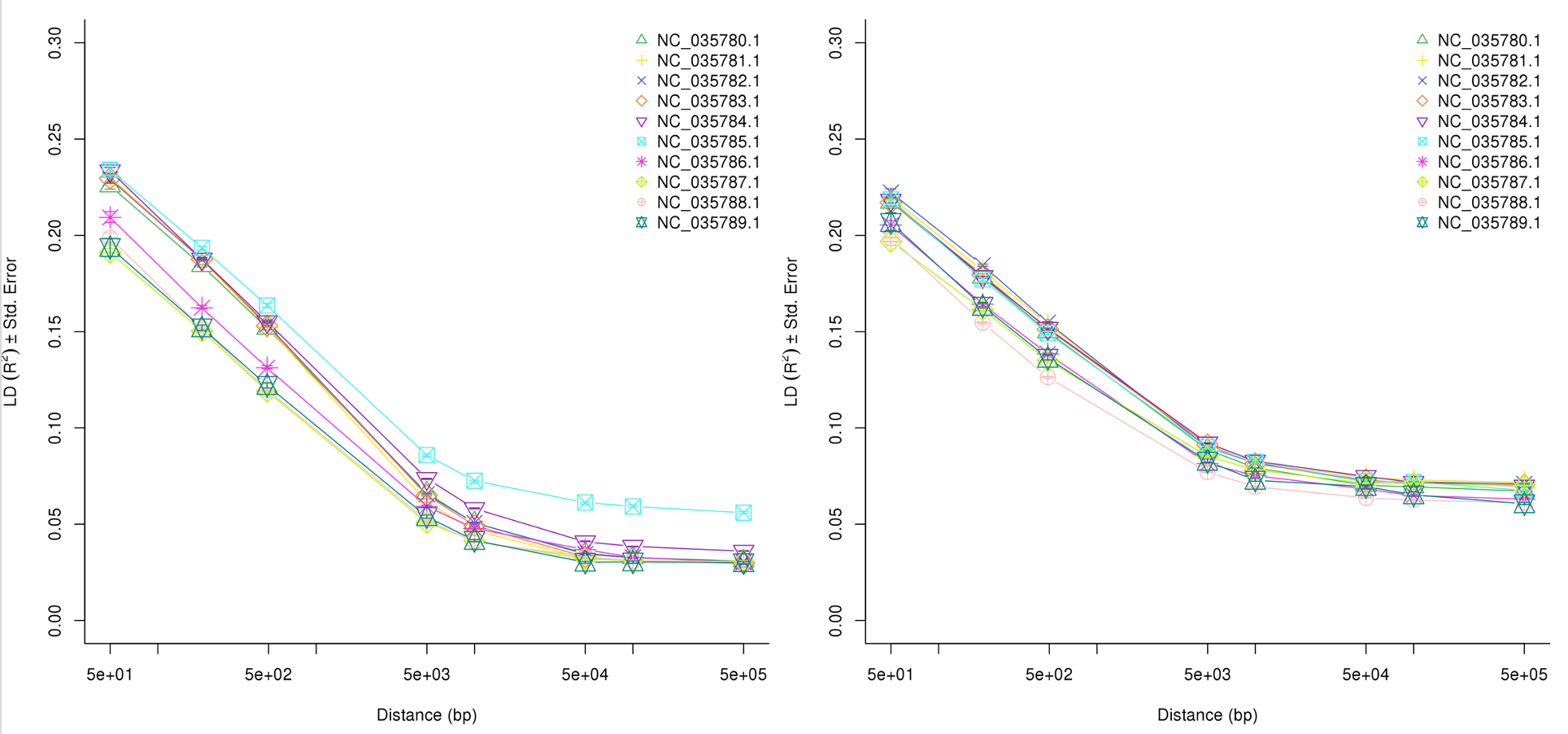

###### Figure S4- Comparison of heterozygosity across wild and selected samples

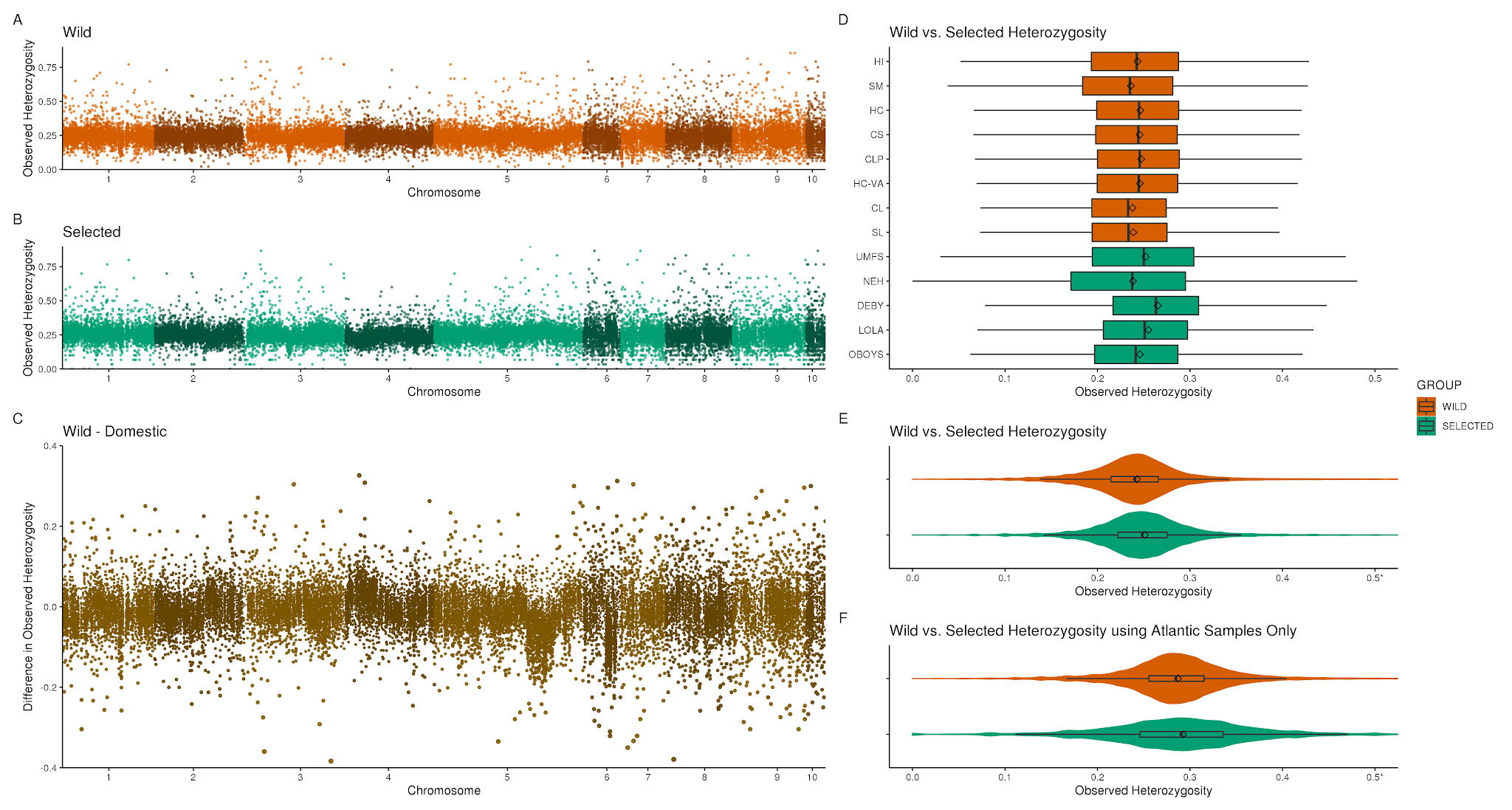
<https://raw.githubusercontent.com/The-Eastern-Oyster-Genome-Project/2021_Genome_Submissions/main/Population_Genomics/Output/Figures/Supplemental/Figure.S4.Heterozygosity.Comparison.W.S.png>**Figure S4. Comparison of heterozygosity across wild populations and selected lines.**

Panel (A) is values averaged across all wild populations in 10 kb windows across the entire genome. Panel (B) is values averaged across all selected populations in 10 kb windows across the entire genome. Panel (C) is the difference between wild and selected in 10 kb windows across the entire genome. Panel (D) is a boxplot of 10kb averaged values broken down by population and line. Panel (E) is a violin and boxplot of 10kb averaged values broken by group. Panel (F) is identical to Panel (E), except it uses only loci that are variable within Atlantic selected and wild populations.

###### Figure S5- Comparison of Tajima's *D* across wild and selected samples

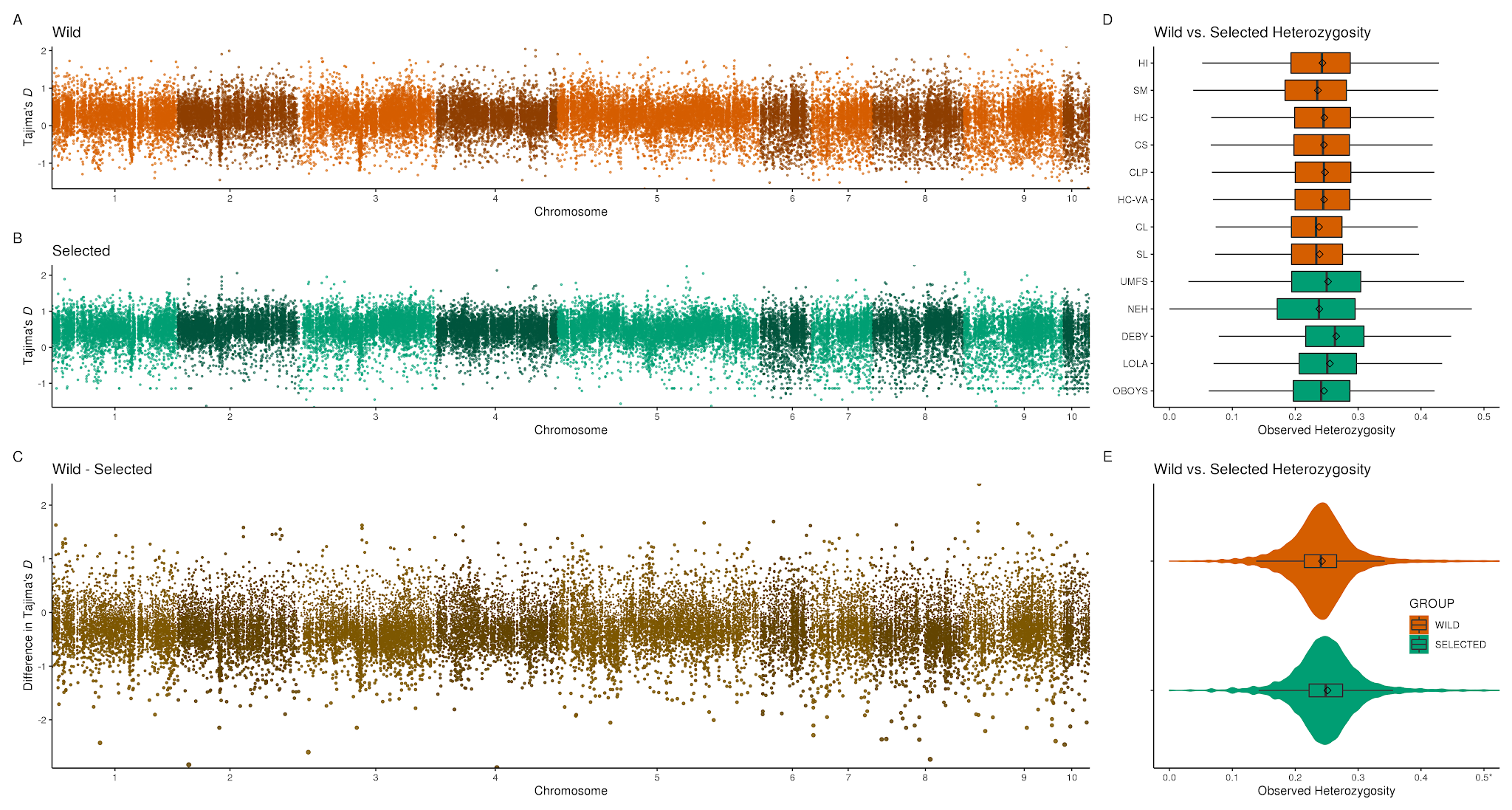
<https://github.com/The-Eastern-Oyster-Genome-Project/2021_Genome_Submissions/raw/main/Population_Genomics/Output/Figures/Supplemental/Figure.S5.TajimasD.Comparison.W.S.png>**Figure S5. Comparison of *Tajima's D* across wild populations and selected lines.**

Panel (A) is values averaged across all wild populations in 10 kb windows across the entire genome. Panel (B) is values averaged across all selected populations in 10 kb windows across the entire genome. Panel (C) is the difference between wild and selected in 10 kb windows across the entire genome. Panel (D) is a boxplot of 10kb averaged values broken down by population and line. Panel (E) is a violin and boxplot of 10kb averaged values broken by group

###### Figure S6- Comparison of copy number allelic richness across wild and selected samples

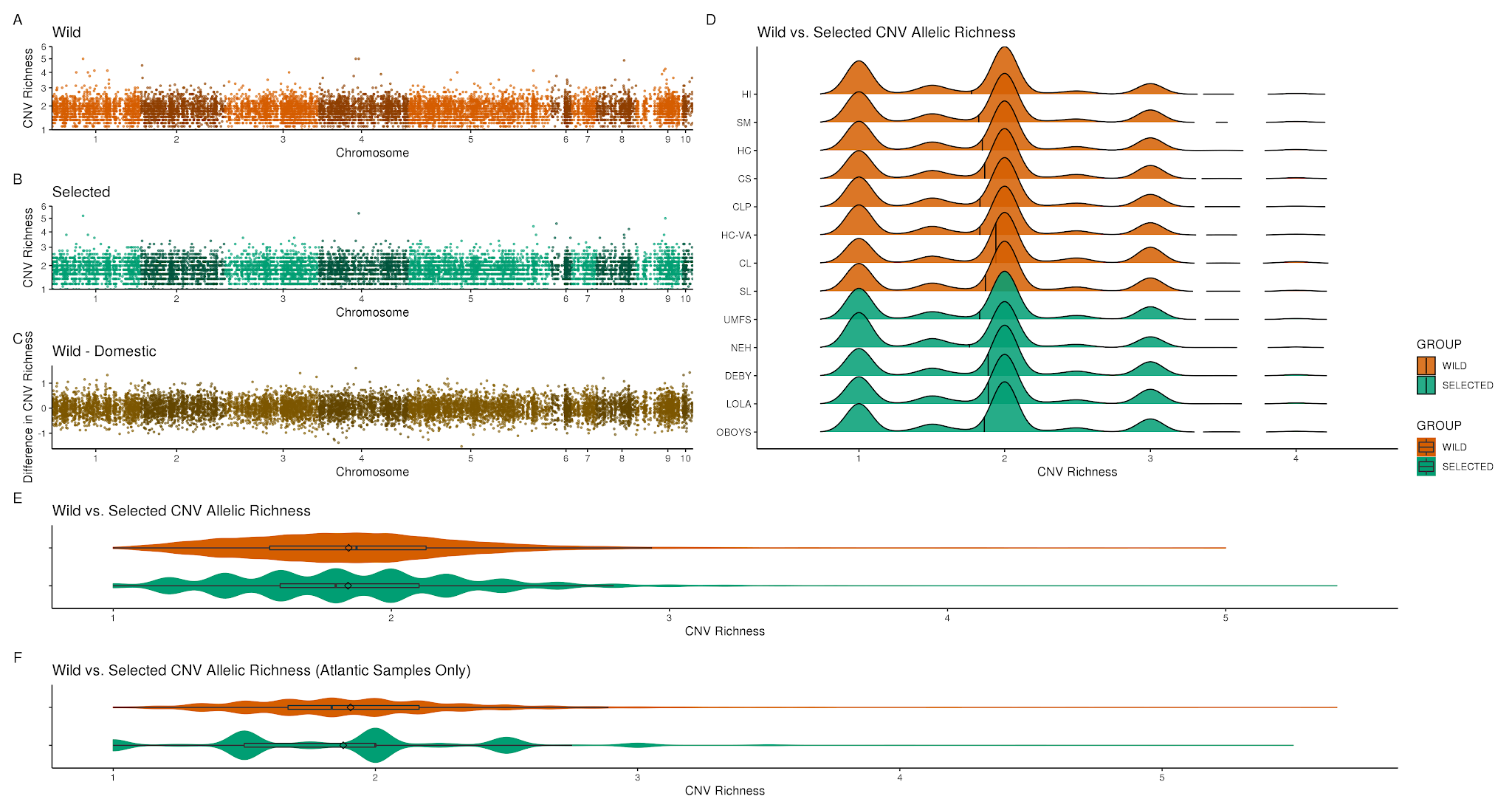
<https://raw.githubusercontent.com/The-Eastern-Oyster-Genome-Project/2021_Genome_Submissions/main/Population_Genomics/Output/Figures/Supplemental/Figure.S6.CNV.Richness.png>

**Figure S6. Comparison of copy number allelic richness across wild and selected samples.**

Panel (A) is values averaged across all wild populations in 10 kb windows across the entire genome. Panel (B) is values averaged across all selected populations in 10 kb windows across the entire genome. Panel (C) is the difference between wild and selected in 10 kb windows across the entire genome. Panel (D) is a density ridge plot of all CNVs broken down by population and line. Panel (E) is a violin and boxplot of 10kb averaged values broken by group. Panel (F) is a violin and boxplot of 10kb averaged values broken by group using only samples originating from the Atlantic.

###### Figure S7- Comparison of copy number diversity across wild and selected samples

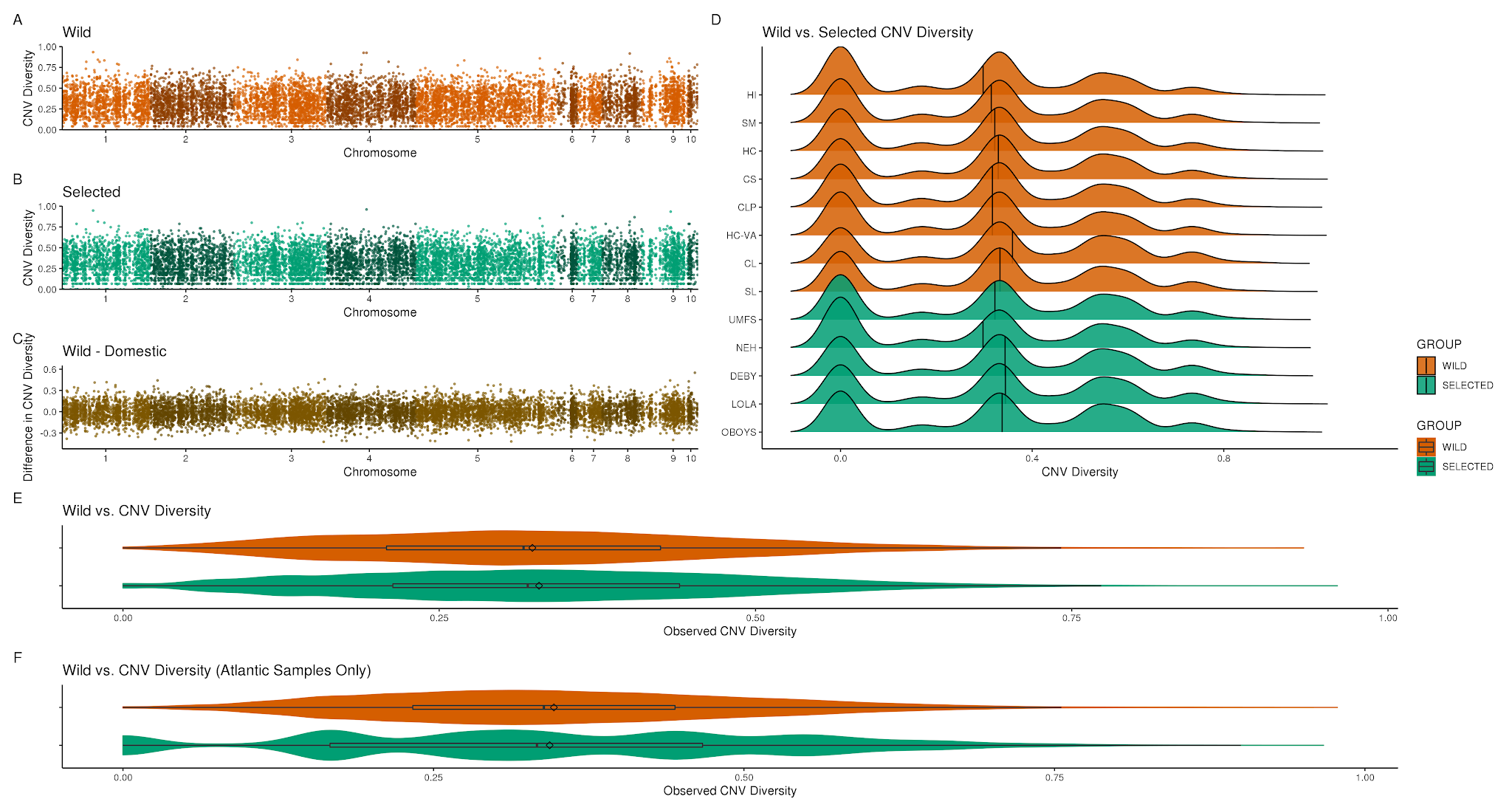
<https://github.com/The-Eastern-Oyster-Genome-Project/2021_Genome_Submissions/raw/main/Population_Genomics/Output/Figures/Supplemental/Figure.S7.CNV.Diversity.png>

**Figure S7. Comparison of copy number diversity across wild and selected samples.**

Panel (A) is values averaged across all wild populations in 10 kb windows across the entire genome. Panel (B) is values averaged across all selected populations in 10 kb windows across the entire genome. Panel (C) is the difference between wild and selected in 10 kb windows across the entire genome. Panel (D) is a density ridge plot of all CNVs broken down by population and line. Panel (E) is a violin and boxplot of 10kb averaged values broken by group. Panel (F) is a violin and boxplot of 10kb averaged values broken by group using only samples originating from the Atlantic.

###### Figure S8- Comparison of copy number allelic richness across wild and selected samples with Atlantic origins

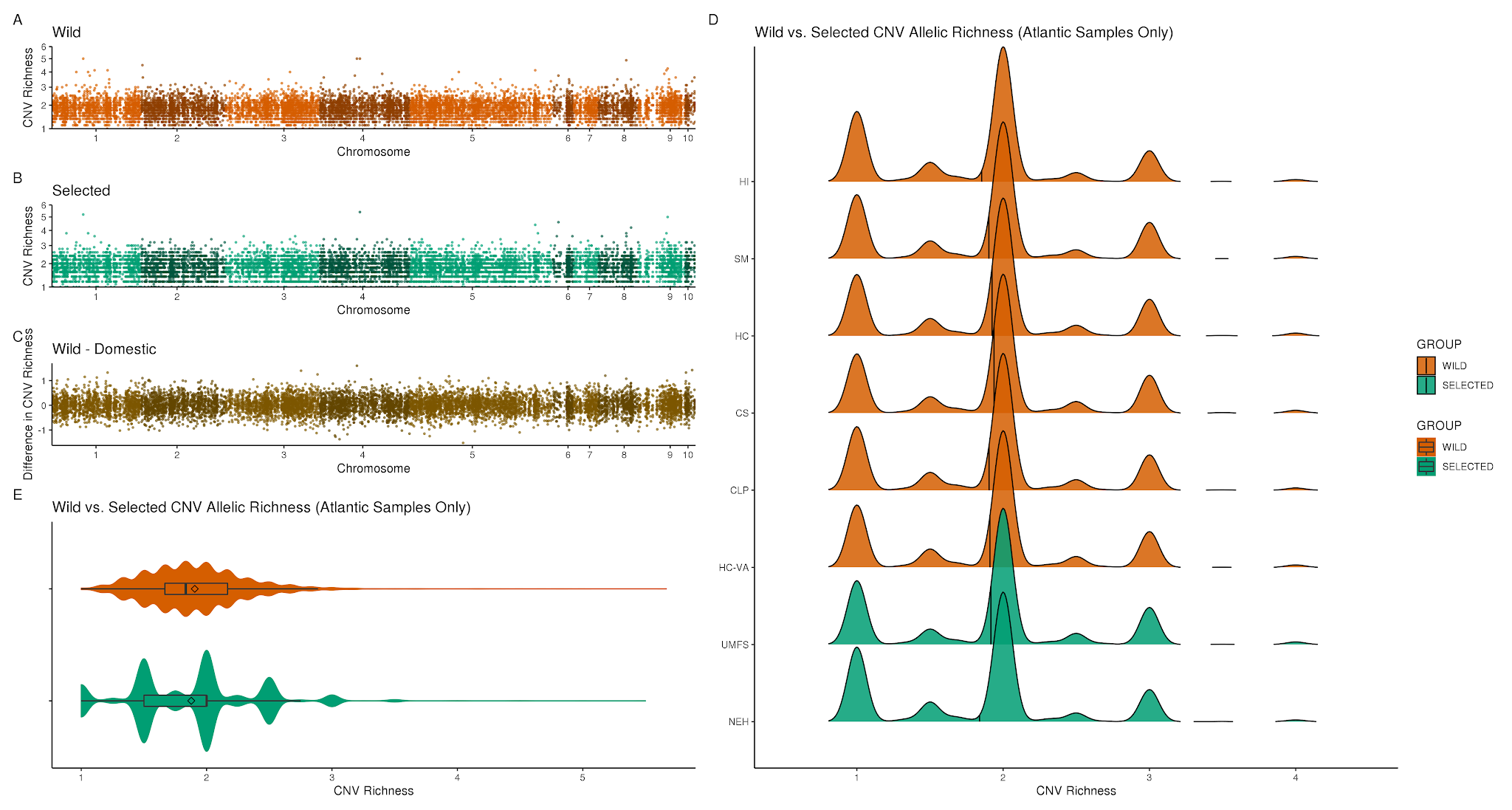
<https://raw.githubusercontent.com/The-Eastern-Oyster-Genome-Project/2021_Genome_Submissions/main/Population_Genomics/Output/Figures/Supplemental/Figure.S8.CNV.ae.Richness.png>

###### **Figure S8. Comparison of copy number allelic richness across wild and selected samples with Atlantic origins.**

Panel (A) is values averaged across all Atlantic wild populations in 10 kb windows across the entire genome. Panel (B) is values averaged across all selected lines with Atlantic origins in 10 kb windows across the entire genome. Panel (C) is the difference between wild and selected in 10 kb windows across the entire genome. Panel (D) a density ridge plot of all CNVs broken down by population and line. Panel (E) is a violin and boxplot of 10kb averaged values broken by group.

###### Figure S9- Comparison of copy number diversity across wild and selected samples with Atlantic origins

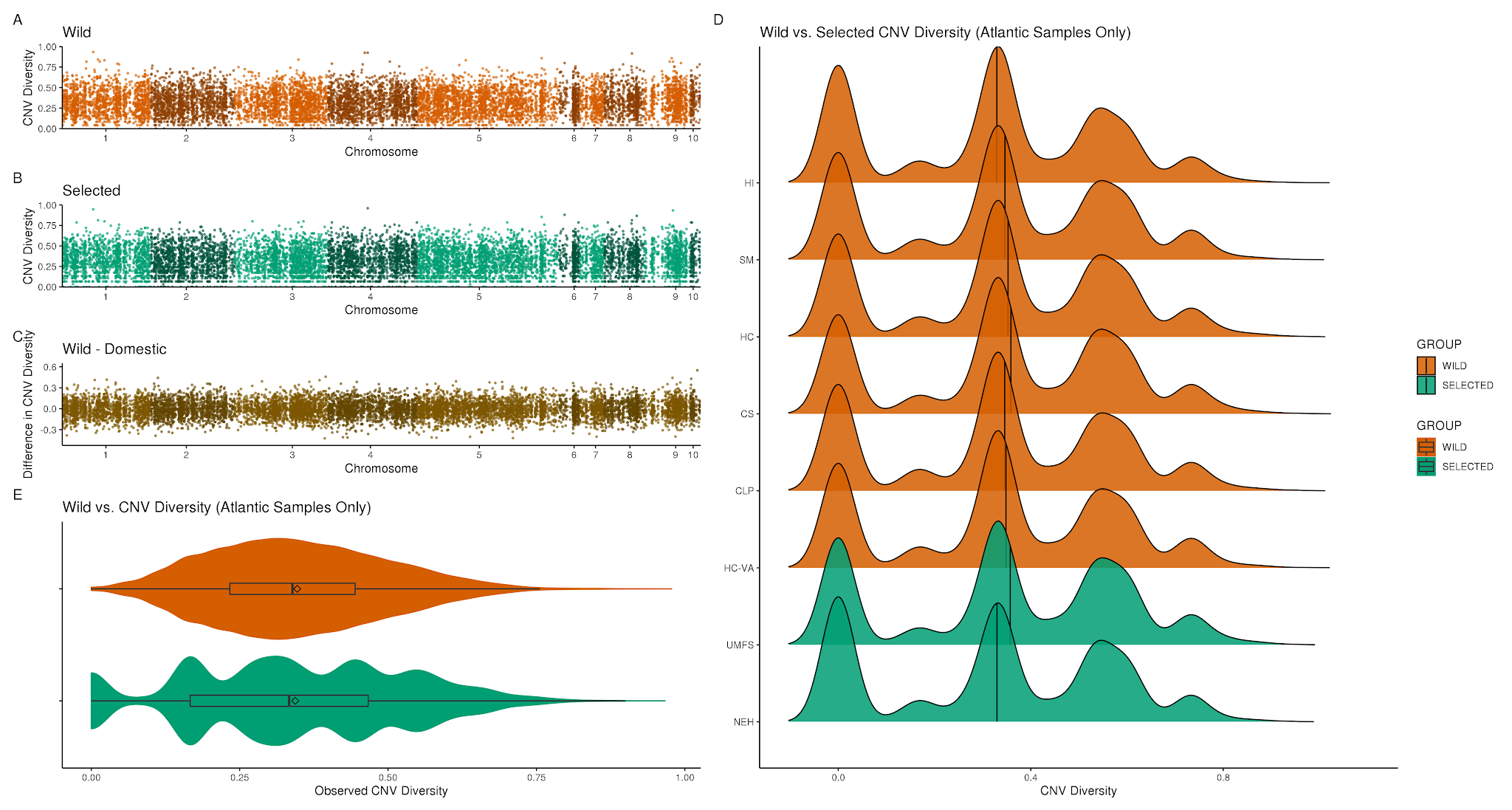

###### <https://raw.githubusercontent.com/The-Eastern-Oyster-Genome-Project/2021_Genome_Submissions/main/Population_Genomics/Output/Figures/Supplemental/Figure.S9.CNV.ae.Diversity.png>

###### **Figure S9. Comparison of copy number diversity across wild and selected samples with Atlantic origin.**

Panel (A) is values averaged across all Atlantic wild populations in 10 kb windows across the entire genome. Panel (B) is values averaged across all selected lines with Atlantic origins in 10 kb windows across the entire genome. Panel (C) is the difference between wild and selected in 10 kb windows across the entire genome. Panel (D) a density ridge plot of all CNVs broken down by population and line. Panel (E) is a violin and boxplot of 10kb averaged values broken by group.

###### Figure S10- Outlier SNPs from genomic windows with large signals of elevated *F_ST_* inside of DELY inversions

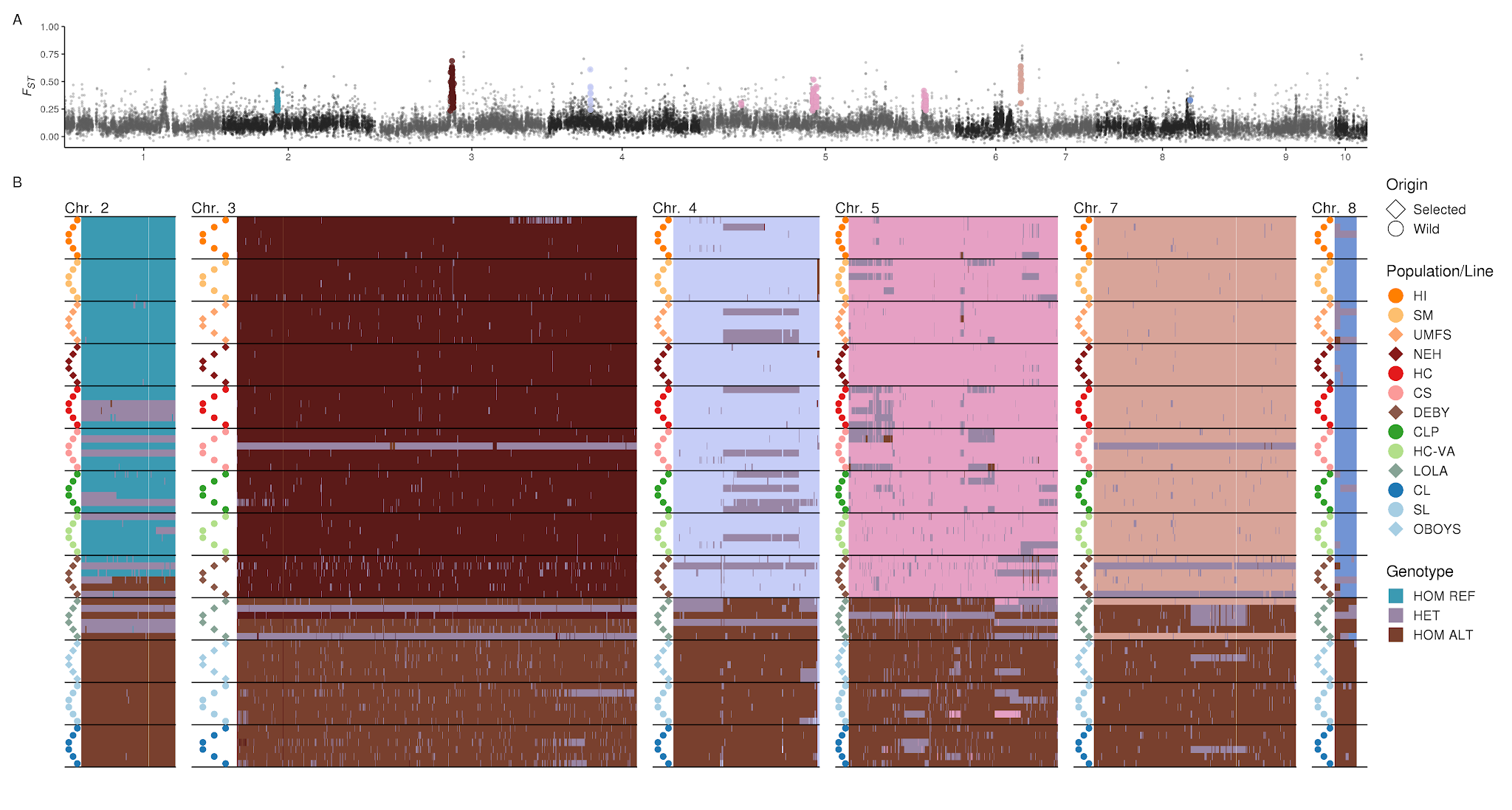
<https://raw.githubusercontent.com/The-Eastern-Oyster-Genome-Project/2021_Genome_Submissions/main/Population_Genomics/Output/Figures/Supplemental/Figure.S10.Large.FST.DI.Outlier.png>**Figure S10. Outlier SNPs from genomic windows with large signals of elevated** *F_ST_* **inside of DELLY inversions**
Outliers identified from genomic windows outside of inversion with large signals of elevated *F_ST_* . Top manhattan plot (A)- *F_ST_* values calculated by OUTFLANK and averaged across 10kb windows. Outlier SNPs are highlighted by different colors per chromosome (teal- Chr 2; maroon- Chr 3, light blue- Chr 4, pink- Chr 5, peach- Chr 7). Panel (B) contains genotype plots for outlier SNPs separated by chromosome with individuals clustered by similarity. Homozygous reference allele genotypes are colored per chromosome identically to the manhattan plot (teal- Chr 2; maroon- Chr 3, light blue- Chr 4, pink- Chr 5, peach- Chr 7)..

###### Figure S11- CNV divergence across wild population and selected lines

###### `<https://raw.githubusercontent.com/The-Eastern-Oyster-Genome-Project/2021_Genome_Submissions/main/Population_Genomics/Output/Figures/Supplemental/Figure.S11.CNV_Divergence.png>

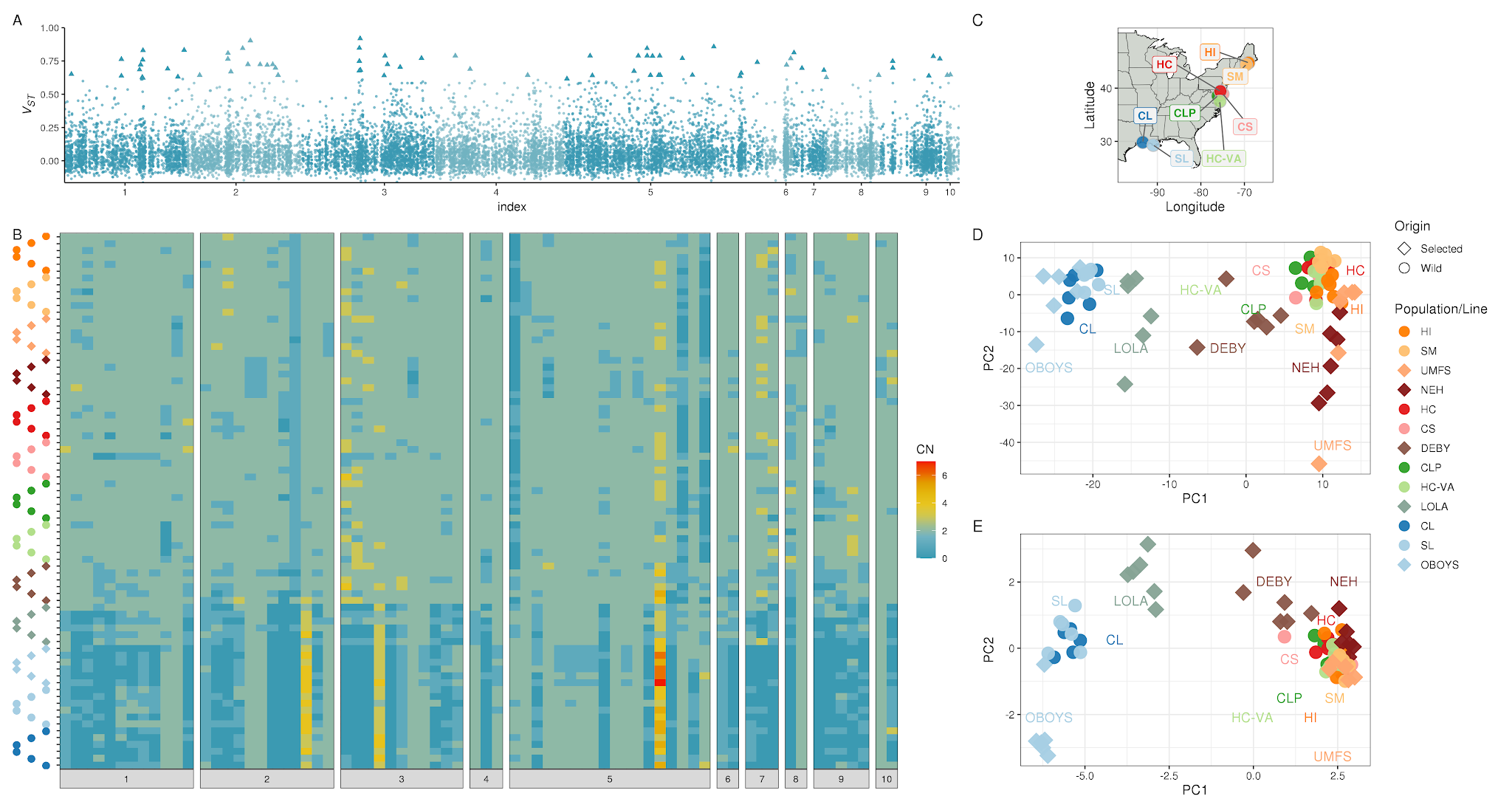

**Supplemental Figure 11. CNV divergence across wild populations and selected lines.**

Top manhattan plot (A)- *V_ST_* values for CNVs and averaged across 10kb windows. The 99.9th percentile CNVs are highlighted as triangles. Panel (B) is a heat map of the copy number per individual for the top 99.9th percentile divergent CNVs.

###### Figure S12- PCA and Genotype plots among wild atlantic individuals of outlier SNPs within large genomic inversions

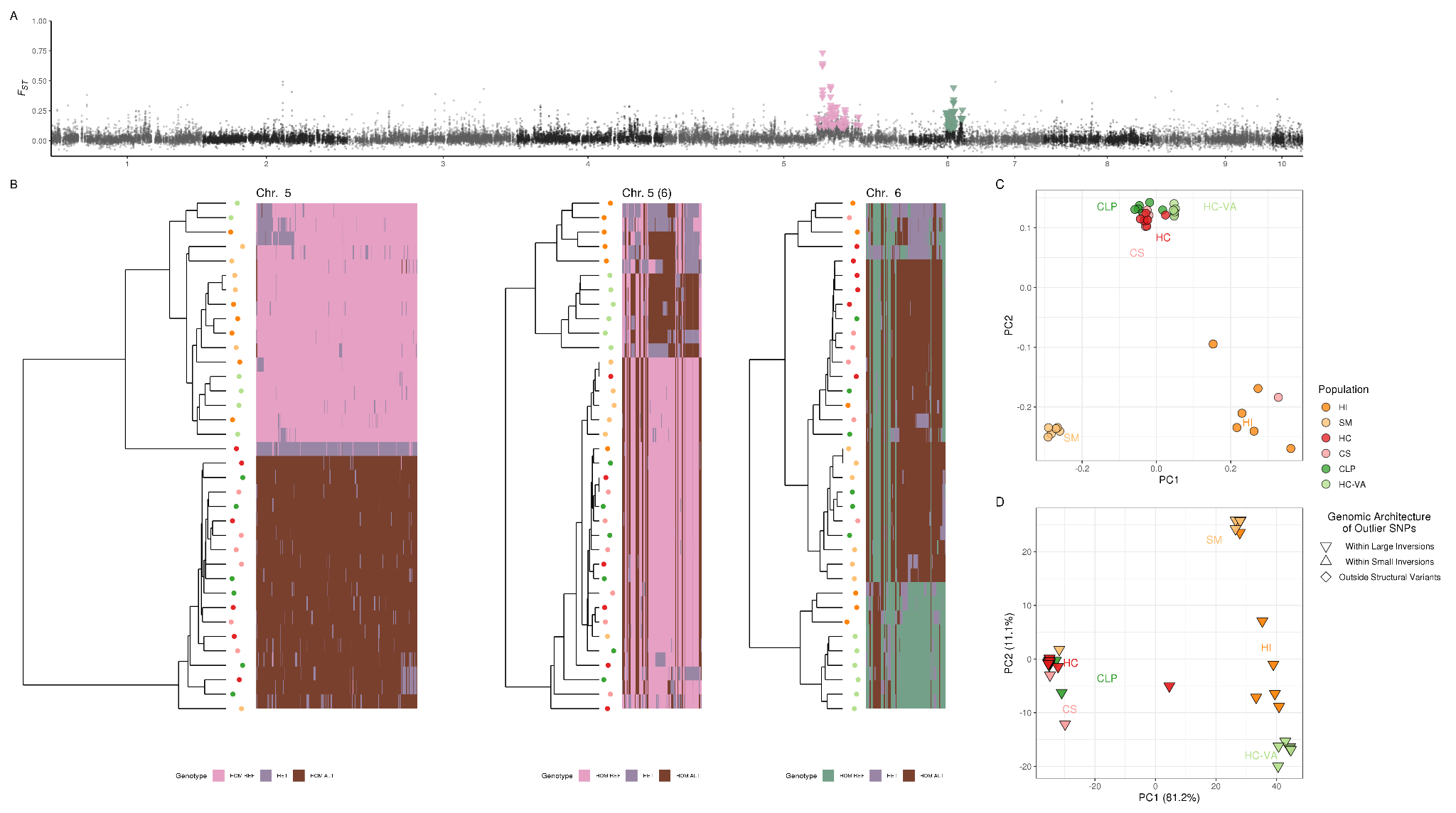

**Supplemental Figure 12. PCA and Genotype plots among wild atlantic individuals of outlier SNPs within large genomic inversions.**

Top manhattan plot (A)- *F_ST_* values calculated by OUTFLANK and averaged across 10kb windows. Outlier SNPs that were inside of large inversions are highlighted by different colors per chromosome (pink- Chr 5, green Chr 6). Panel B contains genotype plots for outlier SNPs separated by chromosome. Homozygous reference allele genotypes are colored per chromosome identically to the manhattan plot (pink- Chr 5 and green Chr 6). Panel C is a PCA analysis of all SNPs after LD clumping and outlier removal using bigsnpr (Privé et al. 2018), and Panel D is a PCA plot using only the outlier loci SNPs with colors representing the individual sampling location.

###### Figure S13- PCA and Genotype plots among wild atlantic individuals of outlier SNPs within small genomic inversions

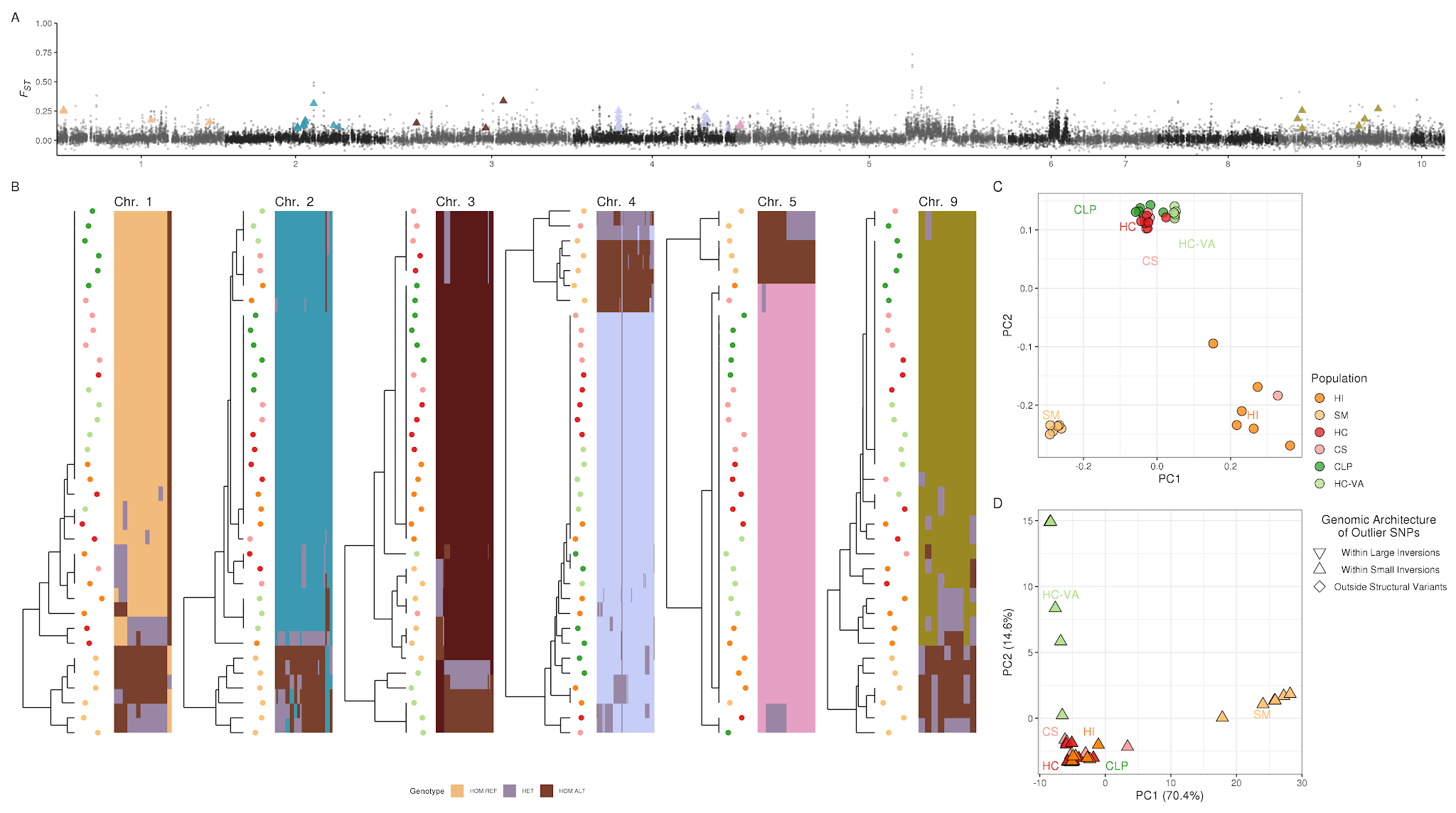

**Supplemental Figure 13. PCA and Genotype plots among wild atlantic individuals of outlier SNPs within small genomic inversions.**

Top manhattan plot (A)- *F_ST_* values calculated by OUTFLANK and averaged across 10kb windows. Outlier SNPs that were inside of small inversions are highlighted by different colors per chromosome (yellow- Chr 1; teal- Chr 2; maroon- Chr 3, light blue- Chr 4, pink- Chr 5, green Chr 6, peach- Chr 7, gold- Chr 8). Panel B contains genotype plots for outlier SNPs separated by chromosome. Homozygous reference allele genotypes are colored per chromosome identically to the manhattan plot (yellow- Chr 1; teal- Chr 2; maroon- Chr 3, light blue- Chr 4, pink- Chr 5, green Chr 6, peach- Chr 7, gold- Chr 8). Panel C is a PCA analysis of all SNPs after LD clumping and outlier removal using bigsnpr (Privé et al. 2018), and Panel D is a PCA plot using only the outlier loci SNPs with colors representing the individual sampling location.

###### Figure S14- PCA and Genotype plots among wild Atlantic individuals of outlier SNPs outside of all detected structural variants

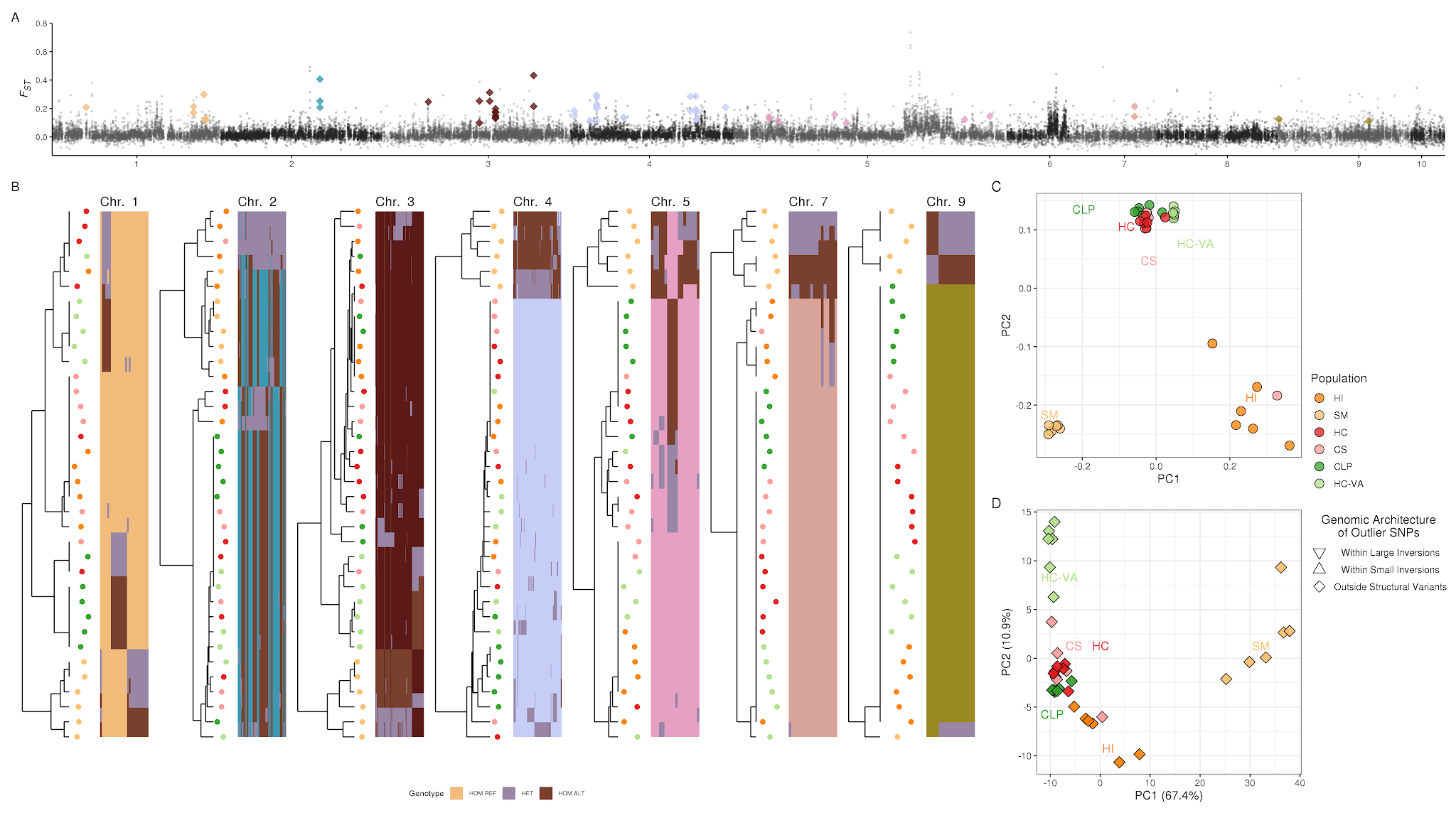

**Supplemental Figure 14. PCA and Genotype plots among wild atlantic individuals of outlier SNPs outside of all detected structural variants.**

Top manhattan plot (A)- *F_ST_* values calculated by OUTFLANK and averaged across 10kb windows. Outlier SNPs that were outside of any detected structural variants are highlighted by different colors per chromosome (yellow- Chr 1; teal- Chr 2; maroon- Chr 3, light blue- Chr 4, pink- Chr 5, green Chr 6, peach- Chr 7, gold- Chr 8). Panel B contains genotype plots for outlier SNPs separated by chromosome. Homozygous reference allele genotypes are colored per chromosome identically to the manhattan plot (yellow- Chr 1; teal- Chr 2; maroon- Chr 3, light blue- Chr 4, pink- Chr 5, green Chr 6, peach- Chr 7, gold- Chr 8). Panel C is a PCA analysis of all SNPs after LD clumping and outlier removal using bigsnpr (Privé et al. 2018), and Panel D is a PCA plot using only the outlier loci SNPs with colors representing the individual sampling location.
